# A sleep-active neuron can promote survival while sleep behavior is disturbed

**DOI:** 10.1101/2022.04.03.486914

**Authors:** Inka Busack, Henrik Bringmann

**Affiliations:** BIOTEC, Technical University Dresden, Germany

## Abstract

Sleep is controlled by neurons that induce behavioral quiescence and physiological restoration. It is not known, however, how sleep neurons link sleep behavior and survival. In *Caenorhabditis elegans*, the sleep-active RIS neuron induces sleep behavior and is required for survival of starvation and wounding. Sleep-active neurons such as RIS might hypothetically promote survival primarily by causing sleep behavior and associated conservation of energy. Alternatively, RIS might provide a survival benefit that does not depend on behavioral sleep. To probe these hypotheses, we tested how activity of the sleep-active RIS neuron in *Caenorhabditis elegans* controls sleep behavior and survival during larval starvation. To manipulate the activity of RIS, we expressed constitutively active potassium channel (*twk-18gf* and *egl-23gf*) or sodium channel (*unc-58gf*) mutant alleles in this neuron. Low levels of *unc-58gf* expression in RIS increased RIS calcium transients and sleep. High levels of *unc-58gf* expression in RIS elevated baseline calcium activity and inhibited calcium activation transients, thus locking RIS activity at a high but constant level. This manipulation caused a nearly complete loss of sleep behavior but increased survival. Long-term optogenetic activation also caused constantly elevated RIS activity and a small trend towards increased survival. Disturbing sleep by lethal blue-light stimulation also overactivated RIS, which again increased survival. FLP-11 neuropeptides were important for both, induction of sleep behavior and starvation survival, suggesting that FLP-11 might have divergent roles downstream of RIS. These results indicate that promotion of sleep behavior and survival are separable functions of RIS. These two functions may normally be coupled but can be uncoupled during conditions of strong RIS activation or when sleep behavior is impaired. Through this uncoupling, RIS can provide survival benefits under conditions when behavioral sleep is disturbed. Promoting survival in the face of impaired sleep might be a general function of sleep neurons.

**Author Summary:** The sleep neuron RIS in *Caenorhabditis elegans* is required to induce sleep behavior and to promote survival upon stress. Here we show that RIS can provide survival benefits even when sleep is disturbed.

## Introduction

Sleep is defined by behavioral inactivity and reduced responsiveness [1]. It is a physiological state that conserves and allocates energy, promotes restorative processes, and is important for brain and cognitive functions [2–4]. Sleep disorders and deviations from the normal sleep amount are widespread in modern societies and pose an unresolved medical and economical problem [5–7].

Sleep is found in most animals ranging from cnidarians to humans. Central to the induction of sleep in all species are so-called sleep-active neurons that activate to inhibit wakefulness behavioral circuits and thus cause sleep behavior [8, 9]. *C. elegans* has a small and invariant nervous system and sleep is controlled by relatively few neurons. RIS is a GABAergic and peptidergic interneuron that activates during sleep, and its ablation virtually abolishes sleep behavior during molting of developing larvae, in arrested larvae, and in adults following cellular stress [10–21]. Optogenetic manipulations showed that acute activation of RIS inhibits wakefulness activity such as feeding and locomotion [10, 15], whereas inactivation of RIS inhibits sleep [12, 18, 22, 23]. RIS is important for survival during starvation and after wounding [12, 18, 24]. This neuron counteracts aging phenotype progression during starvation-induced arrest [12]. It supports the protective transcriptional response that underlies the arrest, and promotes the activity of the FOXO transcription factor DAF-16, which is required for survival [24–26]. Thus, RIS is a sleep-active neuron that is crucial for *C. elegans* sleep and stress survival.

Little is known about how sleep-active neurons such as RIS link sleep behavior and survival. Different hypotheses are conceivable. For example, sleep-active neurons might promote survival by reducing behavioral activities during sleep, thus conserving and allocating energy or optimizing behavior [27, 28]. The close association of sleep and physiological benefits has suggested a causal relationship. For example, memory and information processing are thought to be improved in the human brain during sleep when information inputs are blocked [29, 30], and energy is conserved by reduced behavioral activity [28]. A less explored additional hypothesis is that sleep-active neurons provide benefits during sleep, but independently of sleep behavior. According to this hypothesis, sleep-active neurons have two parallel functions. The first is to induce sleep behavior, and the second is to promote survival. This hypothesis implies that physiological benefits of sleep-active neuron functions can be exerted also when sleep behavior is impaired.

In order to find out how RIS links sleep and survival, we manipulated RIS activity through transgenic expression of ion channel mutants in RIS. By strongly expressing *unc-58gf* in RIS, we achieved an overall increase of RIS activity that was caused by constantly elevated baseline calcium activity in RIS. RIS appeared to be locked at this higher activity level, as it was not able to activate further upon stimulation. Survival was increased in this strain, but sleep behavior was almost completely lacking. The sleep loss phenotype included a lack of dampening of the activity of interneurons, sensory neurons and muscles that is typical for normal sleep. Thus, strong expression of RIS led to an apparent uncoupling of sleep behavior and survival. Long-term optogenetic activation also generated constantly increased calcium activity and a small trend towards increased survival. Stimulation with blue light disturbed sleep and increased RIS activation, which in turn extended survival. Finally, we find that RIS promotes sleep behavior and promotes survival through FLP-11 neuropeptides.

RIS hence appears to be able to promote survival independently of sleep behavior, indicating that promotion of sleep behavior and survival are separable functions of RIS that are normally coupled during sleep but that can be uncoupled. Uncoupling appears to occur during conditions of stimulation that disturb sleep and increase RIS activity. This suggests that an important function of RIS overactivation is to promote survival not only during normal sleep, but also and in particular when sleep behavior is disturbed. Sleep-active neurons might generally function to promote survival both during normal sleep and even more so when sleep behavior is disturbed.

## Results

### Generation of a strain set for controlling RIS activity

We set out to control RIS activity to study sleep and survival during L1 arrest. For this study, we generated strains that express constitutively active ion channels with different ion permeability and expression levels in RIS. To constitutively inactivate RIS, we expressed potassium channels that carry mutations that cause potassium leak currents using the *flp-11* promotor that is strong and highly specific for RIS [11, 17, 31]. We used two different *egl-23* channel mutations to achieve different strengths of inactivation of RIS (*RIS::egl-23gf(weak)* expresses *egl-23(L229N)* in RIS, *RIS::egl-23gf(strong)* expresses *egl-23(A383V)* in RIS) [32]. To create a strong tool for RIS inhibition, we expressed a *twk-18* mutant allele that is known to cause a very strong potassium current (*RIS::twk-18gf* expresses *twk-18(e1913)* in RIS) [33]. In all three strains, the channel genes were knocked into the endogenous *flp-11* locus behind the *flp-11* gene to generate operons that express both *flp-11* and the mutant ion channel. All fusion proteins localized to the cell membrane of RIS (Fig 1A-D).

**Fig 1.**
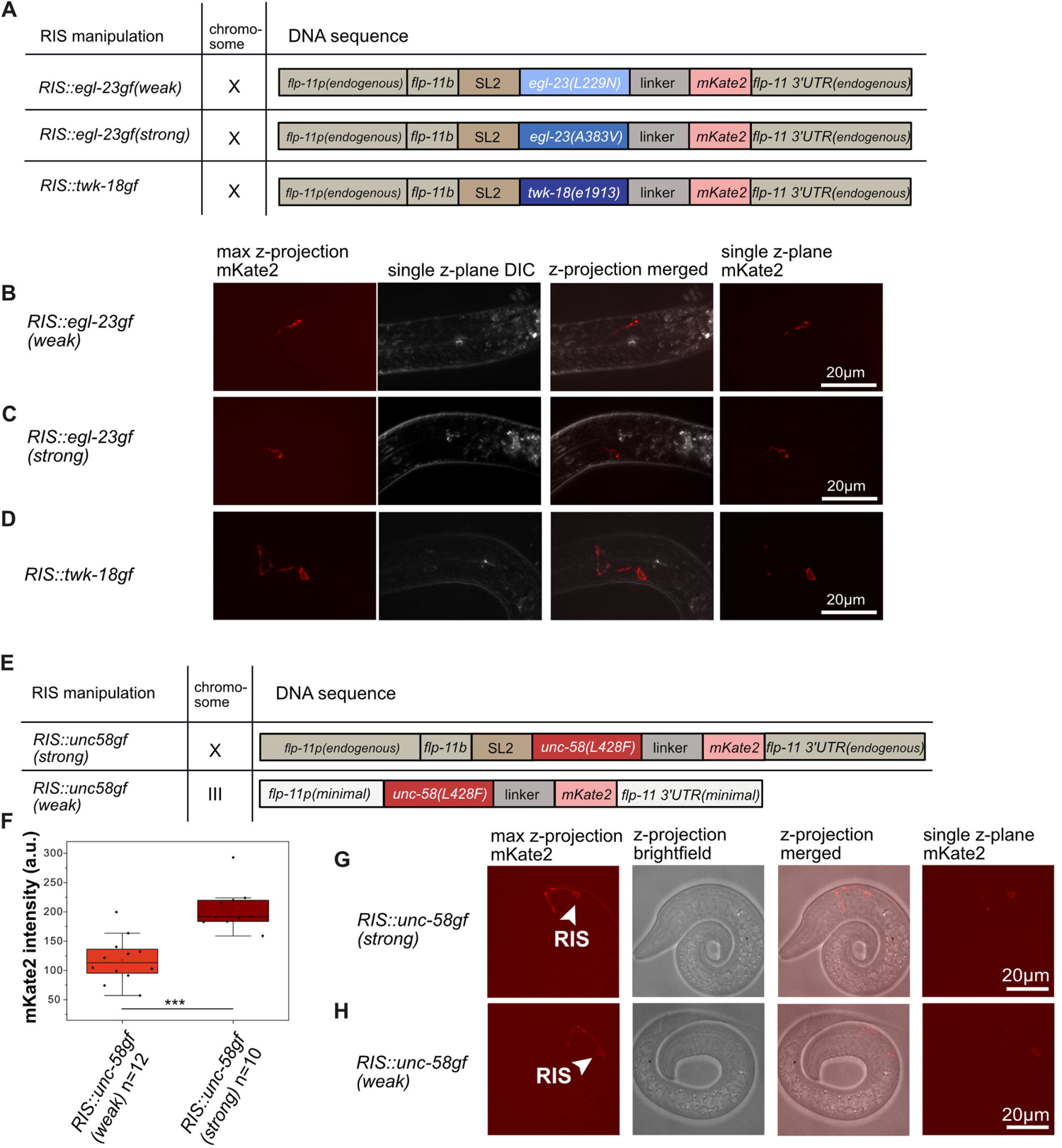
Strains expressing ion channel mutants in RIS. A) Scheme of the genetic design of the different RIS potassium channel mutant transgenes. B-D) Localization and expression of mKate2, which is translationally fused to the ion channels, in the different inactivation strains. E) Schematic representation of the genetic design of the RIS sodium channel mutant transgenes. F) Quantification of the expression levels of UNC-58gf::mKate2 in the two RIS overactivation strains. ***p<0.001, Welch test. G) Localization and expression of the strong overactivation strain (*RIS::unc-58gf(strong)*). H) Localization and expression of the weak overactivation strain (*RIS::unc-58gf(weak)*).

To constitutively over activate RIS, we used an unusual K2P channel mutant that is permeable to sodium [32] (Thomas Boulin, personal communication). We expressed the ion channel mutant in RIS (*unc-58(L428F),* Fig S1A) tagged with mKate2 at either a lower or a higher level in RIS. The different expression levels were achieved by generating two different expression loci for the channel in the genome. To achieve a strong expression of the *unc-58(L428F)* channel (*RIS::unc-58gf(strong)*), we turned the endogenous *flp-11* locus into an operon as above. To achieve a weaker expression of the channel (*RIS::unc-58gf(weak)*), we expressed mKate2-tagged *unc-58(L428F)* from a single-copy knock-in locus on chromosome III [34], which contained only a minimal *flp-11* promoter and 3’ UTR [11] (Fig 1 E-H). Fluorescent imaging after two days of L1 arrest showed that the stronger strain expressed at about double the level of the weaker strain and in both transgenes, the channel was expressed prominently in RIS and was targeted to the cell membrane (Fig 1E-H, S1). Even after 21 days of arrest, mKate2-tagged *unc-58(L428F)* expression could be detected in the strong strain indicating that the sodium channel mutant is properly folded and targeted to the plasma membrane across the entire arrest (FigS1E).

### *RIS::unc-58gf(strong)* promotes survival of L1 arrest

Previous studies have shown that RIS is required for survival during larval arrest and following wounding of the adult [12, 18, 35]. RIS activity controls the gene expression changes that underly the physiological adaptation to starvation [24]. *RIS::unc-58gf(strong)* and *RIS::twk-18gf* displayed opposing gene expression changes compared with the wild type. *RIS::twk-18gf* is more similar to *aptf-1(-)*, which causes loss of RIS function [10, 24]. To identify additional genes whose expression depend on RIS activity, we directly compared the gene expression profiles of *RIS::unc-58gf(strong)* and *RIS::twk-18gf* during L1. This analysis identified 4676 differentially expressed genes including 1910 novel genes that were not previously reported to be controlled by RIS (Fig 2A, Table S5). This comparison confirms the notion that activation and inactivation of RIS have opposing effects on gene expression [24]. These data support the idea that the expression of *twk-18gf* and *unc-58gf* in RIS cause RIS inactivation and RIS activation gene expression phenotypes, respectively [24].

**Fig 2.**
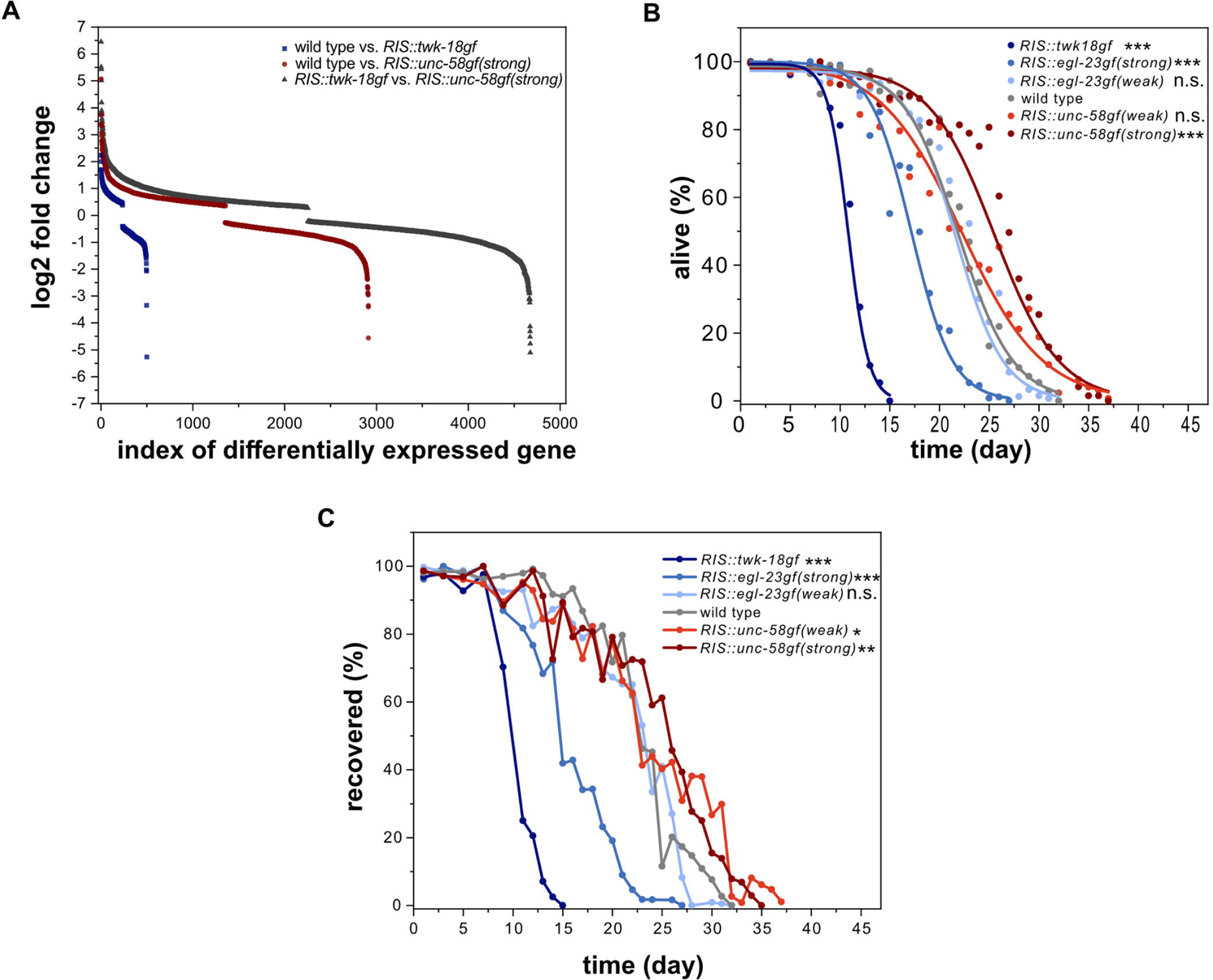
Survival of starvation-induced L1 arrest is a function of RIS activity A) *RIS::twk-18gf* and *RIS::unc-58gf(strong)* strains cause divergent transcriptional profiles. Shown is the log2 fold change for all differentially expressed genes. A gene was counted as differentially expressed with an FDR<0.05. According to our previously published results, *RIS::twk-18gf* expressed 498 genes differentially compared with the wild type. *RIS::twk-18gf* has been shown to be similar to *aptf-1(gk794)* [24]*. RIS::unc-58gf(strong)* was shown to express 2913 genes differentially compared with the wild type [24]. Comparing *RIS::twk-18gf* and *RIS::unc-58gf(strong)* revealed 4676 differentially expressed genes. This analysis supports the view that these two strains have opposing transcriptional profiles and identifies 1910 additional genes that are potentially under the control of RIS activity (Table S5). B) Strong inactivation significantly decreases survival in L1 arrest whereas strong activation prolongs survival (three-parameter logistic fit, see also supplementary table S1, Fisher’s Exact Test was conducted when 50% of wild-type worms were alive (day 22)). The p-values were FDR corrected with Benjamini-Hochberg procedure with a 5% false discovery rate. The plot includes data from three replicates. ***p<0.001. C) Recovery from L1 arrest depends on RIS activity with strong inactivation leading to a reduced recovery and constant activation leading to a prolonged ability to recover. For statistics, the Fisher’s Exact Test was conducted at the time when 50% of wild-type worms recovered (day 23). The p-values were FDR corrected with Benjamini-Hochberg procedure with a 5% false discovery rate. The plot includes data from two replicates. N.s. p>0.05, *p<0.05, **p<0.01, ***p<0.001. Fisher’s Exact Test.

**Table 1.** Calculated functions for the regressions.

| Figure panel | fit | function |
| --- | --- | --- |
| 6C | linear | $y = 0.116 + 0.5494x$ |
| 6G | linear | $y = 0.21203 + 1.10949x$ |

To get an understanding of the role of RIS activation on survival, we investigated how the different RIS-expressed ion channels affect survival in arrested L1 larval worms. For this experiment, we kept arrested L1 larvae as liquid culture, as this allowed for a higher number of animals to be processed. We regularly took a sample of the larvae from the liquid culture and measured two different readouts. We first measured starvation survival by scoring the fraction of animals that had survived and secondly, we scored the ability of starved animals to recover and develop to the adult stage after refeeding [12, 23, 24, 36]. *RIS::twk-18gf* larvae showed strongly reduced starvation survival by about half compared to the wild type. *egl-23(strong)* resulted in a more moderate yet significant starvation survival reduction of approximately 21%. Weak inactivation by *egl-23(weak)* or overactivation by *unc-58gf(weak)* of RIS activity did not significantly change starvation survival. Strong RIS overactivation by *unc-58gf(strong)* had an increased starvation survival of around 19% compared to wild type (Fig 2B). These results are in accordance with previous reports and the expected changes caused by the ion channel transgenes [12, 24, 32, 33]. The recovery upon refeeding matched the survival curves (Fig 2C). To test whether this survival-extending effect was caused by *unc-58gf* expression in RIS, we measured survival of *RIS::unc-58gf(strong)* in an *aptf-1(-)* background, in which expression of *unc-58gf* in RIS is abolished (Fig S1G). In the combined *RIS::unc-58gf(strong); aptf-1(-)* strain starvation survival was extremely shortened compared with wild type (Fig S2) and was similar to *aptf-1(-)* [12] or *RIS::twk-18gf* (Fig 2B), indicating that the starvation survival extension indeed stemmed from *unc-58gf* expression in RIS.

### Ion channel mutant expression in RIS controls baseline RIS activity

How are RIS activity, sleep behavior and survival linked? To shed light on this question, we imaged RIS calcium activity in these mutants. [10]. We used microfluidic chambers to keep and image the larvae. This method allowed for a quantification of RIS baseline activity, RIS calcium activation transients, and sleep behavior [37, 38]. We were able to measure RIS activity for the first 12 days of starvation, and found that the effects of the channel transgenes were consistent across the entire time of starvation that we analyzed (Fig 3A for two-day starvation, 3B for all time points). We first analyzed the baseline activity of RIS in the absence of calcium transients. The mean RIS baseline calcium activity levels were reduced by the strongly inactivating transgenes *twk-18gf* and *egl-23gf(strong)*. Baseline activity levels were modestly increased by *unc-58gf(weak)* and strongly increased by *unc-58gf(strong)*. The magnitude of the baseline calcium signal alterations of the different transgenes were thus in accordance with the expression levels and the expected strengths of the leak currents (Fig 1F). These data demonstrate that transgenic *unc-58gf* can be used as a tool for increasing calcium activity in a neuron that does not normally express *unc-58* [17, 31, 32]. We used these tools to create a set of strains that allows for probing associations of baseline activity levels and survival as well as sleep.

**Fig 3.**
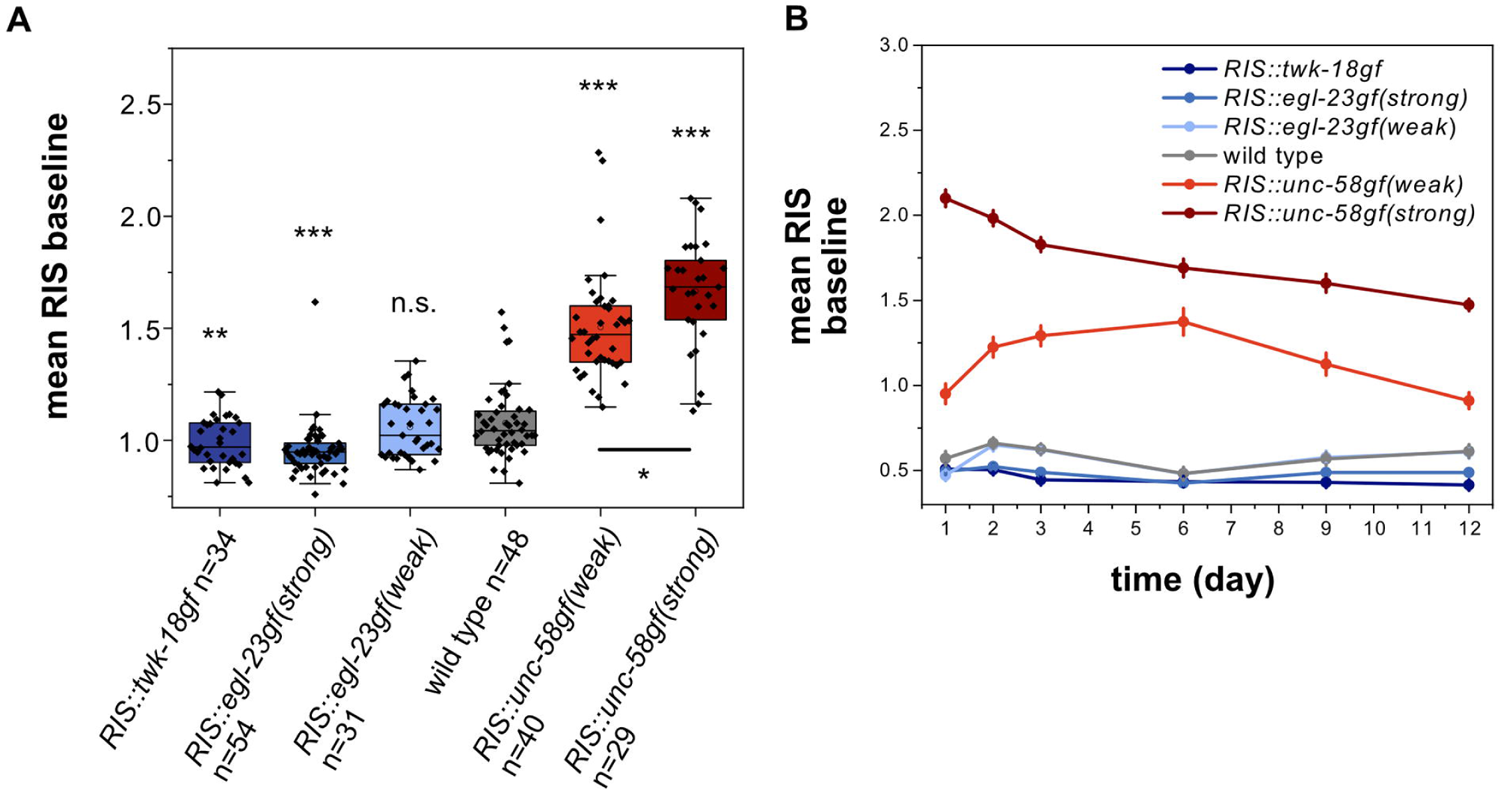
Ion channel mutant expression in RIS controls baseline RIS activity A) Ion channel expression in RIS results in a dose-response strain set with varying levels of RIS baseline activity. n.s. p>0.05, **p<0.001, ***p<0.001, Welch test with FDR correction for multiple testing. B) Long term RIS baseline activity for RIS activity strains from day 1 to day 12 of starvation.

### *RIS::unc-58gf(strong)* inhibits sleep behavior during L1 arrest

We next quantified motion quiescence as a readout for sleep and tracked mobility behavior during L1 arrest inside microfluidic chambers. We used GCaMP fluorescence to quantify RIS activation transients across the entire measurement [10]. Consistent with previous reports [10, 12, 20], wild-type RIS showed activation transients over a constant baseline, and calcium transients were accompanied by phases of behavioral quiescence (Fig 4A-G). Calcium transients were completely abolished in both the strongest deactivation strains (Fig 4A, B) as well as in the strong activation strain (4F). In the mild deactivation strain calcium transients were still visible and correlated with behavioral quiescence, but were decreased in magnitude (Fig 4C/G, S3A). In *RIS::unc-58gf(weak)*, RIS calcium transient activity and bouts of behavioral activity were increased (Fig 4E). These results suggest that RIS activation transients are required for sleep induction and that strong inactivation prevents these transients. Mobility quiescence decreased with inactivation and was increased with modest activation as expected. We anticipated that the elevated RIS calcium baseline of *RIS::unc-58gf(strong)* might be associated with a further increase in sleep behavior compared with the weak activation. Mobility quiescence was, however, completely abolished by strong RIS activation in most of the animals (Fig 4H). These findings were additionally confirmed for a longer period of starvation up to day 12 (Fig 4I). The loss of quiescence of the strong RIS activation strain was similar to, but not as strong as in *aptf-1(-)* (Fig S2B). The sleep amounts for each strain are consistent for several days in L1 arrest (Fig 4I). In summary, baseline RIS activity correlates with behavioral quiescence over a wide range of conditions, except for the *RIS::unc-58gf(strong)* transgene, which appears to cause a lack of RIS calcium transients and sleep behavior.

**Fig 4.**
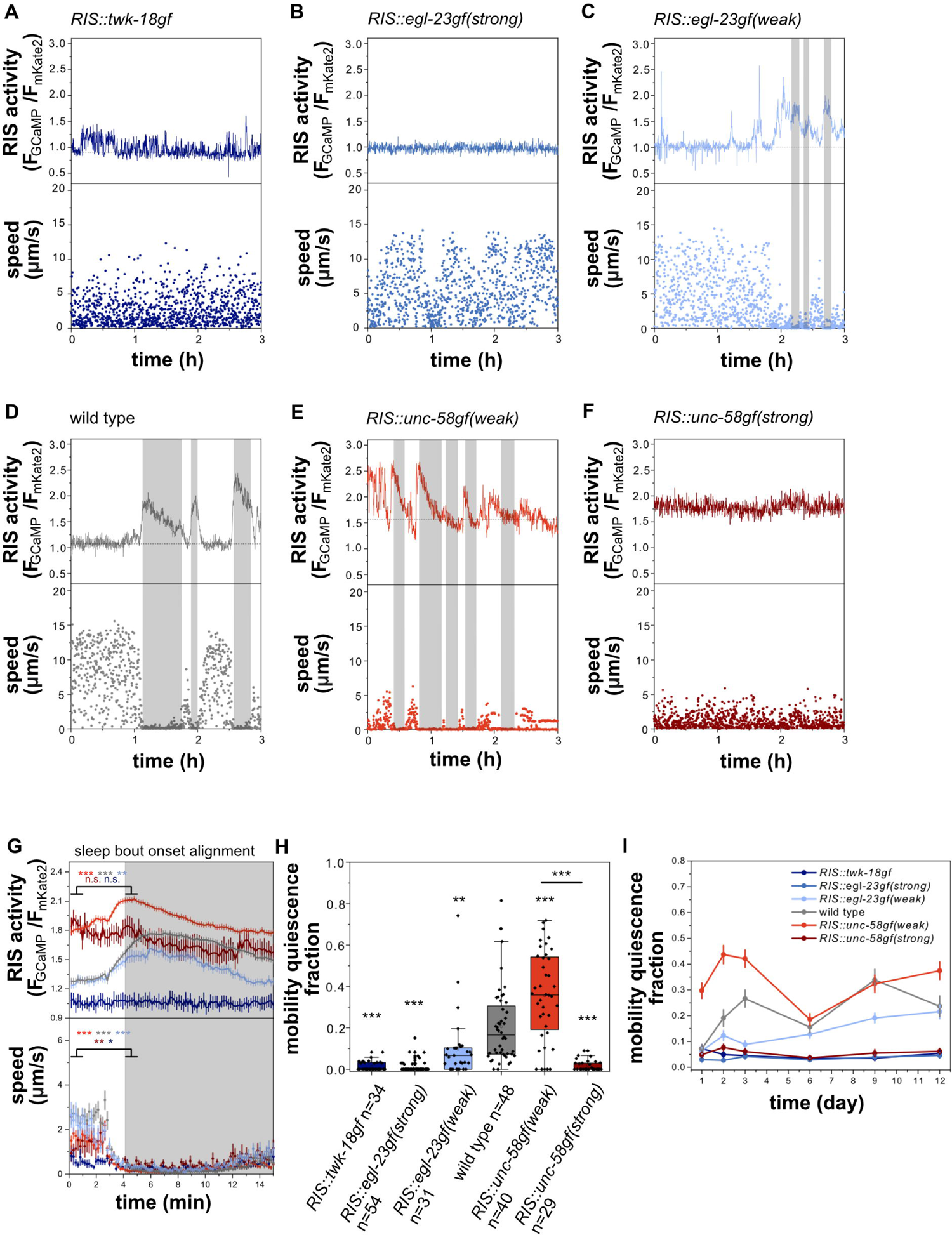
Both, strong inactivation of RIS as well as *RIS::unc-58gf(strong)* prevent sleep behavior during L1 arrest A-F) Ion channel expression in RIS controls calcium activity and sleep behavior. Mobility quiescence bouts (grey shade) correlate with RIS transients. Movement speed was calculated by tracking the position of the head neuron RIS. A) *RIS::twk-18gf* and B) *RIS::egl-23gf(strong)* larvae do not have detectable mobility quiescence bouts or RIS transients. C) *RIS::egl-23gf(weak)* has reduced mobility quiescence bouts. D) Wild-type RIS activation transients and immobility bouts. E) *RIS::unc-58gf(weak)* spends more time in mobility quiescence bouts. F) *RIS::unc-58gf(strong)* has no detectable mobility quiescence bouts and no RIS transients but a high constant activity. G) RIS activation and immobility are correlated only for the strains that also show sleeping behavior. Alignment of RIS activity and speed to mobility quiescence bout onset. **p<0.01, ***p<0.001, Wilcoxon signed rank test. H) Mobility quiescence is a function of RIS activity levels n.s. p>0.05, **p<0.001, ***p<0.001, Welch test with FDR correction for multiple testing. I) Long-term sleep fraction for RIS activity strains from day 1 to day 12 of starvation.

### *RIS::unc-58gf* impairs sleep behavior during larval development and following cellular stress in the adult

*C. elegans* sleeps during various stages and conditions. Previous studies showed that cellular stress increases adult sleep [17, 39–42]. During development, sleep is coupled to the molting cycle and occurs during a period called lethargus, which coincides with formation of a new cuticle [10, 43, 44]. To test whether constant RIS baseline activation is also associated with a loss of behavioral quiescence during stress-induced sleep and during lethargus sleep, we tested whether *RIS::unc-58gf* affects sleep behavior during L1 development as well as following heat stress. For testing for stress-induced sleep, we kept young adult worms inside microfluidic devices and first measured baseline behavioral activity at 22°C, and then exposed the animals to heat stress at 37°C for 20 minutes. We then followed the behavior again after the end of the heat shock [17]. The wild type immobilized during the heat shock and displayed an increase in behavioral quiescence following the heat shock. The quiescence response was attenuated in all strains in which RIS was manipulated. Both *unc-58gf* expressing strains showed decreased behavioral quiescence. Thus, a reduction of behavioral sleep can also be caused by *RIS::unc-58gf(weak)* (Fig 5A-G).

**Fig 5.**
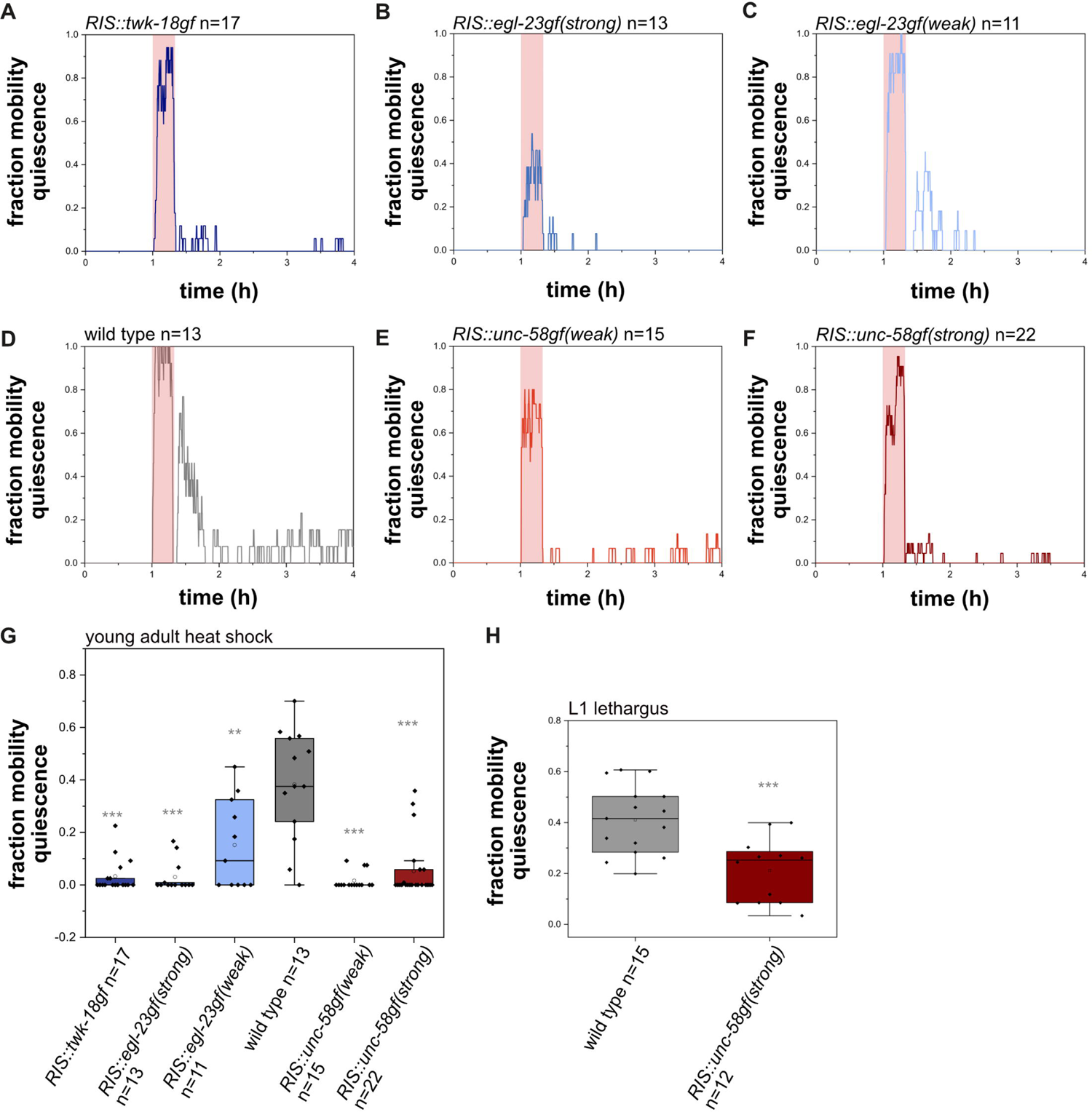
*RIS::unc-58gf* impairs sleep following cellular stress and during the L1 molt in developing larvae A-F) Sleep fraction of worms of different RIS activity strains during the heat shock experiment. Animals were recorded continuously throughout the experiment. First, a 1h baseline was recorded at 22°C, then a 20min heat shock period of 37°C (in red shade) was applied, after the heat shock, the temperature was lowered to at 22°C again to monitor the response to the heat shock. G) Quantification of time spent in sleep 20 min after the heat shock. Wild-type worms sleep on average 38.3%. All other RIS manipulations reduced quiescence. *RIS::unc-58gf(strong)* only sleeps on average 6.5% of the time. **p<0.001, ***p<0.001, Welch test with FDR correction for multiple testing. H) *RIS::unc-58gf(strong)* reduces lethargus sleep compared to wild type. ***p<0.001, Welch test.

To test for sleep that occurs during development, we quantified behavioral quiescence of *RIS::unc-58gf(strong)* during lethargus in fed L1 larvae. Behavioral quiescence of *RIS::unc-58gf(strong)* was significantly reduced compared with the wild type. The loss of sleep behavior in lethargus was, however, not as strong as during L1 arrest or following stress-induced sleep (Fig 5H). Hence, in all the stages and conditions that we tested, sleep behavior was reduced by *RIS::unc-58gf(strong)*.

We also tested whether the lack of sleep behavior in *RIS::unc-58gf(strong)* during L1 arrest causes a compensatory increase in sleep when the animals resume development. For this experiment, we starved larvae for 12 days and then provided them with food, which allowed for a resumption of development. We then imaged sleep behavior during L1 lethargus in these recovered worms. There was no increase of sleep behavior in recovered *RIS::unc-58gf(strong)* larvae compared with wild type (Fig S4). Thus there was no indication for rebound sleep occurring at the subsequent lethargus stage in *RIS::unc-58gf(strong)*.

### *RIS::unc-58gf(strong)* impairs the dampening of wake-active neuronal circuits during L1 arrest

*RIS::unc-58gf(strong)* caused strongly reduced sleep behavior, but the nature of the residual sleep behavior in this mutant was unclear. Did the residual sleep behavior present a generally weakened sleep state or a compensatory deep sleep state? To answer this question, we characterized neuronal activity during in *RIS::unc-58gf(strong)* mutants. For this set of experiments, we imaged larvae in microfluidic chambers and imaged neuronal activity with GCaMP during L1 starvation-induced arrest.

*C. elegans* sleep is characterized as a state of globally reduced nervous system activity [12, 13, 20]. We measured the overall neuronal activity by utilizing a strain that expresses GCaMP6s in all neurons [45]. We measured both motion behavior and overall neuronal activity by extracting the calcium activity signal of all neurons as one combined signal [12, 22]. Consistent with previous reports, periods of behavioral quiescence were accompanied by strongly reduced global neuronal activity (Fig 6A, B, S5) [12, 13, 20, 22]. We extracted phases of neuronal inactivity as well as phases of behavioral inactivity from the data set. In all strains, neuronal inactivity directly correlated with immobility, suggesting that behavioral quiescence and neuronal quiescence are correlated in all strains irrespective of the level of RIS baseline activity (Fig 6C). Thus, both behavioral inactivity and brain inactivity can serve as equivalent readouts for detecting sleep in this system. Strong RIS inactivation led to an increased mean overall neuronal activity of 50.7% compared to wild type. Intermediate strength of inactivation did not affect mean overall neuronal activity. *RIS::unc-58gf(strong)* caused a significant increase of mean overall neuronal activity of 35.7% (Fig 6D). These results support the notion that *RIS::unc-58gf(strong)* worms are in a state of wakefulness for most of the time and lack phases of strong nervous system activity dampening.

**Fig 6.**
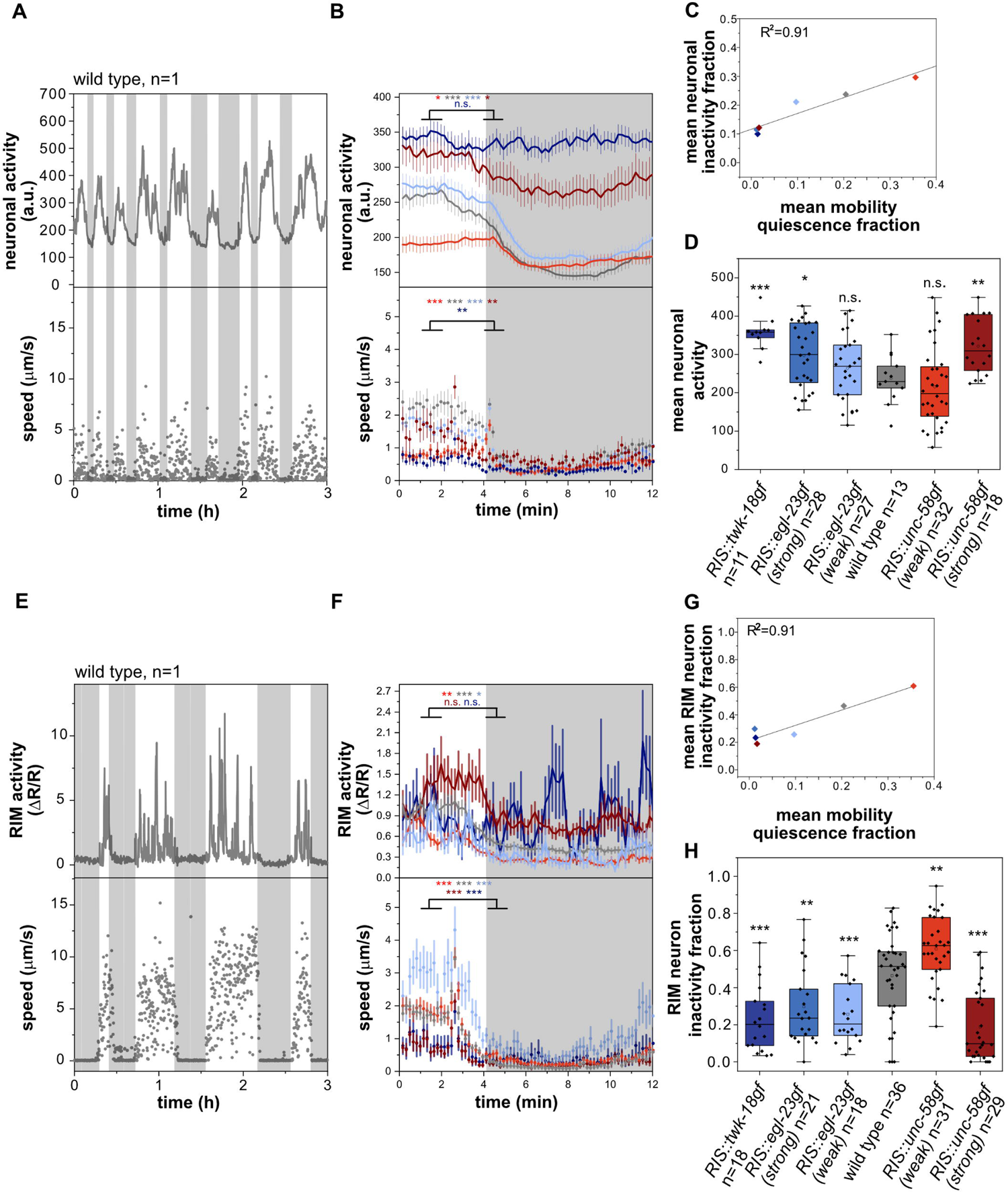
*RIS::unc-58gf(strong)* fails to inhibit wake-active circuits A-B) Reduced global neuronal activity (grey shading) correlates with motion quiescence (supplementary movies S1-S6). Head speed was measured. A) sample trace, B) alignment of overall neuronal activity to mobility quiescence bout onset. ***p<0.001, Wilcoxon signed rank test. C) The mean neuronal inactivity fraction correlates with the mean mobility quiescence fraction (linear fit) for all strains. D) Strong RIS inactivation as well as *RIS::unc-58gf(strong)* caused an overall increase of neuronal activity. n.s. p > 0.05, *p<0.05, **p<0.01, ***p < 0.001, Welch test with FDR correction for multiple testing. E-F) RIM does not activate during mobility quiescence bouts. RIM speed was measured F) Sample trace (grey shade is RIM inactivity bout). Alignment of RIM activity and speed to mobility quiescence bout onset. *p<0.05, **p<0.01, ***p<0.001, Wilcoxon signed rank test. G) The mean RIM inactivity fraction correlates with the mean mobility quiescence fraction (linear fit) for all strains. H) Strong RIS inactivation as well as *RIS::unc-58gf(strong)* caused an overall increase of RIM activity. **p < 0.01, ***p < 0.001, Welch test with FDR correction for multiple testing.

To characterize the effects of *RIS::unc-58gf(strong)* at the level of specific individual neurons, we imaged activity of the RIM neurons, which are second layer interneurons that integrate information from sensory and internal states to control motion behaviors [46, 47]. Again, we used GCaMP imaging in microfluidic chambers to correlate neuronal and behavioral activity. RIM showed activation transients during phases of behavioral activity. During phases of behavioral quiescence, RIM activity was strongly reduced and activation transients were lacking (Fig 6E, F, S6). Similar to the pan-neuronal measurements, the time fraction during which RIM was inactive correlated with phases of motion quiescence (Fig 6G). Consistent with the pan-neuronal imaging data, *RIS::unc-58gf(strong)* lacked phases of RIM inactivity (Fig 6H).

An increased arousal threshold and attenuated sensory information processing are key characteristics of sleep [1, 48]. In *C. elegans*, sensory information processing is inhibited during sleep in developing larvae and the activity of sensory neurons is reduced [49–51]. For example, the mechanosensory neurons ALM and PLM display dampened activation transients during sleep in developing L1 larvae [49, 50]. We measured PLM activity in wild-type, *RIS::unc-58gf(strong)* and *RIS::twk-18gf* larvae. In wild-type animals, PLM showed activation transients only during phases of wakefulness, whereas PLM appeared to be rather inactive during sleep. In both *RIS::unc-58gf(strong)* and *RIS::twk-18gf* mutants, PLM showed activation transients almost continuously throughout the time, with only small episodes where PLM seemed to be not active (Fig 7). Thus, extended periods of PLM neuron inactivity were absent in both *RIS::unc-58gf(strong)* and *RIS::twk-18gf* mutants. Thus, *RIS::unc-58gf(strong)* inhibits the dampening of neuronal activity that is associated with wakefulness behavior and thus lacks the neuronal signature of deep sleep.

**Fig 7.**
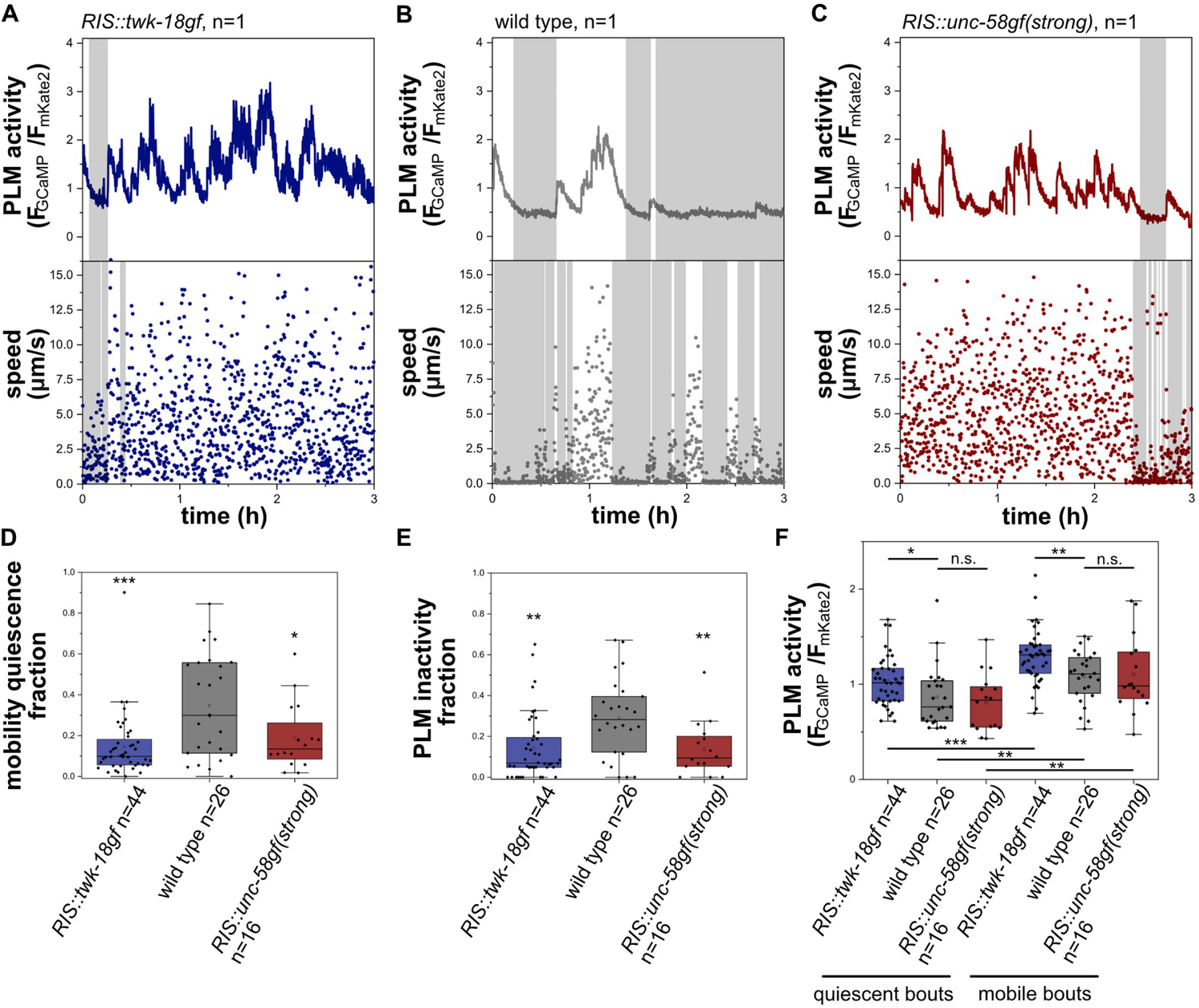
Phases of sensory PLM neuron inactivity occur during sleep and are strongly reduced in *RIS::unc-58gf(strong)* and *RIS::twk-18gf* A-C) Sample traces of PLM activity and speed in wild type, *RIS::unc-58gf(strong)* and *RIS::twk-18gf*. PLM speed was measured. D) *RIS::twk-18gf* and *RIS::unc-58gf(strong)* reduced phases of sensory neuron quiescence. *p<0.05, *** p<0.001, Welch test. E) *RIS::twk-18gf* and *RIS::unc-58gf(strong)* caused an overall increase of PLM activity. **p<0.01, ***p < 0.001, Welch test. F) *RIS::twk-18gf* causes increased PLM activity in detected quiescence as well as mobility bouts. *p<0.05, **p<0.01, ***p < 0.001, Welch test for comparison between strains, Wilcoxon signed rank test for comparison between the same strain quiescent and mobility bouts.

### *RIS::unc-58gf(strong)* promotes muscle activity

An absence of voluntary movement, reduced muscle contraction and a relaxed body posture are hallmarks of sleep across species including *C. elegans* [44, 52–54]. If *RIS::unc-58gf(strong)* lacks deep sleep bouts then this should be reflected by the absence of extended periods of movement quiescence that coincide with reduced muscle activity. During wakefulness, *C. elegans* shows locomotion behavior and feeds by contractions of the pharynx muscle [55, 56]. Locomotion and pharyngeal pumping stops during sleep and the animals assume a relaxed body posture [10, 44, 52, 54]. Underlying these effects, RIS inhibits both locomotion and pharyngeal pumping [10, 12, 15, 57, 58]. To test the hypothesis that *RIS::unc-58gf(strong)* lacks long phases of muscle relaxation that are typical for sleep, we first imaged arrested larvae inside microfluidic chambers and quantified more precisely movement activity of the body and the pharynx.

To quantify mobility behavior of the head and tail regions, we tracked the positions of neurons that are located in these body parts. To calculate movement speed of the head, we tracked the position of the RIM neurons. For tracking the tail we used the position of the PLM neurons [59]. The tracking data revealed that in *RIS::twk-18gf*, both head and tail movement was similar to the wild type during wakefulness. During the few quiescence phases of *RIS::twk-18gf*, movement speed was reduced less than in the wild type, indicating that the quiescence phases that remain in *RIS::twk-18gf* are less pronounces compared with the wild type. In *RIS::unc-58gf(strong)*, head movement was decreased compared to wild-type worms, whereas there was no difference in tail movement (Fig 8A-B). *RIS::unc-58gf(strong)* worms were uncoordinated and the body was more curved (Fig S7, Movies S7 and S8), which might explain the difference of head and tail movement speeds. In the few locomotion quiescence phases of *RIS::unc-58gf(strong)*, there appeared to be less of a reduction of movement speed.

**Fig 8.**
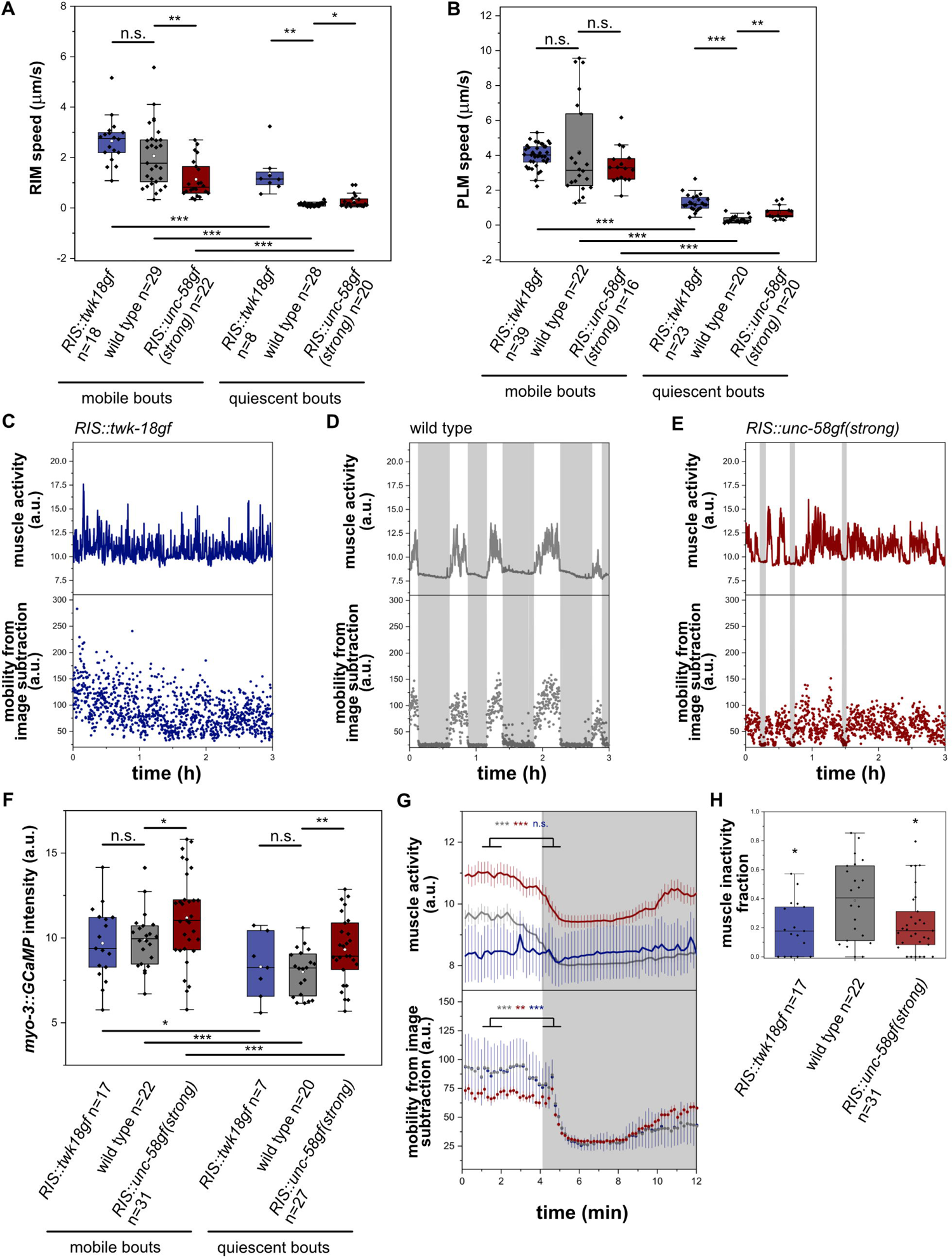
*RIS::unc-58gf(strong)* causes increased muscle activity. A) We calculated the movement speed of the RIM head neuron to measure head movement speed. RIM speed was reduced in *RIS::unc-58gf(strong)*. *p<0.05, **p<0.01, ***p < 0.001, Welch test for comparison between strains, Wilcoxon signed rank test for comparison between the same strain quiescent and mobile bouts. B) We calculated the movement speed of the PLM tail neuron to measure tail movement speed. PLM movement speed was not significantly changed in mobile bouts in *RIS::unc-58gf(strong)* and increased in detected quiescent bouts. *p<0.05, **p<0.01, ***p < 0.001, Welch test for comparison between strains, Wilcoxon signed rank test for comparison between the same strain quiescent and mobile bouts. C-E) Sample traces of muscle activity for *RIS::twk-18gf* and *RIS::unc-58gf(strong)*. F) *RIS::unc-58gf(strong)* causes increased muscle activity in quiescence as well as mobility bouts. *p<0.05, **p<0.01, ***p < 0.001, Welch test for comparison between strains, Wilcoxon signed rank test for comparison between the same strain quiescent and mobility bouts. G) A sleep bout alignment reveals that *RIS::unc-58gf(strong)* has increased muscle activity. **p<0.01, ***p < 0.001, Wilcoxon signed rank test. H) *RIS::twk-18gf* and *RIS::unc-58gf(strong)* reduced phases of muscle inactivity. *p<0.05, Welch test.

We next imaged our transgenic strains in the arrested first larval stage for 1 min with a framerate of 10Hz and manually scored pumping rate. This analysis revealed no significant difference in pharyngeal contraction rates for all strains compared to wild type (Fig S8).

To test how *RIS::unc-58gf(strong)* and *RIS::twk-18gf* affect muscle activity, we measured and quantified muscle GCaMP [52]. In the wild type, muscle calcium activity was high during wakefulness, and strongly reduced during sleep. In *RIS::twk-18gf*, muscle calcium was almost constantly elevated and resembled the activity levels of the wild type. *RIS::unc-58gf(strong)* worms showed a general increase in muscle calcium activity (Fig 8C-F). The increased muscle activity is consistent with and might contribute to the slightly uncoordinated movement of this mutant. Phases of behavioral and muscle activity dampening occurred less often, and if they occurred, then these dampened states appeared to be less deep. (Fig 8G, H).

### In *RIS::unc-58gf(strong)*, RIS calcium activity cannot be easily elevated further by optogenetic stimulation

Calcium imaging suggested that the RIS baseline in *RIS::unc-58gf(strong)* is substantially elevated and that calcium transients are lacking. We aimed to better understand the nature of these physiological changes in RIS that are caused by expression of *RIS::unc-58gf(strong)*. Based on the known characteristics of the *unc-58gf* mutant channel, the expression of *unc-58gf* might plausibly cause a constant depolarization of the membrane of RIS [32]. Depolarization in turn should lead to the activation of voltage-gated calcium channels, and RIS expresses the voltage-gated calcium channels *egl-19* and *unc-2* [31]. The conductivity of voltage-gated calcium channels is increased by depolarization with an optimum in the depolarized range. Further depolarization causes inhibition and long-term stimulation reduces conductivity by desensitization [60–65]. RIS depolarization in *RIS::unc-58gf(strong)* could thus perhaps be increased to allow for elevated RIS baseline calcium activity, but might also inhibit further activation of calcium channel activity. We tested the ability of RIS to activate by optogenetically stimulating RIS in *RIS::unc-58gf(strong)* inside microfluidic chambers and monitored the calcium activity of this neuron as well as the motion behavior of the larvae. For this experiment, we used the red-shifted channelrhodopsin variant ReaChR in combination with GCaMP3 [12, 22, 57]. In the wild type, optogenetic activation caused a strong increase in intracellular calcium and a strong induction of motion quiescence, which is consistent with previous results [12, 15, 22, 57]. By contrast, optogenetic activation of RIS in *RIS::unc-58gf(strong)* did not increase RIS activity further and did not induce behavioral quiescence (Fig 9A-B). This result is consistent with the view that *RIS::unc-58gf(strong)* causes a heightened level of RIS activity that cannot be easily increased further with optogenetic activation. Hence, in *RIS::unc-58gf(strong)*, RIS appears to be locked in a state of heightened activity while further activation is inhibited.

**Fig 9.**
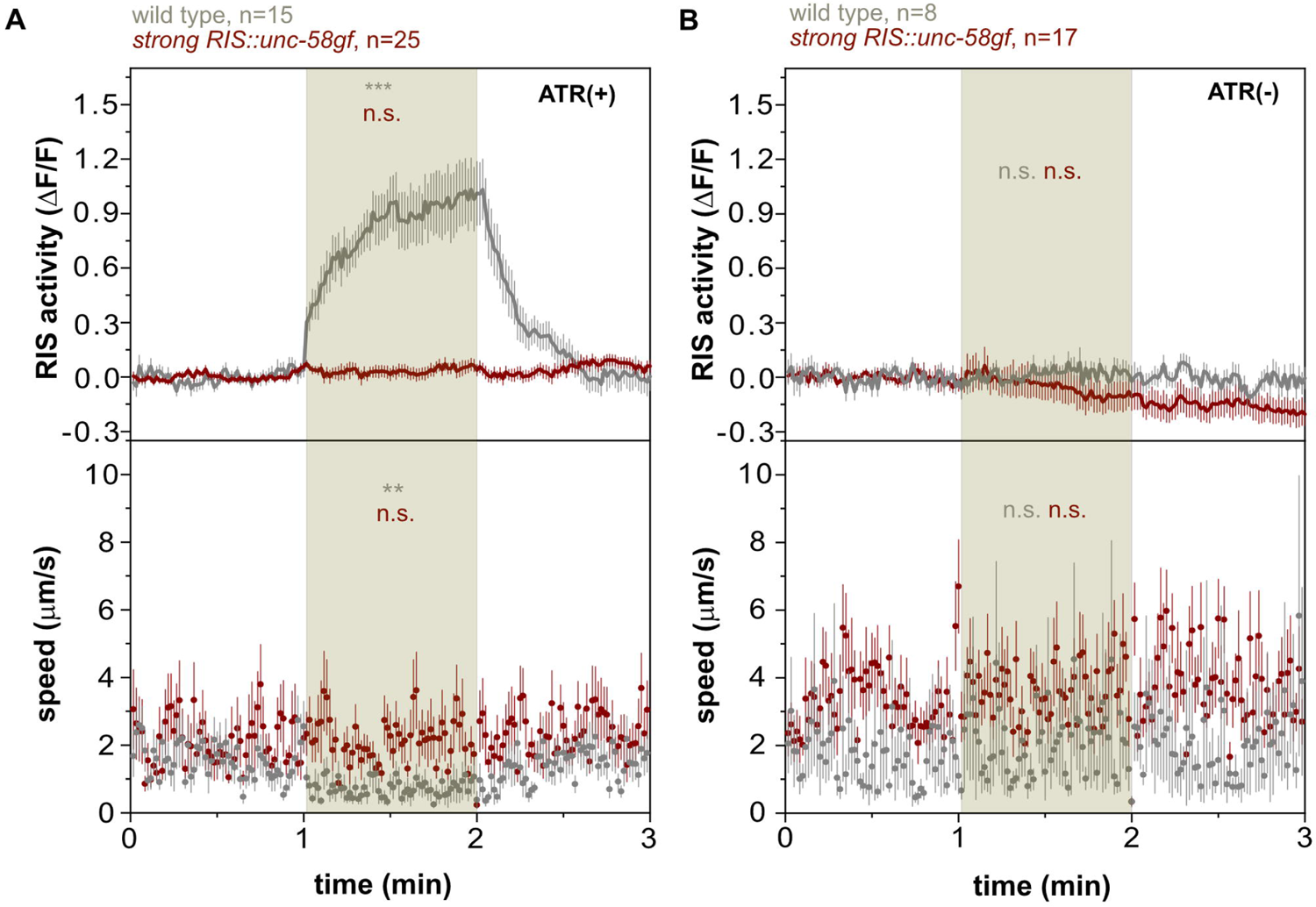
*RIS::unc-58gf(strong)* is refractory to optogenetic activation of RIS A) Optogenetic activation of RIS during the stimulation time (minute 1-2) leads to an increase of RIS activity and reduced mobility in wild-type worms but it does not affect the strong RIS activation strain. RIS speed was measured. A statistical comparison was made between baseline (0-1min) and stimulation (1-2min). n.s. p>0.05, **p<0.01, ***p<0.001, Wilcoxon signed rank test. B) Control for the optogenetic experiment. Without ATR treatment, neither the wild type nor the strong RIS activation strain, shows a response to green light. N.s. p>0.05, Wilcoxon signed rank test.

### Long-term optogenetic activation of RIS increases RIS baseline activity and inhibits activation transients

*RIS::unc-58gf(strong)* appears to cause increased RIS activity that cannot easily be increased further. This could hypothetically be explained by a high level of depolarization in *RIS::unc-58gf(strong)* that causes both RIS baseline elevation as well as inhibition of RIS calcium transients. To test the idea that constant depolarization elevates baseline RIS activity while inhibiting activation transients, we used long-term optogenetic activation. We used again ReaChR to depolarize the membrane and followed the activity of RIS. Optogenetic tools like ReaChR typically display a brief photocurrent peak activity within the first second of light exposure and then plateau at a lower photocurrent level for an extended period of time [66]. We previously used ReaChR to establish long-term optogenetic manipulations in *C. elegans* [23]. To test for the activation of RIS during prolonged optogenetic activation, we activated ReaChR in RIS with orange light for 11h and followed the activity of RIS using GCaMP and measured mobility of the larvae that were kept inside microfluidic chambers. During the first hour of optogenetic stimulation of RIS, there was a strong increase of calcium activity as well as of behavioral quiescence (Fig 10A-C). After this first peak period, the ongoing optogenetic stimulation led to lower but more constant increase of RIS baseline activation (which was, however, weaker than the activation observed in *RIS::unc-58gf(strong)* average calcium baseline activity increase in *RIS::unc-58gf(strong)* compared with wild type was 1.55, and average calcium activity baseline increase from 2-11h of activation in *RIS::ReaChR* compared with wild type was 1.25) (Fig 10D, E). Long-term optogenetic activation caused reduced calcium transients, and the residual calcium transients did not correlate with phases of immobility. This indicates that long-term optogenetic activation has an inhibitory effect on RIS calcium transients (Fig 10F). The optogenetic long-term RIS activation did not cause a measurable decrease in behavioral quiescence (Fig 10C). Thus, long-term optogenetic stimulation leads to a constantly increased RIS baseline activity and an inhibition of RIS calcium transients that is similar to, but weaker than, the activation observed in *RIS::unc-58gf(strong)*. This difference in effect strength might be explained by the difference in ion conductivity of these channels [32, 66]. These data indicate that long-term optogenetic stimulation can lead to a constant increase in baseline RIS calcium activity at a level that is above the baseline activity without optogenetic stimulation, yet lower than the maximum possible optogenetic RIS activation peak intensity. These optogenetic data are consistent with the idea that increased depolarization of RIS leads to a constant elevation of baseline calcium activity and impairs calcium transients.

**Fig 10.**
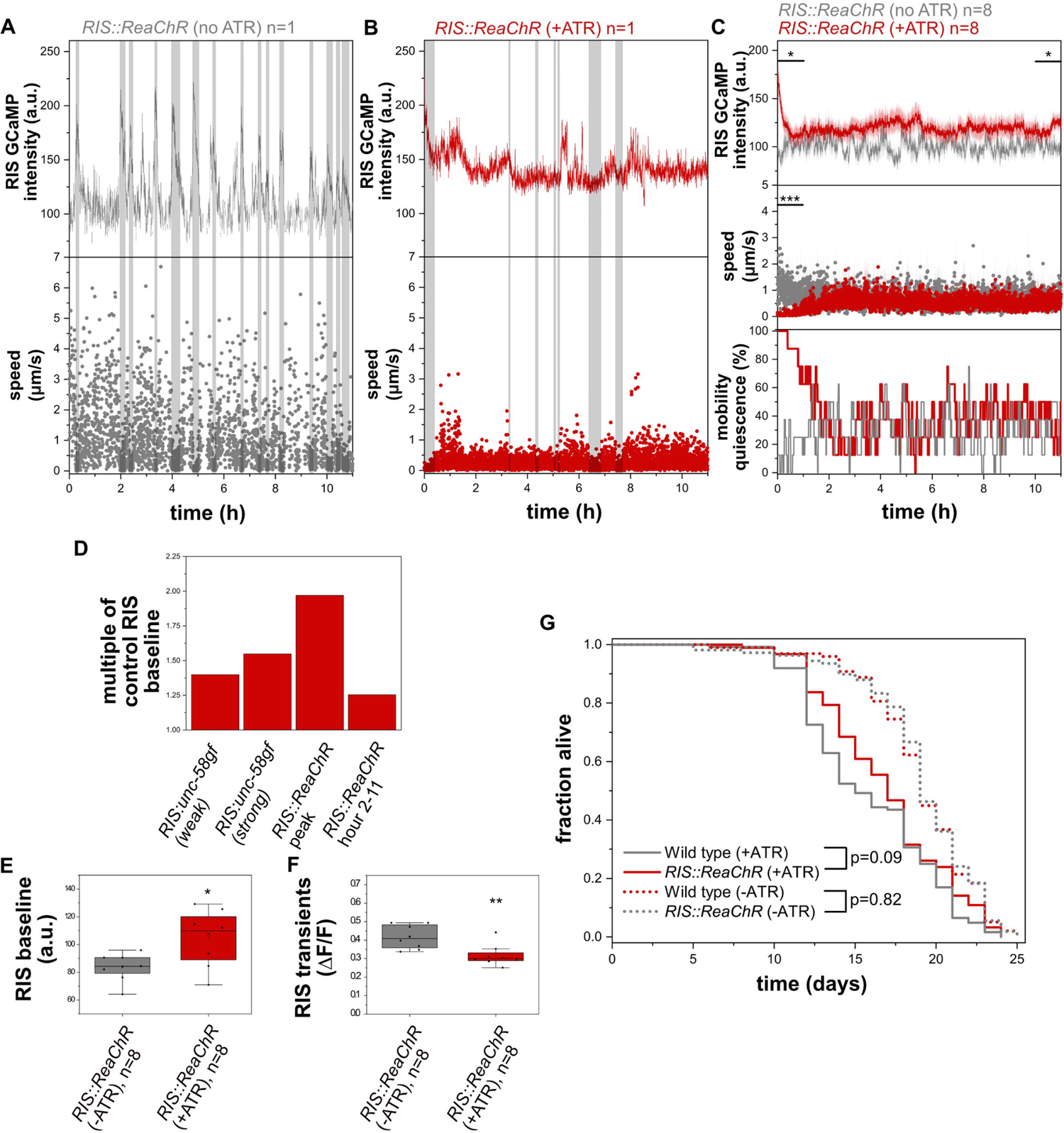
Long-term optogenetic activation of RIS increases overall RIS calcium activity and promotes survival A) Sample trace of control larvae without retinal in which RIS cannot be activated optogenetically. RIS speed was measured. B) Long-term optogenetic RIS activation by orange light in the presence of retinal. C) Long-term activation of RIS by ReaChR leads to an initial strong increase in calcium activity in RIS and simultaneous mobility quiescence followed by a mild RIS activity increase yet no increase in movement speed or movement quiescence. D) Comparison of RIS activity level changes caused by *RIS::unc-58gf(strong)* and by optogenetic activation of RIS. The data was normalized to their respective controls. E) Long-term optogenetic activation of RIS significantly increases RIS baseline. *p<0.05 Welch test. F) Transients in RIS are significantly reduced in magnitude and don’t correlate with quiescence. **p<0.01 Welch test. G) Long-term optogenetic activation of RIS inside microfluidic chambers leads to a small increase in survival (p=0.09 Logrank test, p=0.02 Fisher’s Exact Test, the average of all individual worms of all three technical replicates was averaged, n=92 for *RIS::ReaChR*(+ATR), n=98 *RIS::ReaChR*(-ATR), n=124 wild type (+ATR), n=108 wild type (-ATR))

### Long-term optogenetic activation of RIS causes a small increase in survival

Our data using *RIS::unc-58gf(strong)* indicated that extended survival of L1 arrest is associated with increased overall RIS activity. To confirm this result with an independent approach, we used *RIS::ReaChR* for long-term activation of RIS using the OptoGenBox system [23]. We kept L1 larvae inside microchambers and constantly stimulated the entire chambers with orange light. The optogenetic stimulation was only paused briefly once every day to monitor the survival of the larvae. The stimulation protocol was applied until all larvae had died. The functionality of ReaChR in *C. elegans* requires the addition of all-trans retinal (ATR). Addition of ATR shortened the survival (Fig 10G). We hence compared the survival of N2 exposed to ATR to worms expressing ReaChR that were also exposed to ATR. By this comparison, optogenetic long-term RIS activation caused a small trend for increased survival that was statistically significant according to Fisher’s Exact Test but not significant according to the Logrank test (p=0.02 Fisher’s Exact Test, p=0.09 Logrank, Fig 10G). While this result borders statistical significance, it is consistent with the view that activation of RIS promotes survival. The small effect size is consistent with the small RIS activity increase caused by extended optogenetic stimulation. Thus, this experiment supports the view that RIS activation promotes survival of L1 arrest.

### Lethal blue-light stimulation activates RIS, and RIS supports survival of lethal blue light

Disturbing sleep by sensory stimulation leads to arousal that forces wakefulness and prevents the dampening of underlying neuronal circuits [12, 20, 22, 44, 67, 68]. Sleep disturbance or deprivation can cause increased RIS activation [22, 24, 69]. The high level of RIS activity in turn could be protective under such conditions and promote survival [24]. Thus, *RIS::unc-58gf(strong)* might cause a state that is similar to a state caused by prolonged sleep deprivation or disturbed sleep. In both, disturbed sleep and *RIS::unc-58gf(strong)*, the activity of wakefulness circuits is not dampened yet coincides with elevated RIS activity.

To further illustrate the idea that sensory stimulation activates RIS while impairing sleep, we stimulated worms with a lethal intensity of blue light. We used arrested L1 larvae after 48h in the absence of food, a condition during which the animals normally show regular RIS activation transients and sleeping behavior [12]. We kept L1 larvae in microchambers and repeatedly applied pulses of blue light while monitoring RIS activity and motion behavior. For comparison, we used control animals that were exposed to shorter pulses of blue light of only one tenth the length, thus still allowing for calcium imaging. Control larvae displayed sleeping behavior detected by motion quiescence and RIS calcium activity during about a quarter of the time (Fig 11A, B). Strong blue-light stimulation caused an increase in RIS calcium activity compared with control animals (after 1h of strong blue-light stimulation, the activity of RIS was approximately 5.6-fold higher compared with worms that were stimulated with a lower dose). Many of the calcium transients were very short, however, and did not coincide with increased motion quiescence (Fig 11B-C). Hence, the stronger blue-light stimulus increased RIS activity but apparently did not increase sleep behavior.

**Fig 11.**
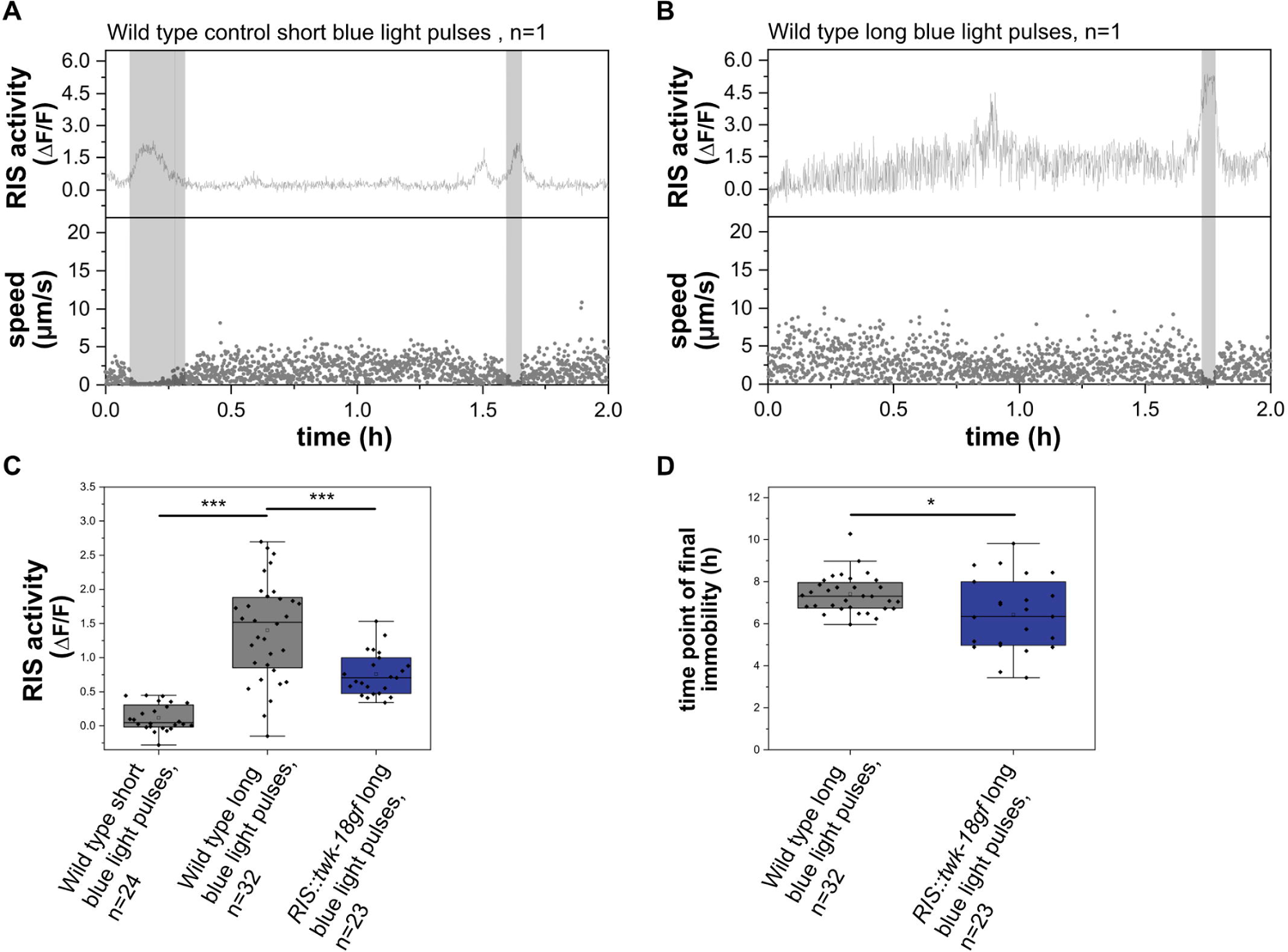
RIS activates upon lethal blue-light stimulation and extends survival A) Sample trace for a control worm stimulated by a short blue light stimulus. RIS speed was measured. B) Sample traces of wild-type worms upon stimulation with a long blue-light stimulus. The grey shaded area presents detected time in motion quiescence. C) The strong stimulus increased normalized RIS activity over baseline compared to control worms. ***p<0.001, Welch test. D) Wild-type worms have a survival advantage over RIS-inhibited worms upon exposure to the lethal blue-light stimulus. *p<0.05, Welch test.

To probe for the role of RIS in survival of lethal blue-light stimulation, we compared wild-type and *RIS::twk-18gf* animals, in which RIS activated less under blue-light stimulation (Fig 11C). We scored the time point at which the animals became terminally immobile, which presents an established proxy to score for death [70]. Control animals with functional RIS died around 7-8h after strong blue-light stimulus onset. RIS-impaired animals already died at around 6h after stimulus onset. The presence of functional RIS hence extended survival by 15% (Fig 11D).

Thus, stimulation by a lethal blue-light stimulus can increase RIS calcium activity without increasing RIS-dependent sleep behavior. While all animals were eventually killed by the blue-light stimulus, RIS activity slightly extended survival. The blue-light experiment thus supports the idea that RIS is a protective neuron that activates upon stressful stimulation. As in *RIS::ReaChR* and during long-term optogenetic stimulation of RIS, increased RIS activity was not associated with increased sleep behavior.

### *flp-11* is a key neuropeptide gene required for quiescence behavior in L1 arrest

What are the key transmitters that RIS uses to control sleep during L1 arrest? GABA is expressed in RIS but has not been shown to play a substantial role in promoting sleep [10, 15]. FLP-11 is a group of neuropeptides of RIS that present the major transmitters of RIS required for sleep induction during development [11]. Expression from the *flp-11* promoter depends on neuronal activity levels [71], and is increased in *RIS::unc-58gf(strong)* (Fig S3B), suggesting that FLP-11 might also be a major transmitter for sleep induction during L1 arrest. To directly test the role of FLP-11 in quiescence during L1 arrest, we measured the effects of *flp-11* deletion (*flp-11(-)*) using microfluidic chamber imaging [11]. *flp-11(-)* strongly reduced quiescence behavior (Fig 12A). *RIS::unc-58gf(strong)* still displayed a small fraction of quiescence behavior during L1 arrest. To test whether this residual quiescence was caused by *flp-11*, we deleted the entire *flp-11* gene in the *RIS::unc-58gf(strong)* operon locus (Fig S9A) and quantified quiescence behavior. The sleep loss of *RIS::unc-58gf(strong)* was reduced further by *flp-11(-),* suggesting that the residual quiescence behavior in *RIS::unc-58gf* indeed depends on *flp-11* (Fig 12A).

**Fig 12.**
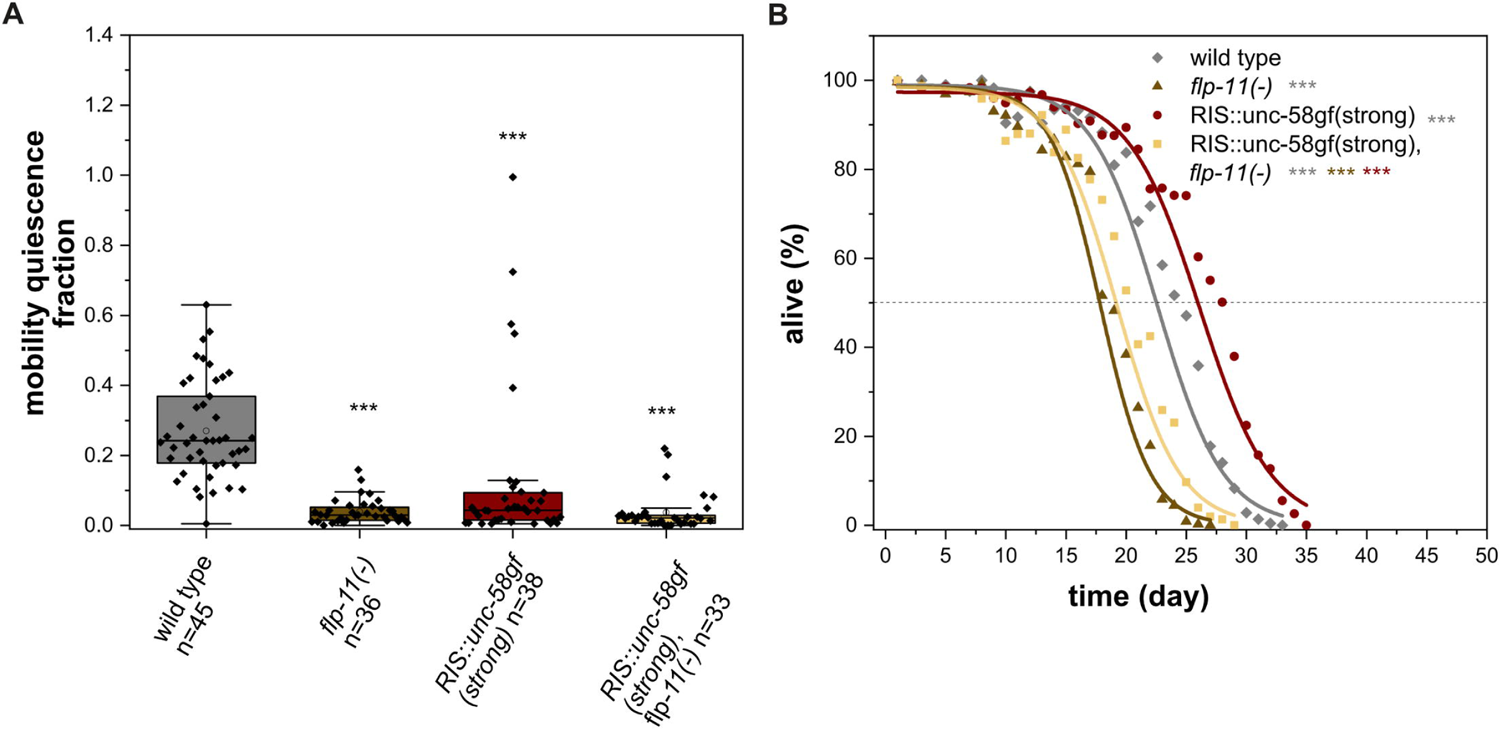
FLP-11 is required for sleep during L1 arrest A) Deletion of the neuropeptide gene *flp-11* causes loss of sleep. Immobility in *RIS::unc-58gf(strong)* is increased when *flp-11* is deleted. ***p<0.001, Welch test with FDR correction for multiple testing. B) Loss of the FLP-11 neuropeptides leads to strongly reduced survival (three-parameter logistic fit, see also supplementary table S1, Fisher’s Exact Test was conducted on day 18) for comparisons when *flp-11(-)* was the shortest lived condition, day 20) when *RIS::unc-58gf(strong), flp-11(-)* was the shortest lived condition or day 24) when wild type was the shortest lived condition. The p-values were FDR corrected with Benjamini-Hochberg procedure with a 5% false discovery rate. The plot includes data from three replicates. ***p<0.001.

### FLP-11 neuropeptides support survival

What is the key transmitter of RIS that promotes survival? RIS is GABAergic, but GABA has been suggested to be a negative regulator of lifespan [72]. We showed above that *flp-11* is required for most of the immobility during L1 arrest sleep. We hence tested whether FLP-11 neuropeptides are also required for survival. Deletion of *flp-11* reduced survival by 24.7%, suggesting that *flp-11* contributes to survival (Fig 12B). To test whether *flp-11* is required for increased survival caused by *RIS::unc-58gf(strong)*, we measured survival in the *RIS::unc-58gf(strong), flp-11(-)* strain. *flp-11(-)* suppressed almost the entire survival extension caused by *RIS::unc-58gf(strong)* (Fig 12B). Similarly, the recovery upon feeding was suppressed by *flp-11(-)* in *RIS::unc-58gf(strong)* (Fig S9B). Together, these experiments indicate that *flp-11* plays an important role in survival and recovery of L1 arrest both in the wild type and in *RIS::unc-58gf(strong)*.

These results suggest that *flp-11* plays two roles, one role is to promote sleep and the other is to promote survival. Deletion of *flp-11* blunts survival less than strong RIS inactivation, indicating that additional neurosecretory factors might also contribute to survival. As RIS expresses multiple neuropeptides and also GABA [11, 15, 17], it may use multiple transmitters to control sleep behavior and survival. In summary these data suggest that FLP-11 neuropeptides are the major transmitters released by RIS to promote both sleep as well as survival during L1 arrest.

## Discussion

Tools to manipulate neuronal activity are instrumental for studying neuronal circuits and physiology. Here we establish the constitutively active sodium channel gene *unc-58gf* as a tool for activation of a neuron that normally does not express *unc-58* [31, 32]. Modest expression of *unc-58gf* in RIS during L1 arrest caused an increased baseline activity of RIS and increased the probability of RIS to experience calcium activation transients. It thus appears that *unc-58gf* can activate RIS when expressed at lower level. Strong expression of *unc-58gf* in RIS caused constant high levels of baseline RIS activity yet prevented further activation transients. When expressed highly, *unc-58gf* thus locks RIS activity at a level that is higher than the normal baseline and at the same time also inhibits RIS calcium transients. Thus, *unc-58gf* can be used as a tool to create different and complex patterns of neuronal activity, but the level of expression is crucial to achieve different effects. Together with constitutively active potassium ion channels that cause various levels of inhibition, activity of a neuron can be modulated from strong inhibition to constant elevation of baseline activity.

RIS inhibits wakefulness neurons to cause sleep behavior and it promotes survival under stressful conditions such as larval starvation or wounding [10, 12, 18, 24, 35]. A parsimonious explanation linking these two observations would be that sleep promotes survival, for example through energy conservation or behavioral optimization [27]. Energy conservation and behavioral optimization certainly are important functions of sleep behavior [28]. In addition, sleep neurons could promote physiological benefits by mechanisms that do not necessarily depend on sleep behavior.

How can the manipulation of neuronal activity be employed to solve how neurons control behavioral and physiological function? Here we used transgenic expression of ion channel mutants in RIS to shed light on the relationship between RIS activity, sleep behavior and survival. *RIS::unc-58gf(strong)* promoted survival, but also inhibited sleep behavior. Thus, *RIS::unc-58gf(strong)* allowed for an uncoupling of sleep behavior and survival functions of RIS. Our data thus suggest that the sleep-active RIS neuron can promote survival independently of sleep.

What are hypothetical mechanisms that allow for the uncoupling of sleep behavior and survival benefits of RIS? The loss of sleep behavior in *RIS::unc-58gf(strong)* could hypothetically be explained by the absence of calcium transients in RIS. This could, for example, be caused by desensitization of wakefulness circuits that are downstream of RIS. According to this hypothesis, wakefulness circuits would decrease their activity with increasing RIS activity, and would start getting more active again only after a period of stagnating or decreasing RIS activity. Both, the induction of sleep behavior and the promotion of survival by RIS depend on FLP-11 neuropeptides, which are thought to be released upon RIS activation [11]. The two functions of FLP-11 might mechanistically separate at the level of divergent receptors, signaling pathways or cell types downstream of FLP-11 that mediate either the sleep or the survival response. For example, the receptors and downstream cell types that mediate the effects of FLP-11 on dampening wakefulness circuits might desensitize quickly, thus limiting the amount of behavioral sleep that can be induced by FLP-11 released from RIS. Along these lines, RIS activity might exert a survival-promoting effect through a mechanism with different desensitization properties. Survival appears to be a function of RIS activity, depends on *flp-11*, but does not seem to depend on activation transients. Thus, the underlying signaling components do not seem to desensitize quickly but might respond to lower levels of RIS activity or might integrate RIS activity over time. This could hypothetically be achieved by FLP-11 activating downstream receptors and cell types that react sensitively to FLP-11 concentration, that do not easily desensitize and that promote survival upon sensing FLP-11. The key FLP-11 receptors required for promoting sleep and survival and the cells in which these receptors function have not yet been identified [11]. The identification of FLP-11 receptors will allow for further studies of the mechanisms by which FLP-11 causes sleep behavior and survival. Another hypothetical explanation for changes in sleep behavior upon expression of ion channel mutants might be changes in the underlying neuronal connectivity. Further experimentation is required to test these speculative hypotheses.

What is the mechanism and role of homeostatic compensation during sleep deprivation? Homeostatic regulation can compensate for lost or disturbed sleep across species and plays an important role in the regulation of sleep in *C. elegans* [22, 53, 67, 69, 73]. The typical effects of disturbing sleep or depriving an animal of sleep are a reduction of sleep behavior caused by continued activity of wakefulness circuits that steer behavioral activity. Increased behavioral activity is often accompanied by a subsequent overactivation of sleep-active neurons that promote a rebound phase of increased sleep duration or intensity. Rebound sleep in turn is thought to compensate for lost or disturbed sleep, at least partially [8, 22, 24, 67, 69, 73, 74].

How is *RIS::unc-58gf(strong)* related to sleep deprivation and sleep homeostasis? Does *RIS::unc-58gf(strong)* trigger homeostatic compensation of the lost sleep? Does the transgene cause a state that is similar to the effects of sleep-deprivation? To answer these questions, we first studied behavioral sleep across stages. Across the life of arrested larvae, behavioral quiescence was reduced in *RIS::unc-58gf(strong)*. Furthermore, sleep was reduced by *RIS::unc-58gf(strong)* during L1 lethargus and after stress in the adult. We also studied the activity of excitable cells in *RIS::unc-58gf(strong)* in arrested L1 larvae. We studied the activity of sensory neurons, interneurons and body wall muscle. All of these excitable cells are known to be rather inactive during sleep and thus the study of the activity of these excitable cells allows for a quantitative description of sleep depth. Our data showed that the sleep bouts in *RIS::unc-58gf(strong)* are less pronounced and hence appear to be less deep. Hence we could not find any signs of deeper or compensatory sleep in *RIS::unc-58gf(strong)*. By contrast, the analysis of excitable cells suggested that the residual bouts of behavioral quiescence in *RIS::unc-58gf(strong)* were generally weakened. *RIS::unc-58gf(strong)* thus displays the neuronal signature of sleep deprivation or disturbed sleep, i.e. the combination of impaired wakefulness circuit dampening and over activation of RIS. Thus, *RIS::unc-58gf(strong)* might be understood and employed as a model of disturbed sleep or of sleep deprivation.

Does RIS activate when the animals are disturbed to promote survival [12, 18, 22, 24, 69]? To further illustrate the idea of RIS acting as a promoter of survival, we disturbed animals during sleep via a lethal blue-light stimulus. Lethal blue light increased RIS activity and functional RIS was required for extending the survival of the animals during blue-light stimulation yet did not increase sleep behavior. This supports the idea that RIS activates upon sensory stimulation to promote survival independently of increasing sleep behavior.

What is the mechanism by which RIS promotes survival? Is there an important role of RIS to conserve energy by shutting down behavior? The reduction of brain and muscle activity likely contributes to survival by conserving energy. While energy consumption appears to be playing a role in survival, behavioral activity and thus energy consumption does not correlate with survival when transmission from RIS is strongly increased. In *RIS::unc-58gf(strong)* animals, neuronal, muscle and behavioral activity are high and consume energy, yet survival is increased. Hence RIS appears to be able to promote survival independently of sleep behavior and despite the energy loss that can be expected to be caused by the increased activity of excitable cells [28]. As both sleep and survival are functions of RIS activity over a wide range of conditions, the two processes should be coupled during normal sleep. During each sleep bout, RIS depolarizes and thus should promote both, sleep and survival. Such coupling of sleep and survival during normal sleep might ensure that phases of brain and behavioral inactivity coincide with phases during which survival-promoting processes are active. Tight coupling might serve to optimize energy expenditure, as energy conserved by sleep could be directly allocated to survival-promoting processes [28].

How does RIS cause physiological protection and increased survival independently of conserving energy? RIS promotes survival of stressful conditions and underlying changes in protective gene expression [12, 18, 24]. Together with the apparent strong reduction of sleep behavior in *RIS::unc-58gf(strong)*, this might suggest that the role of RIS overactivation upon disturbance or sleep deprivation is not only to promote rebound sleep behavior, but also to promote protective physiological benefits that support survival. An important function of RIS might thus be to promote survival when the animal is disturbed by sensory stimulation. Disturbance, e.g. by blue light, might directly be stressful and thus require an increased survival response. Disturbed sleep is often associated with stress, for example when mechanical stimuli are experienced. Hence RIS might hypothetically also act as part of a sensory system that activates during stimulation to signal an increased demand of physiological protection.

The FOXO transcription factor DAF-16 has been shown to be linked to the control and functions of sleep. It is required for sleep under various conditions including during L1 arrest. Also, FOXO has been shown to be activated upon sleep deprivation during both lethargus as well as during L1 arrest and to be required for increased rebound sleep following sleep disturbances [24, 58, 67, 73] (Fig S10). A plausible model hence suggests that RIS activation promotes DAF-16-dependent gene expression changes in order to support survival [24]. Our data indicate that RIS strongly promotes protective gene expression despite the loss of sleep behavior. Thus, *RIS::unc-58gf(strong)* intensifies the survival function of RIS while inhibiting the sleep function of RIS.

By controlling the activity of RIS, we showed that the survival function and the sleep behavior function of this neuron can be decoupled. By constantly elevating RIS baseline activity, we generated a condition in which wakefulness neural activity and sleep-active neuron activity coexist. A similar condition often underlies disturbed sleep. Our data suggest that when sleep is disturbed, RIS promotes survival despite the loss of behavioral sleep. It would be interesting to study whether sleep neurons in other species also can promote physiological benefits in the face of lost sleep.

## Materials and Methods

### Resource and Materials Availability

Key *C. elegans* lines that were used in this project are available at the Caenorhabditis elegans Genetics Center. Other *C. elegans* strains, plasmids, or additional reagents generated for this study are available upon reasonable request.

## Data Availability

All relevant data are within the manuscript and its supporting Information files. Matlab scripts can be found at Github (https://github.com/ibusack/A-sleep-active-neuron-can-promote-survival-while-sleep-behavior-is-disturbed).

## Experimental model and subject details

*C. elegans* hermaphrodites were maintained at 20°C. They were kept on Nematode Growth Medium Plates (NGM). The *Escherichia coli* strain OP50 served as a food source during maintenance [75].

## Method Details

### Crossing *C. elegans* strains

Males were generated via heat shock or maintained by a continuation of crossings of worms of the same genotype [76]. Single worms of the F2 generation were genotyped using duplex PCR genotyping [77]. For crossing fluorescent transgenes, fluorescent markers were used. All strains and primers of this study can be found in the supplementary tables S2 and S3.

### Creating mutants with the CRISPR/Cas9 system

We designed the sequences of the following transgenes in silico. Transgenes were synthesized and inserted into the genome by using CRISPR/Cas9 by SunyBiotech. Transgene sequences were verified by Sanger sequencing.

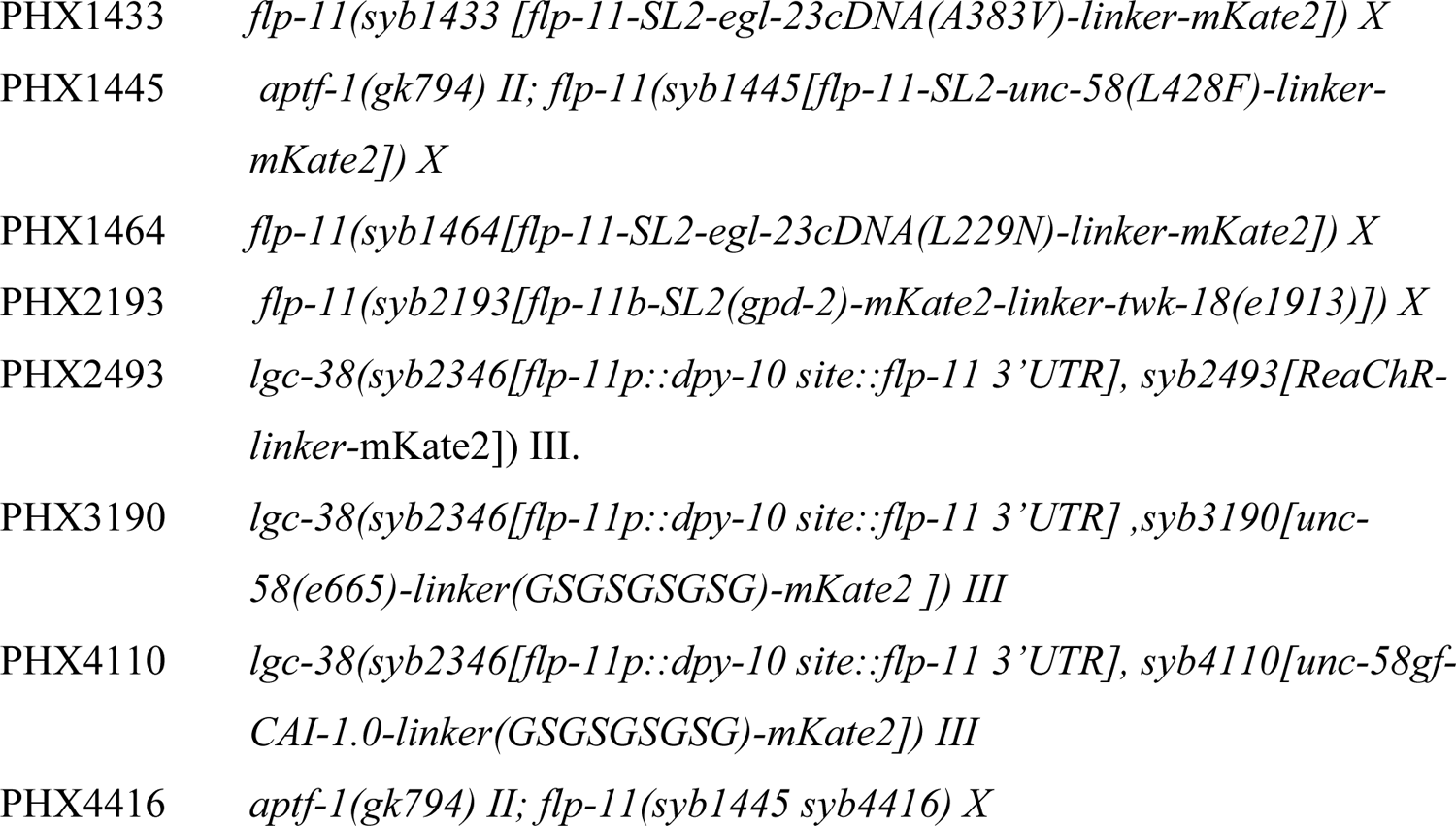

The coding sequences of the ion channel transgenes were synthesized and consisted each of the spliced coding sequence of the ion channel including the mutation that caused the leak current [32], a nine amino acid long C-terminal linker sequence (GSGSGSGSG) followed by a C-terminal mKate2 gene. The coding sequence of mKate2 was codon optimized and two introns were inserted [78]. The coding sequences of the ion channels were not codon optimized to allow for proper folding of the membrane proteins. We tried codon-optimization [78] of the *unc-58(L428F)* channel gene and mKate2 to achieve high expression, but this also caused aggregation of the fluorescent fusion protein (Fig S1A, D). Hence, we used the endogenous gene sequence for the channel and only codon-optimized mKate2. The sequences of the ion channel transgenes can be found in the supplemental information.

Analysis of the RIS transcriptome showed that *flp-11* has the strongest and most specific promoter of all RIS-expressed genes, with 157-fold higher expression in RIS than in other neurons [17]. *flp-11* is also expressed in other neurons, albeit at low levels [11, 17, 31]. Using imaging settings that cause oversaturation of the mKate2 signal in RIS revealed that there is a weak expression of *unc-58gf::mKate2* in a few neurons in the ventral cord in *RIS::unc-58gf(strong)* (Fig S1F). Expression from the *flp-11* promoter in RIS but not in other neurons depends on *aptf-1* [11]. We hence imaged mKate2 expression in *aptf-1(-).* Consistent with previous results [11], mKate2 expression was abolished in RIS but was still present in the ventral cord (Fig S1G). Thus, expression from the *flp-11* promoter is strong, has a high level of RIS specificity, and RIS expression depends on *aptf-1*.

The sequences of the ion channels of the strains PHX1433, PHX1445, PHX1464 and PHX2193 were inserted after the stop codon of the *flp-11* endogenous coding sequence together with an SL2 site. Thus, an operon was created that expressed both *flp-11* and the ion channel gene from the *flp-11* promoter. The transgenes of PHX2493, PHX3190 and PHX4110 were generated via a single-copy knock-in loci for defined gene expression (SKI LODGE) system [34]. For the SKI LODGE expression in RIS we first inserted the minimal *flp-11* 3’ and 5’UTRs separated by a *dpy-10* sgRNA cutting site. The expression from these minimal UTR sequences allowed specific expression in RIS, but the expression levels were lower compared with the endogenous *flp-11* locus (Fig 1A-D, S1A-D). The final sequences of all transgenes can be found in the supplementary Text S1.

### Imaging using microfluidic devices

Worms were filmed in microfluidic devices made from agarose hydrogel as previously described [37, 38]. In brief, a PDMS mold was used to cast box-shaped indentations from melted 5% agarose diluted in M9 buffer. After agarose hydrogel solidification the stamp was removed resulting in hundreds of microchambers sized 110×110×10μm for L1 arrest chambers or 190×190×15μm for L1 lethargus chambers per agarose slab. Depending on the experiment, approximately 9-36 pretzel-stage eggs per strain were then transferred without food for L1 arrest or with food for lethargus into the indentations in the hydrogel and the chambers were closed with a coverslip and kept in a temperature-controlled dish. Hatching of the larvae in the absence of food caused L1 arrest. Chambers were timed so that the worms were incubated for 48h at 25°C before imaging. The number of biological replicates (n) imaged in microfluidic devices is stated in each figure. One or several technical replicates (different microfluidic devices) were imaged for each experiment.

### Software for imaging

All image acquisition was performed with Nikon NIS Elements 5 software (Nikon, Tokyo).

### Dark Differential Contrast (DIC) Imaging

Long-term dark differential contrast (DIC) images were acquired with a standard 100W halogen lamp and an infrared filter (Semrock Brightline HC 785/62) (Idex Health and Science, New York). The images for L1 arrest imaging were taken 48h upon starvation for 12h with a frame rate of 0.1Hz. A 20× objective for imaging was combined with a 0.7 lens. Lethargus data was acquired with similar settings except the frame rate was 0.2Hz.

### Functional Ca^2+^-imaging

For calcium imaging we used either a back-illuminated sCMOS camera (Photometrics Prime 95B, 1200×1200 pixels) (Fig 3, 4, 8C-H, S3) or an Andor iXon EMCCD camera (512×512 pixels, Andor Technology Ltd., Belfast). (Fig 6, 7, 8A-B, S5, S6). The gain of the iXon camera was set to 200. The iXon camera’s read out mode was 10MHz at 14-bit. The images of the Photometrics camera were binned 2 × 2.

A 20× objective was used to simultaneously image 3 x 3 adjacent microchambers containing one worm each (This configuration was used to obtain data for Fig 6, 7, 8A-B, S5, S6). A 10× objective combined with a 1.5x lens was used to simultaneously image 6 x 6 adjacent microchambers containing one worm each (this configuration was used to obtain data for Fig 3, 4, 8C-H, S3).

For calcium imaging we utilized a 480nm LED and a standard GFP filter set (EGFP, Chroma). The EMCCD camera TTL signal triggered the LED. The 490nm intensities were set to either 0.23mW/mm^2^ with an exposure time of 50ms (Fig 3A, 4A-H, 8C-H), 0.33mW/mm^2^ (Fig 6A-D) or 0.51 mW/mm^2^ (Fig 6E-H), both with an exposure time of 5ms or 0.33mW/mm^2^ with an exposure time of 10ms (PLM). Additional mKate2 fluorescence images were acquired for the RIS, RIM and PLM activity experiments. For this measurement we used a 585nm LED and a set of standard Texas Red filters (Chroma). The 585nm LED intensity was set to 0.11mW/mm^2^ while imaging with a 50ms exposure time (RIS) and to 0.15 mW/mm^2^ for acquiring images with a 100ms exposure time (RIM, PLM). Calcium and mKate2 imaging were carried out with a frame rate of 0.1Hz for 3h. As the ion channels were additionally tagged with an mKate2, the parental strains without the GCaMP/mKate2 construct were imaged with the same settings with the 585nm LED. Here, no signal could be observed in the parental strains.

For the long-term calcium imaging from day 1 to day 12 we utilized slightly different LED light intensities. Here, the 490nm intensity was set to 0.34mW/mm^2^ and the 585nm light had an intensity of 0.21mW/mm^2^.

### Optogenetic activation

For optogenetic RIS activation experiments, the agarose containing the microchambers was supplemented with 10μl of 10mM all-trans-retinal (ATR, Sigma Aldrich) dissolved in ethanol 1h before the start of image acquisition. In experiment in Fig 9, A baseline of GCaMP images was recorded for 1min with a frame rate of 0.33Hz. This was followed by a stimulation phase of 1min, in which again GCaMP images were acquired with a frame rate of 0.33Hz. In between the camera exposures, the worms were illuminated with 585nm LED light with an intensity of 0.17mW/mm^2^ for 800ms triggered by the software. After the stimulation period, a recovery period of 1min was imaged, in which GCaMP images were acquired with a frame rate of 0.33Hz. The 490nm LED intensities for imaging were set to 0.39mW/mm^2^ and images were acquired with an exposure time of 5ms.

For the long-term optogenetic imaging worms (Fig 10), a stimulation phase of 11h was recorded in which GCaMP images were acquired with a frame rate of 0.33Hz. In between the camera exposures, the worms were illuminated with 585nm LED light with an intensity of 0.17mW/mm^2^ for 800ms triggered by the software.

### Quantification of expression of mKate2 of the RIS activity strains

Larvae were synchronized via bleaching [17] and hatched in M9 buffer at 25°C. Imaging of most strains was performed after 48h in the L1 arrest stage by transferring worms with a 10 µl liquid drop onto an agarose pad (5% agarose solved in M9 buffer). The *RIS::unc-58gf and flp-11(-)* worms were imaged 24h after bleaching. Larvae were then fixed through the addition of 10 µl of 25mM levamisole (Sigma Aldrich).

To image the expression levels of the different RIS activity transgenes, mKate2 was imaged with an Andor Revolution disc system (Andor Technology Ltd.) and a CSU-X1 spinning disc head (Yokogawa). The 561nm laser light intensity was set to 0.49mW/mm^2^. The worms were imaged through a 100× oil objective and by an Andor iXon Ultra EMCCD camera (1024×1024 pixels) (Andor Technology Ltd., Belfast). An EM gain of 200 was included and the camera read-out time was set to 30MHz. A z-stack of 21planes, each 0.5μm apart (10μm in total) was acquired. For comparison, a maximum z-stack projection was calculated.

### DIC image analysis

Mobility of worms was analyzed from DIC images though image subtraction as previously described [17, 79]. The matrix of intensity values of the preceding image was subtracted from the matrix of intensity of the following image. A mean of all values of the difference matrix was then calculated to attain a mean smoothed image subtraction value, which represents the movement of the worm.

To identify lethargus in fed worms, the non-pumping time prior to molting was identified. For the sleep analysis in recovery lethargus after 12 days of starvation followed by refeeding, 3h prior to molt were analyzed for mobility quiescence.

### Analyzing functional Ca^2+^-images

Neuronal intensities were automatically analyzed via custom-written MATLAB scripts that are available at Github (https://github.com/ibusack/A-sleep-active-neuron-can-promote-survival-while-sleep-behavior-is-disturbed/blob/main/analysisfluorescentsignal.m). These codes extracted the regions of interest through an intensity threshold and acquired the intensities and position data. The extracted regions were set to be slightly bigger than the actual size of the neuron to allow for background intensity detection and background subtraction. The pan-neuronal activity was measured treating the head neurons as a combined signal. The RIS GCaMP activity in Figure 6 was determined by extracting the position data of the corresponding mKate2 signal. A larger region of interest was extracted and a fitting percentage of that was calculated as the signal. The extracted position data of the signal served to calculate the speed values. RIS, RIM and PLM activities were normalized by dividing GCaMP intensities through corresponding mKate2 intensities. In the optogenetic experiment, the baseline measurements were used for normalization.

### Mobility quiescence detection

First, speeds or image subtraction mean intensities were smoothed over 30 time points through the *smooth* function in MATLAB, which is a first-degree polynomial local regression model. The code is available at Github (https://github.com/ibusack/A-sleep-active-neuron-can-promote-survival-while-sleep-behavior-is-disturbed/blob/main/boutdetection.m). Smoothed speeds and image subtraction values were normalized between 0 and 1 for each individual animal with 0 being the lowest speed value and 1 the highest speed value the animal possessed. A mobility quiescence bout was counted when the normalized speed was below a specific percentage for a certain amount of time. For calcium imaging experiments with the 20× objective (Fig 6), the normalized speed had to be below 10% for at least 3min. The RIS activity experiment, which was filmed with a 10× objective, required a percentage threshold of 5% for a minimum of 4min. For DIC images filmed with the 20× objective, the normalized mean image subtraction values had to be below 15% for at least 3min to be counted as sleep time points. The code for detecting sleep bouts based on RIM tracking can be found at Github (https://github.com/ibusack/A-sleep-active-neuron-can-promote-survival-while-sleep-behavior-is-disturbed/blob/main/RIM_sleepboutdetection.m.)

### Mobility quiescence bout onset alignment

Speeds and GCaMP intensities were aligned to the start of mobility quiescence bouts, which had been detected by the mobility quiescence bout analysis. Mobility quiescence bouts were only counted for the alignment when the worm was mobile throughout 3min prior to the mobility quiescence bout onset. This alignment was carried out with a custom-written MATLAB script which is published on Github (https://github.com/ibusack/A-sleep-active-neuron-can-promote-survival-while-sleep-behavior-is-disturbed/blob/main/boutdetection.m).

### Neuronal inactivity detection

The neuronal activities were treated similarly as speed values for the mobility quiescence detection. Hence the boutdetection MATLAB script was utilized, which can be found on Github (https://github.com/ibusack/A-sleep-active-neuron-can-promote-survival-while-sleep-behavior-is-disturbed/blob/main/boutdetection.m). The nervous system was counted as inactive when it was below 20% of the smoothed and normalized activity for at least 3min.

### RIM neuron inactivity detection

RIM peaks were detected through a custom-written MATLAB script, which is available at Github (https://github.com/ibusack/A-sleep-active-neuron-can-promote-survival-while-sleep-behavior-is-disturbed/blob/main/analyze_RIM.m). First, the activity values were smoothed using the inbuilt *smooth* function over 5 time points. Then, peaks of a minimum prominence of 0.2 were extracted via the inbuilt function *islocalmax.* RIM was counted as inactive when there were no detected peaks for at least 5min.

### Induction of cellular stress by heat shock

The heat shock was carried out following a previously described protocol and utilizing a previously described heat shock device [17]. To summarize, young adult worms were imaged in 370×370×45μm microfluidic chambers and for a total of 4h. 5 to 12 worms were transferred from plates grown at 20°C to the microchamber. The worms were starved for 90min at 20°C before filming. During the entire duration of the movie, DIC images were acquired at a frequency of 0.1Hz. First, a baseline was recorded for 1h with the sample holder temperature set to 22°C and the heating lid set to 25°C. Then the heat shock was conducted with the sample holder temperature set to 37°C and the heating lid set to 39°C for 20 minutes. Finally, a recovery phase was recorded with the sample holder temperature set to 22°C and the heating lid set to 25°C for 2h 40min.

### Pumping assay

Arrested L1 worms were imaged in 110×110×10μm microfluidic chambers. DIC images were acquired with a frequency of 10 Hz for 1min. Pumps of the grinder were then counted manually with the help of a clicker.

### Survival assay in liquid culture

RIS impairment has been shown to confer a reduction of survival in both liquid culture and in microfluidic chambers [12, 23, 24, 36]. Worms were synchronized by bleaching [17]. 3-4 days before bleaching, 10-12 L4 stage worms were transferred onto NGM plates with a diameter of 6cm. 1-2 plates were bleached per strain and the day of bleaching was defined as day 0. The worms were kept in 1ml of M9 buffer in 2ml Eppendorf tubes in a rotator at 20°C for most survival assays. The following day was counted as day 1 of the lifespan. During the survival assay, the worms were kept in liquid culture in 1ml M9 buffer in 2ml Eppendorf tubes. This resulted in a population density of approximately 100 worms per 10μl. The tubes were placed in a rotator at 20°C, which rotated at 20rpm. For counting, 10μl drops of the liquid culture were pipetted onto fresh NGM plates for each strain respectively. The worms were allowed to recover for at least 2h after pipetting before counting. Counting was performed manually with the assistance of a clicker. The recovery number, i.e., the number of worms that developed into at least the L4 stage, was counted 3-5 days after pipetting. Around 50-100 worms were counted for each strain of each replicate on each time point.

### Survival assay in the OptoGenBox

For the long-term optogenetic survival assay worms were kept in microfluidic chambers (110×110×15μm) as described above. Between 27 and 43 worms were used for each strain per agarose slab each containing many microfluidic chambers. A maximum of one worm was placed per microfluidic chamber. Three replicates were performed per condition. We then calculated the average of all individuals of all three replicates combined. For the optogenetic activation 10μl of 10mM all-trans-retinal (ATR, Sigma Aldrich) dissolved in ethanol was added every 3-4 days. 20μl of 10μg/ml nystatin dissolved in DMSO was added 3-4 times throughout the lifespan to prevent fungal contamination. To prevent drying out of the agarose, 20μl of sterile water was added every 2 days. In the first week, the worms were counted every second day, then the survival was counted every day until all worms had died. A worm was counted as dead if a 2 min blue-light stimulus did not cause a visible behavioral response. For long-term optogenetic activation of RIS, worms were placed in the OptoGenBox [23] and continuously illuminated with 10mW at 20°C.

### Blue light survival assay

For the blue-light survival assays worms were imaged in 110×110×10μm microchambers. A blue light stimulus of 0.74mW/mm^2^ was given for 1s every 5s while simultaneously taking a GCaMP image. For the control, the same intensity stimulus was given for only 100ms through the camera exposure time.

### Imaging of growing worms

The images of growing plates were taken at a Leica M125 stereomicroscope. A Jenoptik ProgRes MF captured the images utilizing the corresponding software ProgResCapturePro 2.10.0.1.

### RNA sequencing data

RNA sequencing data is described in [24].

### Fitting

Origin software was used for the regression analyses. A linear regression was fitted to the data in Figure 6C and 6G. R^2^ values can be found in the corresponding figures. The exact functions are in Table 1.

### Quantification and Statistical Analysis

The experimenter was blind to the genotype information during the lifespan experiments. All other experiments relied on an automated analysis and hence were not blinded. The Wilcoxon signed rank test was utilized for the comparison of speeds and GCaMP intensities within the same strain. The Welch test was conducted to compare data from different strains. The survival assay experiments were statistically analyzed by utilizing the Fisher’s exact test at 50% survival of the wild type. Experiments with four or more genotypes were FDR corrected by the Benjamini-Hochberg procedure with a 5% false discovery rate. The graphs depict the mean and standard error of the mean. Box plots display individual data points, the interquartile range and the median. The whiskers represent the 10-90^th^ percentile. A table listing all exact p-values can be found in the supplemental information (Table S4).

## Supporting information

supplemental data sheets

## Acknowledgements

Some strains were provided by the CGC, which is funded by NIH Office of Research Infrastructure Programs (P40 OD010440). The CRISPR strains were generated by SunyBiotech based on our design. We thank Thomas Boulin for communicating the sodium permeability of *unc-58gf* and suggesting its use as a tool for neuronal activation. This work was supported by a European Research Council Starting Grant (ID: 637860, SLEEPCONTROL).

## Author contributions

I.B.: Data curation, Formal analysis, Investigation, Methodology, Project administration, Resources, Software, Visualization, Writing – review & editing. H.B.: Conceptualization, Funding acquisition, Investigation, Methodology, Project administration, Resources, Supervision, Writing – original draft.

## List of Legends

The supplementary file includes: Supplementary text

CRIPR sequences Supplementary Figures

Fig S1 to S10 Supplementary Tables

Table S1 fitting parameters

Table S2 strain list

Table S3 primer list

Table S4 p-values

Table S5 DEG genes from the transcriptome data

Movie legends for movies S1-S8

References for the supplementary information

## References

1. Campbell SS, Tobler I. Animal sleep: a review of sleep duration across phylogeny. Neurosci Biobehav Rev. 1984;8(3):269–300. Epub 1984/01/01. PubMed PMID: 6504414.

2. Diekelmann S, Born J. The memory function of sleep. Nature reviews Neuroscience. 2010;11(2):114–26. Epub 2010/01/05. doi: 10.1038/nrn2762. PubMed PMID: 20046194.

3. Krueger JM, Frank MG, Wisor JP, Roy S. Sleep function: Toward elucidating an enigma. Sleep medicine reviews. 2016;28:46–54. Epub 2015/10/09. doi: 10.1016/j.smrv.2015.08.005. PubMed PMID: 26447948; PubMed Central PMCID: PMC4769986.

4. Anafi RC, Kayser MS, Raizen DM. Exploring phylogeny to find the function of sleep. Nature reviews Neuroscience. 2019;20(2):109–16. Epub 2018/12/24. doi: 10.1038/s41583-018-0098-9. PubMed PMID: 30573905.

5. Hublin C, Partinen M, Koskenvuo M, Kaprio J. Sleep and mortality: a population-based 22-year follow-up study. Sleep. 2007;30(10):1245–53. Epub 2007/11/01. doi: 10.1093/sleep/30.10.1245. PubMed PMID: 17969458; PubMed Central PMCID: PMCPMC2266277.

6. Van Cauter E, Spiegel K, Tasali E, Leproult R. Metabolic consequences of sleep and sleep loss. Sleep Med. 2008;9 Suppl 1:S23-8. Epub 2008/12/17. doi: 10.1016/S1389-9457(08)70013-3. PubMed PMID: 18929315; PubMed Central PMCID: PMCPMC4444051.

7. Irwin MR. Why sleep is important for health: a psychoneuroimmunology perspective. Annu Rev Psychol. 2015;66:143–72. Epub 2014/07/26. doi: 10.1146/annurev-psych-010213-115205. PubMed PMID: 25061767; PubMed Central PMCID: PMCPMC4961463.

8. Saper CB, Fuller PM, Pedersen NP, Lu J, Scammell TE. Sleep state switching. Neuron. 2010;68(6):1023–42. Epub 2010/12/22. doi: 10.1016/j.neuron.2010.11.032. PubMed PMID: 21172606; PubMed Central PMCID: PMC3026325.

9. Bringmann H. Sleep-Active Neurons: Conserved Motors of Sleep. Genetics. 2018;208(4):1279–89. Epub 2018/04/06. doi: 10.1534/genetics.117.300521. PubMed PMID: 29618588; PubMed Central PMCID: PMC5887131.

10. Turek M, Lewandrowski I, Bringmann H. An AP2 transcription factor is required for a sleep-active neuron to induce sleep-like quiescence in C. elegans. Curr Biol. 2013;23(22):2215–23. Epub 2013/11/05. doi: 10.1016/j.cub.2013.09.028. PubMed PMID: 24184105.

11. Turek M, Besseling J, Spies JP, Konig S, Bringmann H. Sleep-active neuron specification and sleep induction require FLP-11 neuropeptides to systemically induce sleep. eLife. 2016;5. Epub 2016/03/08. doi: 10.7554/eLife.12499. PubMed PMID: 26949257; PubMed Central PMCID: PMC4805538.

12. Wu Y, Masurat F, Preis J, Bringmann H. Sleep Counteracts Aging Phenotypes to Survive Starvation-Induced Developmental Arrest in C. elegans. Curr Biol. 2018;28(22):3610–24 e8. Epub 2018/11/13. doi: 10.1016/j.cub.2018.10.009. PubMed PMID: 30416057; PubMed Central PMCID: PMCPMC6264389.

13. Skora S, Mende F, Zimmer M. Energy Scarcity Promotes a Brain-wide Sleep State Modulated by Insulin Signaling in C. elegans. Cell reports. 2018;22(4):953–66. Epub 2018/02/02. doi: 10.1016/j.celrep.2017.12.091. PubMed PMID: 29386137.

14. Gonzales DL, Zhou J, Fan B, Robinson JT. A microfluidic-induced C. elegans sleep state. Nat Commun. 2019;10(1):5035. Epub 2019/11/07. doi: 10.1038/s41467-019-13008-5. PubMed PMID: 31695031.

15. Steuer Costa W, Van der Auwera P, Glock C, Liewald JF, Bach M, Schuler C, et al. A GABAergic and peptidergic sleep neuron as a locomotion stop neuron with compartmentalized Ca2+ dynamics. Nat Commun. 2019;10(1):4095. Epub 2019/09/12. doi: 10.1038/s41467-019-12098-5. PubMed PMID: 31506439.

16. Grubbs JJ, Lopes LE, van der Linden AM, Raizen DM. A salt-induced kinase is required for the metabolic regulation of sleep. PLoS biology. 2020;18(4):e3000220. Epub 20200421. doi: 10.1371/journal.pbio.3000220. PubMed PMID: 32315298; PubMed Central PMCID: PMCPMC7173979.

17. Konietzka J, Fritz M, Spiri S, McWhirter R, Leha A, Palumbos S, et al. Epidermal Growth Factor Signaling Promotes Sleep through a Combined Series and Parallel Neural Circuit. Curr Biol. 2020;30(1):1–16 e3. Epub 2019/12/17. doi: 10.1016/j.cub.2019.10.048. PubMed PMID: 31839447.

18. Sinner MP, Masurat F, Ewbank JJ, Pujol N, Bringmann H. Innate Immunity Promotes Sleep through Epidermal Antimicrobial Peptides. Curr Biol. 2021;31(3):564–77 e12. Epub 2020/12/02. doi: 10.1016/j.cub.2020.10.076. PubMed PMID: 33259791.

19. Makino M, Ulzii E, Shirasaki R, Kim J, You YJ. Regulation of Satiety Quiescence by Neuropeptide Signaling in Caenorhabditis elegans. Front Neurosci. 2021;15:678590. Epub 2021/08/03. doi: 10.3389/fnins.2021.678590. PubMed PMID: 34335159; PubMed Central PMCID: PMCPMC8319666.

20. Nichols ALA, Eichler T, Latham R, Zimmer M. A global brain state underlies C. elegans sleep behavior. Science. 2017;356(6344). Epub 2017/06/24. doi: 10.1126/science.aam6851. PubMed PMID: 28642382.

21. Xu Y, Zhang L, Liu Y, Topalidou I, Hassinan C, Ailion M, et al. Dopamine receptor DOP-1 engages a sleep pathway to modulate swimming in C. elegans. iScience. 2021;24(4):102247. Epub 2021/04/03. doi: 10.1016/j.isci.2021.102247. PubMed PMID: 33796839; PubMed Central PMCID: PMCPMC7995527.

22. Maluck E, Busack I, Besseling J, Masurat F, Turek M, Busch KE, et al. A wake-active locomotion circuit depolarizes a sleep-active neuron to switch on sleep. PLoS biology. 2020;18(2):e3000361. Epub 2020/02/23. doi: 10.1371/journal.pbio.3000361. PubMed PMID: 32078631.

23. Busack I, Jordan F, Sapir P, Bringmann H. The OptoGenBox - a device for long-term optogenetics in C. elegans. J Neurogenet. 2020:1–9. Epub 2020/06/17. doi: 10.1080/01677063.2020.1776709. PubMed PMID: 32543249.

24. Koutsoumparis A, Welp LM, Wulf A, Urlaub H, Meierhofer D, Börno S, et al. Sleep neuron depolarization promotes protective gene expression changes and FOXO activation. Current Biology. 2022. doi: https://doi.org/10.1016/j.cub.2022.04.012.

25. Kenyon CJ. The genetics of ageing. Nature. 2010;464(7288):504-12. Epub 2010/03/26. doi: 10.1038/nature08980. PubMed PMID: 20336132.

26. Baugh LR, Hu PJ. Starvation Responses Throughout the Caenorhabditis elegans Life Cycle. Genetics. 2020;216(4):837–78. Epub 2020/12/04. doi: 10.1534/genetics.120.303565. PubMed PMID: 33268389; PubMed Central PMCID: PMCPMC7768255.

27. Siegel JM. Sleep viewed as a state of adaptive inactivity. Nature reviews Neuroscience. 2009;10(10):747–53. Epub 2009/08/06. doi: 10.1038/nrn2697. PubMed PMID: 19654581.

28. Schmidt MH. The energy allocation function of sleep: a unifying theory of sleep, torpor, and continuous wakefulness. Neurosci Biobehav Rev. 2014;47:122–53. Epub 2014/08/15. doi: 10.1016/j.neubiorev.2014.08.001. PubMed PMID: 25117535.

29. Krause AJ, Simon EB, Mander BA, Greer SM, Saletin JM, Goldstein-Piekarski AN, et al. The sleep-deprived human brain. Nature reviews Neuroscience. 2017;18(7):404–18. Epub 2017/05/19. doi: 10.1038/nrn.2017.55. PubMed PMID: 28515433.

30. Klinzing JG, Niethard N, Born J. Mechanisms of systems memory consolidation during sleep. Nat Neurosci. 2019;22(10):1598–610. Epub 2019/08/28. doi: 10.1038/s41593-019-0467-3. PubMed PMID: 31451802.

31. Taylor SR, Santpere G, Weinreb A, Barrett A, Reilly MB, Xu C, et al. Molecular topography of an entire nervous system. Cell. 2021. Epub 2021/07/09. doi: 10.1016/j.cell.2021.06.023. PubMed PMID: 34237253.

32. Ben Soussia I, El Mouridi S, Kang D, Leclercq-Blondel A, Khoubza L, Tardy P, et al. Mutation of a single residue promotes gating of vertebrate and invertebrate two-pore domain potassium channels. Nat Commun. 2019;10(1):787. Epub 2019/02/17. doi: 10.1038/s41467-019-08710-3. PubMed PMID: 30770809; PubMed Central PMCID: PMCPMC6377628.

33. Kunkel MT, Johnstone DB, Thomas JH, Salkoff L. Mutants of a temperature-sensitive two-P domain potassium channel. J Neurosci. 2000;20(20):7517–24. doi: 10.1523/JNEUROSCI.20-20-07517.2000. PubMed PMID: 11027209; PubMed Central PMCID: PMCPMC6772866.

34. Silva-Garcia CG, Lanjuin A, Heintz C, Dutta S, Clark NM, Mair WB. Single-Copy Knock-In Loci for Defined Gene Expression in Caenorhabditis elegans. G3 (Bethesda). 2019;9(7):2195-8. Epub 2019/05/09. doi: 10.1534/g3.119.400314. PubMed PMID: 31064766; PubMed Central PMCID: PMCPMC6643881.

35. Bringmann H. Genetic sleep deprivation: using sleep mutants to study sleep functions. EMBO Rep. 2019;20(3). Epub 2019/02/26. doi: 10.15252/embr.201846807. PubMed PMID: 30804011; PubMed Central PMCID: PMCPMC6399599.

36. Roux AE, Langhans K, Huynh W, Kenyon C. Reversible Age-Related Phenotypes Induced during Larval Quiescence in C. elegans. Cell Metab. 2016;23(6):1113–26. Epub 2016/06/16. doi: 10.1016/j.cmet.2016.05.024. PubMed PMID: 27304510.

37. Bringmann H. Agarose hydrogel microcompartments for imaging sleep- and wake-like behavior and nervous system development in Caenorhabditis elegans larvae. J Neurosci Methods. 2011;201(1):78–88. Epub 2011/08/02. doi: S0165-0270(11)00418-3 [pii]10.1016/j.jneumeth.2011.07.013. PubMed PMID: 21801751.

38. Turek M, Besseling J, Bringmann H. Agarose Microchambers for Long-term Calcium Imaging of Caenorhabditis elegans. Journal of visualized experiments: JoVE. 2015;(100):e52742. Epub 2015/07/02. doi: 10.3791/52742. PubMed PMID: 26132740; PubMed Central PMCID: PMC4544933.

39. Van Buskirk C, Sternberg PW. Epidermal growth factor signaling induces behavioral quiescence in Caenorhabditis elegans. Nat Neurosci. 2007;10(10):1300–7. Epub 2007/09/25. doi: 10.1038/nn1981. PubMed PMID: 17891142.

40. Hill AJ, Mansfield R, Lopez JM, Raizen DM, Van Buskirk C. Cellular stress induces a protective sleep-like state in C. elegans. Curr Biol. 2014;24(20):2399–405. Epub 2014/09/30. doi: 10.1016/j.cub.2014.08.040. PubMed PMID: 25264259; PubMed Central PMCID: PMC4254280.

41. Nelson MD, Lee KH, Churgin MA, Hill AJ, Van Buskirk C, Fang-Yen C, et al. FMRFamide-like FLP-13 neuropeptides promote quiescence following heat stress in Caenorhabditis elegans. Curr Biol. 2014;24(20):2406–10. Epub 2014/09/30. doi: 10.1016/j.cub.2014.08.037. PubMed PMID: 25264253; PubMed Central PMCID: PMC4254296.

42. Iannacone MJ, Beets I, Lopes LE, Churgin MA, Fang-Yen C, Nelson MD, et al. The RFamide receptor DMSR-1 regulates stress-induced sleep in C. elegans. eLife. 2017;6. Epub 2017/01/18. doi: 10.7554/eLife.19837. PubMed PMID: 28094002; PubMed Central PMCID: PMCPMC5241116.

43. Cassada RC, Russell RL. The dauerlarva, a post-embryonic developmental variant of the nematode Caenorhabditis elegans. Dev Biol. 1975;46(2):326–42. Epub 1975/10/01. doi: 0012-1606(75)90109-8 [pii]. PubMed PMID: 1183723.

44. Raizen DM, Zimmerman JE, Maycock MH, Ta UD, You YJ, Sundaram MV, et al. Lethargus is a Caenorhabditis elegans sleep-like state. Nature. 2008;451(7178):569-72. Epub 2008/01/11. doi: nature06535 [pii]10.1038/nature06535. PubMed PMID: 18185515.

45. Yemini E, Lin A, Nejatbakhsh A, Varol E, Sun R, Mena GE, et al. NeuroPAL: A Multicolor Atlas for Whole-Brain Neuronal Identification in C. elegans. Cell. 2021;184(1):272–88 e11. Epub 2020/12/31. doi: 10.1016/j.cell.2020.12.012. PubMed PMID: 33378642.

46. Piggott BJ, Liu J, Feng Z, Wescott SA, Xu XZ. The neural circuits and synaptic mechanisms underlying motor initiation in C. elegans. Cell. 2011;147(4):922–33. Epub 2011/11/15. doi: 10.1016/j.cell.2011.08.053. PubMed PMID: 22078887; PubMed Central PMCID: PMC3233480.

47. Li W, Kang L, Piggott BJ, Feng Z, Xu XZ. The neural circuits and sensory channels mediating harsh touch sensation in Caenorhabditis elegans. Nat Commun. 2011;2:315. Epub 2011/05/19. doi: 10.1038/ncomms1308. PubMed PMID: 21587232; PubMed Central PMCID: PMC3098610.

48. McCormick DA, Bal T. Sensory gating mechanisms of the thalamus. Curr Opin Neurobiol. 1994;4(4):550–6. doi: 10.1016/0959-4388(94)90056-6. PubMed PMID: 7812144.

49. Schwarz J, Lewandrowski I, Bringmann H. Reduced activity of a sensory neuron during a sleep-like state in Caenorhabditis elegans. Curr Biol. 2011;21(24):R983-4. Epub 2011/12/24. doi: S0960-9822(11)01207-3[pii]10.1016/j.cub.2011.10.046. PubMed PMID: 22192827.

50. Schwarz J, Bringmann H. Reduced sleep-like quiescence in both hyperactive and hypoactive mutants of the Galphaq Gene egl-30 during lethargus in Caenorhabditis elegans. PLoS One. 2013;8(9):e75853. Epub 2013/09/28. doi: 10.1371/journal.pone.0075853. PubMed PMID: 24073282; PubMed Central PMCID: PMC3779211.

51. Cho JY, Sternberg PW. Multilevel modulation of a sensory motor circuit during C. elegans sleep and arousal. Cell. 2014;156(1-2):249–60. Epub 2014/01/21. doi: 10.1016/j.cell.2013.11.036. PubMed PMID: 24439380; PubMed Central PMCID: PMC3962823.

52. Schwarz J, Spies JP, Bringmann H. Reduced muscle contraction and a relaxed posture during sleep-like Lethargus. Worm. 2012;1(1):12–4. Epub 2012/01/01. doi: 10.4161/worm.19499. PubMed PMID: 24058817; PubMed Central PMCID: PMC3670164.

53. Iwanir S, Tramm N, Nagy S, Wright C, Ish D, Biron D. The microarchitecture of C. elegans behavior during lethargus: homeostatic bout dynamics, a typical body posture, and regulation by a central neuron. sleep. 2013;36(3):385–95. Epub 2013/03/02. doi: 10.5665/sleep.2456. PubMed PMID: 23449971; PubMed Central PMCID: PMC3571756.

54. Tramm N, Oppenheimer N, Nagy S, Efrati E, Biron D. Why do sleeping nematodes adopt a hockey-stick-like posture? PLoS One. 2014;9(7):e101162. Epub 2014/07/16. doi: 10.1371/journal.pone.0101162. PubMed PMID: 25025212; PubMed Central PMCID: PMC4099128.

55. Schwartz HT. A protocol describing pharynx counts and a review of other assays of apoptotic cell death in the nematode worm Caenorhabditis elegans. Nature protocols. 2007;2(3):705–14. doi: 10.1038/nprot.2007.93. PubMed PMID: 17406633.

56. Song BM, Avery L. The pharynx of the nematode C. elegans: A model system for the study of motor control. Worm. 2013;2(1):e21833. doi: 10.4161/worm.21833. PubMed PMID: 24058858; PubMed Central PMCID: PMCPMC3670459.

57. Urmersbach B, Besseling J, Spies JP, Bringmann H. Automated analysis of sleep control via a single neuron active at sleep onset in C. elegans. Genesis. 2016;54(4):212–9. Epub 2016/02/03. doi: 10.1002/dvg.22924. PubMed PMID: 26833569.

58. Bennett HL, Khoruzhik Y, Hayden D, Huang H, Sanders J, Walsh MB, et al. Normal sleep bouts are not essential for C. elegans survival and FoxO is important for compensatory changes in sleep. BMC neuroscience. 2018;19(1):10. Epub 2018/03/11. doi: 10.1186/s12868-018-0408-1. PubMed PMID: 29523076.

59. White JG, Southgate E, Thomson JN, Brenner S. The structure of the nervous system of the nematode Caenorhabditis elegans. Philos Trans R Soc Lond B Biol Sci. 1986;314(1165):1-340. Epub 1986/11/12. PubMed PMID: 22462104.

60. Schafer WR, Kenyon CJ. A calcium-channel homologue required for adaptation to dopamine and serotonin in Caenorhabditis elegans. Nature. 1995;375(6526):73-8. Epub 1995/05/04. doi: 10.1038/375073a0. PubMed PMID: 7723846.

61. Lee RY, Lobel L, Hengartner M, Horvitz HR, Avery L. Mutations in the alpha1 subunit of an L-type voltage-activated Ca2+ channel cause myotonia in Caenorhabditis elegans. EMBO J. 1997;16(20):6066–76. Epub 1997/10/08. doi: 10.1093/emboj/16.20.6066. PubMed PMID: 9321386.

62. Mathews EA, Garcia E, Santi CM, Mullen GP, Thacker C, Moerman DG, et al. Critical residues of the Caenorhabditis elegans unc-2 voltage-gated calcium channel that affect behavioral and physiological properties. J Neurosci. 2003;23(16):6537–45. doi: 10.1523/JNEUROSCI.23-16-06537.2003. PubMed PMID: 12878695; PubMed Central PMCID: PMCPMC6740628.

63. Laine V, Frokjaer-Jensen C, Couchoux H, Jospin M. The alpha1 subunit EGL-19, the alpha2/delta subunit UNC-36, and the beta subunit CCB-1 underlie voltage-dependent calcium currents in Caenorhabditis elegans striated muscle. The Journal of biological chemistry. 2011;286(42):36180–7. Epub 2011/09/01. doi: 10.1074/jbc.M111.256149. PubMed PMID: 21878625; PubMed Central PMCID: PMC3196126.

64. Schuler C, Fischer E, Shaltiel L, Steuer Costa W, Gottschalk A. Arrhythmogenic effects of mutated L-type Ca 2+-channels on an optogenetically paced muscular pump in Caenorhabditis elegans. Sci Rep. 2015;5:14427. Epub 20150924. doi: 10.1038/srep14427. PubMed PMID: 26399900; PubMed Central PMCID: PMCPMC4585839.

65. Huang YC, Pirri JK, Rayes D, Gao S, Mulcahy B, Grant J, et al. Gain-of-function mutations in the UNC-2/CaV2alpha channel lead to excitation-dominant synaptic transmission in Caenorhabditis elegans. eLife. 2019;8. Epub 20190805. doi: 10.7554/eLife.45905. PubMed PMID: 31364988; PubMed Central PMCID: PMCPMC6713474.

66. Lin JY, Knutsen PM, Muller A, Kleinfeld D, Tsien RY. ReaChR: a red-shifted variant of channelrhodopsin enables deep transcranial optogenetic excitation. Nat Neurosci. 2013;16(10):1499–508. Epub 2013/09/03. doi: 10.1038/nn.3502. PubMed PMID: 23995068; PubMed Central PMCID: PMC3793847.

67. Driver RJ, Lamb AL, Wyner AJ, Raizen DM. DAF-16/FOXO Regulates Homeostasis of Essential Sleep-like Behavior during Larval Transitions in C. elegans. Curr Biol. 2013;23(6):501–6. Epub 2013/03/13. doi: 10.1016/j.cub.2013.02.009. PubMed PMID: 23477722.

68. Choi S, Chatzigeorgiou M, Taylor KP, Schafer WR, Kaplan JM. Analysis of NPR-1 reveals a circuit mechanism for behavioral quiescence in C. elegans. Neuron. 2013;78(5):869–80. Epub 2013/06/15. doi: 10.1016/j.neuron.2013.04.002. PubMed PMID: 23764289; PubMed Central PMCID: PMC3683153.

69. Spies J, Bringmann H. Automated detection and manipulation of sleep in C. elegans reveals depolarization of a sleep-active neuron during mechanical stimulation-induced sleep deprivation. Sci Rep. 2018;8(1):9732. Epub 2018/06/29. doi: 10.1038/s41598-018-28095-5. PubMed PMID: 29950594.

70. Stroustrup N, Ulmschneider BE, Nash ZM, Lopez-Moyado IF, Apfeld J, Fontana W. The Caenorhabditis elegans Lifespan Machine. Nat Methods. 2013;10(7):665–70. Epub 20130512. doi: 10.1038/nmeth.2475. PubMed PMID: 23666410; PubMed Central PMCID: PMCPMC3865717.

71. Laurent P, Soltesz Z, Nelson GM, Chen C, Arellano-Carbajal F, Levy E, et al. Decoding a neural circuit controlling global animal state in C. elegans. eLife. 2015;4. Epub 20150311. doi: 10.7554/eLife.04241. PubMed PMID: 25760081; PubMed Central PMCID: PMCPMC4440410.

72. Chun L, Gong J, Yuan F, Zhang B, Liu H, Zheng T, et al. Metabotropic GABA signalling modulates longevity in C. elegans. Nat Commun. 2015;6:8828. Epub 20151105. doi: 10.1038/ncomms9828. PubMed PMID: 26537867; PubMed Central PMCID: PMCPMC4667614.

73. Nagy S, Tramm N, Sanders J, Iwanir S, Shirley IA, Levine E, et al. Homeostasis in C. elegans sleep is characterized by two behaviorally and genetically distinct mechanisms. eLife. 2014;3:e04380. Epub 2014/12/05. doi: 10.7554/eLife.04380. PubMed PMID: 25474127; PubMed Central PMCID: PMC4273442.

74. Alam MA, Kumar S, McGinty D, Alam MN, Szymusiak R. Neuronal activity in the preoptic hypothalamus during sleep deprivation and recovery sleep. J Neurophysiol. 2014;111(2):287–99. Epub 2013/11/01. doi: 10.1152/jn.00504.2013. PubMed PMID: 24174649; PubMed Central PMCID: PMC3921380.

75. Brenner S. The genetics of Caenorhabditis elegans. Genetics. 1974;77(1):71–94. Epub 1974/05/01. PubMed PMID: 4366476.

76. Stiernagle T. Maintenance of C. elegans. WormBook. 2006:1–11. Epub 2007/12/01. doi: 10.1895/wormbook.1.101.1. PubMed PMID: 18050451.

77. Ahringer A. Reverse genetics. WormBook, ed The C elegans Research Community. 2006. doi: doi/10.1895/wormbook.1.47.1.

78. Redemann S, Schloissnig S, Ernst S, Pozniakowsky A, Ayloo S, Hyman AA, et al. Codon adaptation-based control of protein expression in C. elegans. Nat Methods. 2011;8(3):250–2. Epub 2011/02/01. doi: nmeth.1565 [pii]10.1038/nmeth.1565. PubMed PMID: 21278743.

79. Nagy S, Raizen DM, Biron D. Measurements of behavioral quiescence in Caenorhabditis elegans. Methods. 2014. Epub 2014/03/20. doi: 10.1016/j.ymeth.2014.03.009. PubMed PMID: 24642199.

