## supplemental information for "A sleep-active neuron can promote survival while sleep behavior is disturbed"

### Supplementary Text

CRISPR sequences

PHX1433: *flp-11(syb1433[flp-11-SL2-egl-23cDNA(A383V)-linker-mKate2])X*

>flp-11b-SL2(gpd-2)-egl-23(n601)-linker(GSGSGSGSG)-mKate2(two introns)
ctattgtagTGCGGAGAAACGTGCCATGCGGAACGCCTTGGTTCGATTTGGAAGAGCTAGTGGTGGAATGAGAAA
TGCTCTCGTTAGATTCGGAAAGAGGTCTCCATTGGACGAGGAAGACTTTGCTCCAGAGAGCCCACTCCAGGGAAA
ACGGAACGGTGCCCCACAACCATTTGgtaagttgtcttaaaatttttcttccgctttttgcctttgcttcatgtg
tcgtttattttgctttgcagttcgctttggccgatccggtcaactcgaccacatgcacgaccttttgtcgactct
tcagAAGCTCAAGTTCGCCAACAACAAGTAATGACCGAGGACGACCGTCTTCTGCTCGAACAACTCCTGCGACGA
ATTCATCATTAAgctgtctcatcctactttcacctagttaactgcttgtcttaaaatctatgcttctctttagta
tctaaaattttcctagaagcttacaagtatataaatggtctcttctcaataaaggttgtatatttattcatctta
ttgaatctgccatttcctcgtttttgcgagtttatataccttccaatttctttctattgtattttcaacttcta
attttaattcagggaaactgcttcaacgcatcATGAAGCTCACGTTGAAGAAATCCGTATTCTCAAGGGATAAAC
ATATCTTGCAAAAGGCGACACCACTATTCGTTCACTTTCTAATGATAGTAAGTGTGGGTGCCTACGCAATATTTG
GAGCACTTGTAATGAGAAGTCTTGAATCGAGAACTGTCACAAGTATTGAAAAGAAGACGGATGTTCACAGAAGAC
ATGTTAATTTGACTAATTTTCAACATCCACCAACTCCAATCACATTAGAGCAAAGACATCGACGGAGACGCAGAC
ATAATGAGACAGCTTTGGAAGACCATTTGTCTGAAAAATTATCCAGAGAAAAACGTGCAGCAGCGCATATCATGA
GAAGTCGAAAATGCGTTATCAGTGTGATAAAGAAAATGTCAAGCATGGAATGTTCATTTGACACTCTCGACGAGA
AGCTCGTAAAAGCACTCGATGAATGTTACCACGTGGCAGTTGAACATAATACTCATGTGAATCATGTACTTTTCA
CGAATAGTAAGGAAGAAGTGGAGTCAGTTGGGGAAGAAGCAGAAGAAGATGTTTCCGAATGGTCATTTATGGACT
CGTTGTTGTTTGCATTCACTGTTATTACGACGATTGGATACGGAAACGTGGCACCTCGAACATTTGGTGGCCGTC
TATTTGTCATTGGTTATGGTCTAATTGGTATTCCGTTTACACTGCTGGCAATTGCAGATCTCGGAAAATTCATAT
CAGAAATGATGGTGGAGGCGAAAAGTTTTTGTAGGAAAACCTGGAAAAAACTCAAAAAAGCGTGGAACCCGAATT
TCATTCGCGCAAAGGATCTTTCAAATACGGATATTGAGGAGAAAATATTGGATAATGAGAAAATCGAAAATGAGC
CGGAAACTTCAGAAGTATCAGAAGAAGAAGACGATTTGACAGAGACAGAAGCCACGTCACTTTTCATTTTATTTT
TGGTTTATATCGCATTTGGAGGGTTCATGTTAGCTGCTTATGAACCTGATATGGACTTTTTCAAAGCGGTCTACT
TTAATTTTGTGACATTGACATCAATTGGTCTGGGAGATATTGTACCGAGAAGTGAAACCTATATGCTCATTACAA
TAGTCTACATTGCAATTGGTCTTGCTCTTACCACAATTGCCATTGAAATCGCCGTAGATGCATTGAAGAAGCTGC
ATTATTTTGGAAGAAAAATCGAAAATGTTGGAAATGTGGCTATATGGTTTGGAGGAAAGAAAATTACAATGAAAG
CACTGGTCAAAAACCTCGGTGACCAGTTCAATCTTCCAACTACCGTAGTCAAGAATTTGAATTTGGATCATTTTG
TGGATCAAGCGATTAAAGTGGAGGAAGGAGAGATTGAGACACTCAGACCGCCGCCTTATGAGCCACCGTCTGATC
GATTTGAAGCTGAATTCGCTGATGAACCAGAATGTGAATGGATCCGTGATCCGACTCCAACTCCACCCCCATCAC
CTCAACCGGTTTATCGTCTTCCATCTCCAAAACCAGTTACACCGGAACCTCTACCAAGTCCAACAATAACTGATG
TATCACTTGCGATTGCAACACCATCACCTGAAGAATCGGATGATGATCAAGAACTAATTCTCCCATCACCTGAAC
CGAGTCCAGTTCGAGAGCCAACTCCACCACCGCCTCCACGGGAGCCAACACCTCGTGAGCCAACTCCTGAGCCAG
AGCCAGTTCGAGAGCCGACGCCTCCACCTCCGCCACCTGCCAAGCCCCGTCCACTGACTGCCGCTGAAATCGCGG
CTCAAAAACGCAAAGCGTACAGCGAAGAAGCATGGCGTCGATACCAAGAATACCAGAAACAATGGAAGAAGTTCC
GTCAAACTCAGAAAACTCCCGCACCATCTGGAGCCTCTACATCAGGGGCATCAACATCAAAACCGTCTGGAACAT
CACCGGAAAGTGGAGCCGGAGTATGTACTGGACCATCAACAAGGAGTCAATCAATAACATCAGTTGCATCTGGAA
AGACATCAAGAAGTGCAACACCGGAAAGCAAGAAATCATCACATTTGAGTGGCTCATCGAGAAGAGAAAGCGGTG
GAAAAGGATCCGGATCCGGATCCGGATCCGGAATGTCCGAGCTCATCAAGGAGAACATGCACATGAAGCTCTACA
TGGAGGGAACCGTCAACAACCACCACTTCAAGTGCACCTCCGAGGGAGAGGGAAAGCCATACGAGGGAACCCAAA
CCATGCGTATCAAGgtaagtttaaacatatatatactaactaaccctgattatttaaattttcagGCCGTCGAGG
GAGGACCACTCCCATTCGCCTTCGACATCCTCGCCACCTCCTTCATGTACGGATCCAAGACCTTCATCAACCACA
CCCAAGGAATCCCAGACTTCTTCAAGCAATCCTTCCCAGAGGGATTCACCTGGGAGCGTGTCACCACCTACGAGG
ACGGAGGAGTCCTCACCGCCACCCAAGACACCTCCCTCCAAGACGGATGCCTCATCTACAACGTCAAGATCCGTG
GAGTCAACTTCCCATCCAACGGACCAGTCATGCAAAAGAAGACCCTCGGATGGGAGGCCTCCACCGAGACCCTCT
ACCCAGCCGACGGAGGACTCGAGGGACGTGCCGACATGGCCCTCAAGCTCGTCGGAGGAGGACACCTCATCTGCA
ACCTCAAGgtaagtttaaacatgattttactaactaactaatctgatttaaattttcagACCACCTACCGTTCCA
AGAAGCCAGCCAAGAACCTCAAGATGCCAGGAGTCTACTACGTCGACCGTCGTCTCGAGCGTATCAAGGAGGCCG
ACAAGGAGACCTACGTCGAGCAACACGAGGTCGCCGTCGCCCGTTACTGCGACCTCCCATCCAAGCTCGGACACC
GTTAAaaatcatatgtttttctctctcacactctcttttttc

PHX1445: *aptf-1(gk794)II; flp-11(syb1445[flp-11-SL2-unc-58(L428F)-linker-mKate2])X*

>flp-11b-SL2(gpd-2)-unc-58(e665)-linker(GSGSGSGSG)-mKate2(two introns)
TGGAAGAGCTAGTGGTGGAATGAGAAATGCTCTCGTTAGATTCGGAAAGAGGTCTCCATTGGACGAGGAAGACTT
TGCTCCAGAGAGCCCACTCCAGGGAAAACGGAACGGTGCCCCACAACCATTTGgtaagttgtcttaaattttttc
ttccgctttttgcctttgcttcatgtgtcgtttatttttgctttgcagttcgctttggccgatccggtcaactcga
ccacatgcacgaccttttgtcgactcttcagAAGCTCAAGTTCGCCAACAACAAGTAATGACCGAGGACGACCGT
CTTCTGCTCGAACAACTCCTGCGACGAATTCATCATTAAgctgtctcatcctactttcacctagttaactgcttg
tcttaaaatctatgcttctctttagtatctaaaattttcctagaagcttacaagtatataaatggtctcttctca
ataaaggttgtatatttattcatcttattgaatctgccatttcctcgtttttgcgagtttatataccttccaatt
ttctttctattgtattttcaacttctaattttaattcagggaaactgcttcaacgcatcATGGCTCCACTGACTG
TGAAAAGCTCACCTCCAAAAAAGGCAAAAGGAATATCAAAATTTCGGAGAAAAAAGAAGCAGCCACCACCAGACT
CAACCGTATTCGTCGCATGGGCACTCCGAAGTGTCCGAGAGTCTCTGATCCAAGTTGATCCATTGGCCGCTGCAC
TTGCACATCAGGCTCGAAAGACAAATAGTGTGCCAGCTGTCTCGAGAACTCCACTGCTTCTACAGTTCACTCCTT
TCGGACCACCTCTCAGTGCGTATCATGTGACAGCTCGGTGGGAAGGTGCAAATATCAATTCACAATCAGCATTGC
TCGATGCAGATGATGGAGCTACAGTTATCACAGATACCATCAAAGATGACCAAGATGATAAAGAACCAAAAAGCT
GCCCGCAACAGACTGTCAAATACATCAAAATACTTACACCTCACGTGATCTTGGTGTCAGTGTTAATTGGATATT
TATGCTTGGGAGCTTGGATACTCATGTTACTGGAAACAAGGACGGAACTTCTTGCCAGATCCAAAAAACTTGTCA
GGTTAACAAATTTGATGTCAAACTTCACTGCCGAAAGTTGGAAGATGCTCAATAATGCTCAACACGGGGTTAGTA
ATATGGATGAAGGTGAATGGGCTGCAACATTTCGAGAATGGATGGTACGAGTATCAGAAACAGTGGACGATAGGA
GACCTATACGACGTGAATTAAACCGGCCTGATGACTTATCAAATATGCATAATAAATGGACATTTCCAACTGCAA
TATTATATGTTCTCACTGTGTTAACTACTTGCGGTTATGGAGAAGTATCTGTCGACACAGACGTCGGAAAGGTTT
TCTCAGTAGCATTCGCGCTTGTTGGTATACCACTTATGTTCATAACAGCTGCCGATATTGGTAAATTTTTATCTG
AAACATTACTCCAGTTTGTGAGCTTTTGGAATCGAAGTGTCCGAAAAGTGAAGCAATGGATGAGTCGTATTCGTC
ACGGCAGGAGAAAGTCATTACAATCAACGGGTGGTCCCAACGATACTCTCGATATTCTTGGTGTCGACGGAACTG
AAGAGAAACTTTGGTTCCCAATAGGTGCATATGTATCATGTATTTGCATATATTGCTCAATTGGGTCTGCCATGT
TTATCACATGGGAAAGAACTTGGTCTTTCATTCATGCGTTTCATTTTGGTTTCAATTTGATTGTAACAGTCGGAC
TCGGAGATATCGTTGTGACTGATTACATATTTTTATCACTTATCGTTGCATTTGTGATAGTTGGTTTTCCGTAG
TGACCATGTGCGTGGATCTTGCGTCCACACATCTCAAGGCGTACTTCACCAGAATTCACTACTTTGGTCGAGCAA
AACGATTCTTAGGAATGAGTGAGGAACTCAAAGAAATCGTTGCTTTACTGGGGGCGATGCGACGGAAAAAAGGCG
GTAAAGTTACATGGAATGATGTGCGAGACTTCCTGGATAACGAACTCCGCGATCGACCTTTTGAACCTCATGAGC
TTCTGATGAAGCTCAGATTTATTGACGAAACATCTTCTGGAATGTCTACAATCCGTCACAATTCCTTCCAGTCAG
ATTTTTTCCGAGAATCAGAGTACATCCGAAGAGTGGCTGCGCTGAGGCCAGAACAGCCAGCATATTTGGGATCCG
GATCCGGATCCGGATCCGGAATGTCCGAGCTCATCAAGGAGAACATGCACATGAAGCTCTACATGGAGGGAACCG
TCAACAACCACCACTTCAAGTGCACCTCCGAGGGAGAGGGAAAGCCATACGAGGGAACCCAAACCATGCGTATCA
AGgtaagtttaaacatatatatactaactaaccctgattatttaaattttcagGCCGTCGAGGGAGGACCACTCC
CATTCGCCTTCGACATCCTCGCCACCTCCTTCATGTACGGATCCAAGACCTTCATCAACCACACCCAAGGAATCC
CAGACTTCTTCAAGCAATCCTTCCCAGAGGGATTCACCTGGGAGCGTGTCACCACCTACGAGGACGGAGGAGTCC
TCACCGCCACCCAAGACACCTCCCTCCAAGACGGATGCCTCATCTACAACGTCAAGATCCGTGGAGTCAACTTCC
CATCCAACGGACCAGTCATGCAAAAGAAGACCCTCGGATGGGAGGCCTCCACCGAGACCCTCTACCCAGCCGACG
GAGGACTCGAGGGACGTGCCGACATGGCCCTCAAGCTCGTCGGAGGAGGACACCTCATCTGCAACCTCAAGgtaa
gtttaaacatgattttactaactaactaatctgatttaaattttcagACCACCTACCGTTCCAAGAAGCCAGCCA
AGAACCTCAAGATGCCAGGAGTCTACTACGTCGACCGTCGTCTCGAGCGTATCAAGGAGGCCGACAAGGAGACCT
ACGTCGAGCAACACGAGGTCGCCGTCGCCCGTTACTGCGACCTCCCATCCAAGCTCGGACACCGTTAAaaatcat
atgtttttctctctcacactctcttttttc

PHX1464: *flp-11(syb1464[flp-11-SL2-egl-23cDNA(L229N)-linker-mKate2])X*

>flp-11b-SL2(gpd-2)-egl-23(L229N)-linker(GSGSGSGSG)-mKate2(two introns)
ctattgtagTGCGGAGAAACGTGCCATGCGGAACGCCTTGGTTCGATTTGGAAGAGCTAGTGGTGGAATGAGAAA
TGCTCTCGTTAGATTCGGAAAGAGGTCTCCATTGGACGAGGAAGACTTTGCTCCAGAGAGCCCACTCCAGGGAAA
ACGGAACGGTGCCCCACAACCATTTGgtaagttgtcttaaattttttcttccgctttttgcctttgcttcatgtg
tcgtttatttttgctttgcagttcgctttggccgatccggtcaactcgaccacatgcacgaccttttgtcgactct
tcagAAGCTCAAGTTCGCCAACAACAAGTAATGACCGAGGACGACCGTCTTCTGCTCGAACAACTCCTGCGACGA
ATTCATCATTAAgctgtctcatcctactttcacctagttaactgcttgtcttaaaatctatgcttctctttagta
tctaaaattttcctagaagcttacaagtatataaatggtctcttctcaataaaggttgtatatttattcatctta
ttgaatctgccatttcctcgtttttgcgagtttatataccttccaattttctttctattgtattttcaacttcta
attttaattcagggaaactgcttcaacgcatcATGAAGCTCACGTTGAAGAAATCCGTATTCTCAAGGGATAAAC
ATATCTTGCAAAAGGCGACACCACTATTCGTTCACTTTCTAATGATAGTAAGTGTGGGTGCCTACGCAATATTTG
GAGCACTTGTAATGAGAAGTCTTGAATCGAGAACTGTCACAAGTATTGAAAAGAAGACGGATGTTCACAGAAGAC
ATGTTAATTTGACTAATTTTCAACATCCACCAACTCCAATCACATTAGAGCAAAGACATCGACGGAGACGCAGAC
ATAATGAGACAGCTTTGGAAGACCATTTGTCTGAAAAATTATCCAGAGAAAAACGTGCAGCAGCGCATATCATGA

GAAGTCGAAAATGCGTTATCAGTGTGATAAAGAAAATGTCAAGCATGGAATGTTCATTTGACACTCTCGACGAGA
AGCTCGTAAAAGCACTCGATGAATGTTACCACGTGGCAGTTGAACATAATACTCATGTGAATCATGTACTTTTCA
CGAATAGTAAGGAAGAAGTGGAGTCAGTTGGGGAAGAAGCAGAAGAAGATGTTTCCGAATGGTCATTTATGGACT
CGTTGTTGTTTGCATTCACTGTTATTACGACGATTGGATACGGAAACGTGGCACCTCGAACATTTGGTGGCCGTC
TATTTGTCATTGGTTATGGTCTAATTGGTATTCCGTTTACA<u>AAC</u>CTGGCAATTGCAGATCTCGGAAAATTCATAT
CAGAAATGATGGTGGAGGCGAAAAGTTTTTGTAGGAAAACCTGGAAAAAACTCAAAAAAGCGTGGAACCCGAATT
TCATTCGCGCAAAGGATCTTTCAAATACGGATATTGAGGAGAAAATATTGGATAATGAGAAAATCGAAAATGAGC
CGGAAACTTCAGAAGTATCAGAAGAAGAAGACGATTTGACAGAGACAGAAGCCACGTCACTTTTCATTTTATTTT
TGGTTTATATCGCATTTGGAGGGTTCATGTTAGCTGCTTATGAACCTGATATGGACTTTTTCAAAGCGGTCTACT
TTAATTTTGTGACATTGACATCAATTGGTCTGGGAGATATTGTACCGAGAAGTGAAACCTATATGCTCATTACAA
TAGTCTACATTGCAATTGGTCTTGCTCTTACCACAATTGCCATTGAAATCGCCGCAGATGCATTGAAGAAGCTGC
ATTATTTTGGAAGAAAAATCGAAAATGTTGGAAATGTGGCTATATGGTTTGGAGGAAAGAAAATTACAATGAAAG
CACTGGTCAAAAACCTCGGTGACCAGTTCAATCTTCCAACTACCGTAGTCAAGAATTTGAATTTGGATCATTTTG
TGGATCAAGCGATTAAAGTGGAGGAAGGAGAGATTGAGACACTCAGACCGCCGCCTTATGAGCCACCGTCTGATC
GATTTGAAGCTGAATTCGCTGATGAACCAGAATGTGAATGGATCCGTGATCCGACTCCAACTCCACCCCCATCAC
CTCAACCGGTTTATCGTCTTCCATCTCCAAAACCAGTTACACCGGAACCTCTACCAAGTCCAACAATAACTGATG
TATCACTTGCGATTGCAACACCATCACCTGAAGAATCGGATGATGATCAAGAACTAATTCTCCCATCACCTGAAC
CGAGTCCAGTTCGAGAGCCAACTCCACCACCGCCTCCACGGGAGCCAACACCTCGTGAGCCAACTCCTGAGCCAG
AGCCAGTTCGAGAGCCGACGCCTCCACCTCCGCCACCTGCCAAGCCCCGTCCACTGACTGCCGCTGAAATCGCGG
CTCAAAAACGCAAAGCGTACAGCGAAGAAGCATGGCGTCGATACCAAGAATACCAGAAACAATGGAAGAAGTTCC
GTCAAACTCAGAAAACTCCCGCACCATCTGGAGCCTCTACATCAGGGGCATCAACATCAAAACCGTCTGGAACAT
CACCGGAAAGTGGAGCCGGAGTATGTACTGGACCATCAACAAGGAGTCAATCAATAACATCAGTTGCATCTGGAA
AGACATCAAGAAGTGCAACACCGGAAAGCAAGAAATCATCACATTTGAGTGGCTCATCGAGAAGAGAAAGCGGTG
GAAAAGGATCCGGATCCGGATCCGGATCCGGAATGTCCGAGCTCATCAAGGAGAACATGCACATGAAGCTCTACA
TGGAGGGAACCGTCAACAACCACCACTTCAAGTGCACCTCCGAGGGAGAGGGAAAGCCATACGAGGGAACCCAAA
CCATGCGTATCAAGgtaagtttaaacatatatatactaactaaccctgattatttaaattttcagGCCGTCGAGG
GAGGACCACTCCCATTCGCCTTCGACATCCTCGCCACCTCCTTCATGTACGGATCCAAGACCTTCATCAACCACA
CCCAAGGAATCCCAGACTTCTTCAAGCAATCCTTCCCAGAGGGATTCACCTGGGAGCGTGTCACCACCTACGAGG
ACGGAGGAGTCCTCACCGCCACCCAAGACACCTCCCTCCAAGACGGATGCCTCATCTACAACGTCAAGATCCGTG
GAGTCAACTTCCCATCCAACGGACCAGTCATGCAAAAGAAGACCCTCGGATGGGAGGCCTCCACCGAGACCCTCT
ACCCAGCCGACGGAGGACTCGAGGGACGTGCCGACATGGCCCTCAAGCTCGTCGGAGGAGGACACCTCATCTGCA
ACCTCAAGgtaagtttaaacatgattttactaactaactaatctgatttaaattttcagACCACCTACCGTTCCA
AGAAGCCAGCCAAGAACCTCAAGATGCCAGGAGTCTACTACGTCGACCGTCGTCTCGAGCGTATCAAGGAGGCCG
ACAAGGAGACCTACGTCGAGCAACACGAGGTCGCCGTCGCCCGTTACTGCGACCTCCCATCCAAGCTCGGACACC
GTTAAaaatcatatgtttttctctctcacactctctttttc

PHX2193: *flp-11(syb2193[flp-11b-SL2(gpd-2)-mKate2-linker-twk-18(e1913)])X*

>flp-11b- SL2(gpd-2)-mKate2(two introns)-linker(GSGSGSGSG)- twk-
18(e1913)(one intron)
gtaagttgtcttaaaatttttcttccgctttttgcctttgcttcatgtgtcgtttattttgctttgcagttcgct
ttggccgatccggtcaactcgaccacatgcacgaccttttgtcgactcttcagAAGCTCAAGTTCGCCAACAACA
AGTAATGACCGAGGACGACCGTCTTCTGCTCGAACAACTCCTG<u>CGA</u>CGAATTCATCATTAAgctgtctcatccta
ctttcacctagttaactgcttgtcttaaaatctatgcttctctttagtatctaaaattttcctagaagcttacaa
gtatataaatggtctcttctcaataaaggttgtatatttattcatcttattgaatctgccatttcctcgttttttg
cgagtttatataccttccaatttctttctattgtattttcaacttctaattttaattcagggaaactgcttcaa
cgcatcaaaaATGTCCGAGCTCATCAAGGAGAACATGCACATGAAGCTCTACATGGAGGGAACCGTCAACAACCA
CCACTTCAAGTGCACCTCCGAGGGAGAGGGAAAGCCATACGAGGGAACCCAAACCATGCGTATCAAGgtaagttt
aaacatatatatactaactaaccctgattatttaaattttcagGCCGTCGAGGGAGGACCACTCCCATTCGCCTT
CGACATCCTCGCCACCTCCTTCATGTACGGATCCAAGACCTTCATCAACCACACCCAAGGAATCCCAGACTTCTT
CAAGCAATCCTTCCCAGAGGGATTCACCTGGGAGCGTGTCACCACCTACGAGGACGGAGGAGTCCTCACCGCCAC
CCAAGACACCTCCCTCCAAGACGGATGCCTCATCTACAACGTCAAGATCCGTGGAGTCAACTTCCCATCCAACGG
ACCAGTCATGCAAAAGAAGACCCTCGGATGGGAGGCCTCCACCGAGACCCTCTACCCAGCCGACGGAGGACTCGA
GGGACGTGCCGACATGGCCCTCAAGCTCGTCGGAGGAGGACACCTCATCTGCAACCTCAAGgtaagtttaaacat
gattttactaactaactaatctgatttaaattttcagACCACCTACCGTTCCAAGAAGCCAGCCAAGAACCTCAA
GATGCCAGGAGTCTACTACGTCGACCGTCGTCTCGAGCGTATCAAGGAGGCCGACAAGGAGACCTACGTCGAGCA
ACACGAGGTCGCCGTCGCCCGTTACTGCGACCTCCCATCCAAGCTCGGACACCGTGGATCCGGATCCGGATCCGG
ATCCGGAATGGCCATCGTCGCCCAAGGAGTCTCCACCATCCTCACCACCTTCCAAAAGACCTTCAAGGGACTCCT
CCCACTCATCATCCTCGTCGCCTACACCCTCCTCGGAGCCTGGATCTTCTGGATGATCGAGGGAGAGAACGAGCG
TGAGATGCTCATCGAGCAACAAAAGGAGCGTGACGAGCTCATCCGTCGTACCGTCTACAAGATCAACCAACTCCA
AATCAAGCGTCAACGTCGTCTCATGACCGCCGAGGAGGAGTACAACCGTACCGCCAAGGTCCTCACCACCTTCCA
AGAGACCCTCGGAATCGTCCCAGCCGACATGGACAAGGACATCCACTGGACCTTCCTCGGATCCATCTTCTACTG
CATGACCGTCTACACCACCATCGGATACGGAAACATCGTCCCAGGAACCGGATGGGGACGTTTCGCCACCATCCT

CTACGCCTTCATCGGAATCCCACTCACCGTCCTCTCCCTCTACTGCCTCGACTCCCTCTTCGCCAAGGGATGCAA
GATGCTCTGGCGTTTCTTCCTCAAGTCCACCCGTGTCGTCTCCAAGGACCTCTCCAACAAGATCTCCGAGGCCGC
CGACAACATCGAGGAGGGAACCACCGCCATCACCCCATCCGCCGAGAAGgtaagtttaaacagttcggtactaac
taaccatacatatttaaattttcagACCGAGAACAACGACGACGACCTCCTCTCCTTCCCAATCTCCGGACTCCT
CCTCATCACCGTCATCTGGGTCATCTTCTGCGCCGTCCTCTTCACCTTCCTCGAGGAGTGGGACTTCGGAACCTC
CCTCTACTTCACCCTCATCTCCTTCACCACCATCGGATTCGGAGACATCCTCCCATCCGACTACGACTTCATGCC
AATCGTCGGAGTCCTCCTCCTCATCGGACTCTCCCTCGTCTCCACCGTCATGACCCTCATCCAACAACAAATCGA
GGCCCTCGCCTCCGGAATGAAGGACAACATCGACCAAGAGTACGCCCGTGCCCTCAACGAGGCCCGTGAGGACGG
AGAGGTCGACGAGCACGTCGACCCAGAGGAGGACCCAGAGAACAACAAGAAGTCCTTCGACGCCGTCATCTCCCG
TATGAACTGGTCCAAGCGTGGACTCTACTACCTCCTCCCAGACTCCCAAAAGAAGGAGCTCGCCAAGCAATCCGA
GAAGAAGATGGACGTAAGTCCATCAAGATCCAAACCGACAACGACCTCCTCGAGACCCTCATCCGTGAGGAGAT
CCTCAAGGCCGAGCTCAACAACGAGATGCACAAGTACACCGCCCCACGTTCCTCCCACCAACCAAAGCTCGTCTA
CTCCGACGTCCGTGAGAAGGAGGTCCCAATCGAGGTCGTCCGTGTCGAGCACTTCAACCACGGAAACGAGGACTA
CCTCGAGCACGACATCTAAaaatcatatgtttttctctctcacactctcttttttc

PHX2493: *lgc-38(syb2346[pflp-11::dpy-10 site::flp-11 3'UTR], syb2493[ReaChR-linker-mKate2])*III

>ReaChR-linker-mKate2 in HB33(flp-11-5'utr::dpy-10 Crispr site::flp-11b-3'utr Chr III)_pre lgc-38
tagcttttccttcctttccgaaatttaatgctattttcaagatgactttttttgcttgcgtttttctcagtttcct
cacacacacacacacacaagtaggcgtggcctgtggaacgtttcagagcgcagaacacctgcatttgatctattc
acttcttgcttttgaaaagcccaaagacaccctacacttcggtttcgttttggaaaccattgacatcatcctatt
ttccataagaagtttccttgagaagaatccatttcgcaaatttttcattaaaacgttcaaaactcatcaaaccat
ttgtaaatagtaataaagtatgtcctgcggctatttgctttctcttcggaatctacaacgccccctcctaataca
tcgtttcaggtataaaaagactgcgcctagccgctcgtctcacttttttgcagttcatactgaata*aaaaa*ATGGT
CTCCCGTCGTCCATGGCTCCTCGCCCTCGCCCTCGCCGTCGCCCTCGCCGCCGGATCCGCCGGAGCCTCCACCGG
ATCCGACGCCACCGTCCCAGTCGCCACCCAAGACGGACCAGACTACGTCTTCCACCGTGCCCACGAGCGTATGCT
CTTCCAAACCTCCTACACCCTCGAGAACAACGGATCCGTCATCTGCATCCCAAACAACGGACAATGCTTCTGCCT
CGCCTGGCTCAAGTCCAACGGAACCAACGCCGAGAAGCTCGCCGCCAACATCCTCCAATGGGTCGTCTTCGCCCT
CTCCGTCGCCTGCCTCGGATGGTACGCCTACCAAGCCTGGCGTGCCACCTGCGGATGGGAGGAGGTCTACGTCGC
CCTCATCGAGATGATGAAGTCCATCATCGAGGCCTTCCACGAGTTCGACTCCCCAGCCACCCTCTGGCTCTCCTC
CGGAAACGGAGTCGTCTGGATGCGTTACGGAGAGTGGCTCCTCACCTGCCCAGTCATCCTCATCCACCTCTCCAA
CCTCACCGGACTCAAGgtaagtttaaacatatatatactaactaaccctgattatttaaattttcagGACGACTA
CTCCAAGCGTACCATGGGACTCCTCGTCTCCGACGTCGGATGCATCGTCTGGGGAGCCACCTCCGCCATGTGCAC
CGGATGGACCAAGATCCTCTTCTTCCTCATCTCCCTCTCCTACGGAATGTACACCTACTTCCACGCCGCCAAGGT
CTACATCGAGGCCTTCCACACCGTCCCAAAGGGACTCTGCCGTCAACTCGTCCGTGCCATGGCCTGGCTCTTCTT
CGTCTCCTGGGGAATGTTCCCAGTCCTCTTCCTCCTCGGACCAGAGGGATTCGGACACATCTCCCCATACGGATC
CGCCATCGGACACTCCATCCTCGACCTCATCGCCAAGgtaagtttaaacagttcggtactaactaaccatacata
tttaaattttcagAACATGTGGGGAGTCCTCGGAAACTACCTCCGTGTCAAGATCCACGAGCACATCCTCCTCTA
CGGAGACATCCGTAAGAAGCAAAAGATCACCATCGCCGGACAAGAGATGGAGGTCGAGACCCTCGTCGCCGAGGA
GGAGGACAAGTACGAGTCCTCCGGAGGATCCGGAGGAGGATCCGGAGGAATGTCCGAGCTCATCAAGGAGAACAT
GCACATGAAGCTCTACATGGAGGGAACCGTCAACAACCACCACTTCAAGTGCACCTCCGAGGGAGAGGGAAAGCC
ATACGAGGGAACCCAAACCATGCGTATCAAGGCCGTCGAGGGAGGACCACTCCCATTCGCCTTCGACATCCTCGC
CACCTCCTTCATGTACGGATCCAAGgtaagtttaaacatgattttactaactaactaatctgatttaaattttca
gACCTTCATCAACCACACCCAAGGAATCCCAGACTTCTTCAAGCAATCCTTCCCAGAGGGATTCACCTGGGAGCG
TGTCACCACCTACGAGGACGGAGGAGTCCTCACCGCCACCCAAGACACCTCCCTCCAAGACGGATGCCTCATCTA
CAACGTCAAGATCCGTGGAGTCAACTTCCCATCCAACGGACCAGTCATGCAAAAGAAGACCCTCGGATGGGAGGC
CTCCACCGAGACCCTCTACCCAGCCGACGGAGGACTCGAGGGACGTGCCGACATGGCCCTCAAGCTCGTCGGAGG
AGGACACCTCATCTGCAACCTCAAGACCACCTACCGTTCCAAGAAGCCAGCCAAGAACCTCAAGATGCCAGGAGT
CTACTACGTCGACCGTCGTCTCGAGCGTATCAAGGAGGCCGACAAGGAGACCTACGTCGAGCAACACGAGGTCGC
CGTCGCCCGTTACTGCGACCTCCCATCCAAGCTCGGACACCGTTAAaaatcatatgtttttctctctcacactct
ctttttcatactctctcttgctgtctagaatttgattggtgtcgcttaaccccctttccctccgaaggaaagt
tatctccccagatctcttttggtgtttttatcagctaacaacacacattttctgatatttctatgctctgtcta
tgaacaataaaggcgttgttaattactcgcaaaatcactttgtttatttttttcacattttcagatagtgaacaa
aagaaaattaaattctaaaatctgaatcggaaaattcaaattaaaaattaaatttattttttttatattacacct
gtttttttcaaatattagatcaaaaactattcaacaagtggcatgtaaagcata**acg**aggtatatgggcttcag
atcttcaactg

PHX3190: *lgc-38(syb2346[pflp-11::dpy-10 site::flp-11 3'UTR], syb2493[[unc-58(e665)-linker-mKate2])*III

>unc-58(e665)-linker(GSGSGSGSG)-mKate2 in HB33(flp-11-5’utr::dpy-10 Crispr site::flp-11b-3’utr Chr III)_pre lgc-38
cacttttttgcagttcatactgaata*aaaaa*ATGGCTCCACTGACTGTGAAAAGCTCACCTCCAAAAAAGGCAAAA
GGAATATCAAAATTTCGGAGAAAAAAGAAGCAGCCACCACCAGACTCAACCGTATTCGTCGCATGGGCACTCCGA
AGTGTCCGAGAGTCTCTGATCCAAGTTGATCCATTGGCCGCTGCACTTGCACATCAGGCTCGAAAGACAAATAGT
GTGCCAGCTGTCTCGAGAACTCCACTGCTTCTACAGTTCACTCCTTTCGGACCACCTCTCAGTGCGTATCATGTG
ACAGCTCGGTGGGAAGGTGCAAATATCAATTCACAATCAGCATTGCTCGATGCAGATGATGGAGCTACAGTTATC
ACAGATACCATCAAAGATGACCAAGATGATAAAGAACCAAAAAGCTGCCCGCAACAGACTGTCAAATACATCAAA
ATACTTACACCTCACGTGATCTTGGTGTCAGTGTTAATTGGATATTTATGCTTGGGAGCTTGGATACTCATGTTA
CTGGAAACAAGGACGGAACTTCTTGCCAGATCCAAAAAACTTGTCAGGTTAACAAATTTGATGTCAAACTTCACT
GCCGAAAGTTGGAAGATGCTCAATAATGCTCAACACGGGGTTAGTAATATGGATGAAGGTGAATGGGCTGCAACA
TTTCGAGAATGGATGGTACGAGTATCAGAAACAGTGGACGATAGGAGACCTATACGACGTGAATTAAACCGGCCT
GATGACTTATCAAATATGCATAATAAATGGACATTTCCAACTGCAATATTATATGTTCTCACTGTGTTAACTACT
TGCGGTTATGGAGAAGTATCTGTCGACACAGACGTCGGAAAGGTTTTCTCAGTAGCATTCGCGCTTGTTGGTATA
CCACTTATGTTCATAACAGCTGCCGATATTGGTAAATTTTTATCTGAAACATTACTCCAGTTTGTGAGCTTTTGG
AATCGAAGTGTCCGAAAAGTGAAGCAATGGATGAGTCGTATTCGTCACGGCAGGAGAAAGTCATTACAATCAACG
GGTGGTCCCAACGATACTCTCGATATTCTTGGTGTCGACGGAACTGAAGAGAAACTTTGGTTCCCAATAGGTGCA
TATGTATCATGTATTTGCATATATTGCTCAATTGGGTCTGCCATGTTTATCACATGGGAAAGAACTTGGTCTTTC
ATTCATGCGTTTCATTTTGGTTTCAATTTGATTGTAACAGTCGGACTCGGAGATATCGTTGTGACTGATTACATA
TTTTTATCACTTATCGTTGCATTTGTGATAGTTGGTTTTTCCGTAGTGACCATGTGCGTGGATCTTGCGTCCACA
CATCTCAAGGCGTACTTCACCAGAATTCACTACTTTGGTCGAGCAAAACGATTCTTAGGAATGAGTGAGGAACTC
AAAGAAATCGTTGCTTTACTGGGGGCGATGCGACGGAAAAAAGGCGGTAAAGTTACATGGAATGATGTGCGAGAC
TTCCTGGATAACGAACTCCGCGATCGACCTTTTGAACCTCATGAGCTTCTGATGAAGCTCAGATTTATTGACGAA
ACATCTTCTGGAATGTCTACAATCCGTCACAATTCCTTCCAGTCAGATTTTTTCCGAGAATCAGAGTACATCCGA
AGAGTGGCTGCGCTGAGGCCAGAACAGCCAGCATATTTGGGATCCGGATCCGGATCCGGATCCGGAATGTCCGAG
CTCATCAAGGAGAACATGCACATGAAGCTCTACATGGAGGGAACCGTCAACAACCACCACTTCAAGTGCACCTCC
GAGGGAGAGGGAAAGCCATACGAGGGAACCCAAACCATGCGTATCAAGgtaagtttaaacatatatatactaact
aaccctgattatttaaattttcagGCCGTCGAGGGAGGACCACTCCCATTCGCCTTCGACATCCTCGCCACCTCC
TTCATGTACGGATCCAAGACCTTCATCAACCACACCCAAGGAATCCCAGACTTCTTCAAGCAATCCTTCCCAGAG
GGATTCACCTGGGAGCGTGTCACCACCTACGAGGACGGAGGAGTCCTCACCGCCACCCAAGACACCTCCCTCCAA
GACGGATGCCTCATCTACAACGTCAAGATCCGTGGAGTCAACTTCCCATCCAACGGACCAGTCATGCAAAAGAAG
ACCCTCGGATGGGAGGCCTCCACCGAGACCCTCTACCCAGCCGACGGAGGACTCGAGGGACGTGCCGACATGGCC
CTCAAGCTCGTCGGAGGAGGACACCTCATCTGCAACCTCAAGgtaagtttaaacatgattttactaactaactaa
tctgatttaaattttcagACCACCTACCGTTCCAAGAAGCCAGCCAAGAACCTCAAGATGCCAGGAGTCTACTAC
GTCGACCGTCGTCTCGAGCGTATCAAGGAGGCCGACAAGGAGACCTACGTCGAGCAACACGAGGTCGCCGTCGCC
CGTTACTGCGACCTCCCATCCAAGCTCGGACACCGTTAAaaatcatatgtttttctctctcacactctc

PHX4110: *lgc-38(syb2346[flp-11p::dpy-10 site::flp-11 3’UTR], syb4110[unc-58gf-CAI-1.0-linker(GSGSGSGSG)-mKate2]) III*

>unc-58(e665CAI)-linker(GSGSGSGSG)-mKate2 in HB33(flp-11-5’utr::dpy-10 Crispr site::flp-11b-3’utr Chr III)_pre lgc-38
cacttttttgcagttcatactgaata*aaaaa*ATGGCCCCACTCACCGTCAAGTCCTCCCCACCAAAGAAGGCCAAG
GGAATCTCCAAGTTCCGTCGTAAGAAGAAGCAACCACCACCAGACTCCACCGTCTTCGTCGCCTGGGCCCTCCGT
TCCGTCCGTGAGTCCCTCATCCAAGTCGACCCACTCGCCGCCGCCCTCGCCCACCAAGCCCGTAAGACCAACTCC
GTCCCAGCCGTCTCCCGTACCCCACTCCTCCTCCAATTCACCCCATTCGGACCACCACTCTCCGCCTACCACGTC
ACCGCCCGTTGGGAGGGAGCCAACATCAACTCCCAATCCGCCCTCCTCGACGCCGACGACGGAGCCACCGTCATC
ACCGACACCATCAAGGACGACCAAGACGACAAGGAGCCAAAGTCCTGCCCACAACAAACCGTCAAGTACATCAAG
ATCCTCACCCCACACGTCATCCTCGTCTCCGTCCTCATCGGATACCTCTGCCTCGGAGCCTGGATCCTCATGCTC
CTCGAGACCCGTACCGAGCTCCTCGCCCGTTCCAAGAAGCTCGTCCGTCTCACCAACCTCATGTCCAACTTCACC
GCCGAGTCCTGGAAGATGCTCAACAACGCCCAACACGGAGTCTCCAACATGGACGAGGGAGAGTGGGCCGCCACC
TTCCGTGAGTGGATGGTCCGTGTCTCCGAGACCGTCGACGACCGTCGTCCAATCCGTCGTGAGCTCAACCGTCCA
GACGACCTCTCCAACATGCACAACAAGTGGACCTTCCCAACCGCCATCCTCTACGTCCTCACCGTCCTCACCACC
TGCGGATACGGAGAGGTCTCCGTCGACACCGACGTCGGAAAGGTCTTCTCCGTCGCCTTCGCCCTCGTCGGAATC
CCACTCATGTTCATCACCGCCGCCGACATCGGAAAGTTCCTCTCCGAGACCCTCCTCCAATTCGTCTCCTTCTGG
AACCGTTCCGTCCGTAAGGTCAAGCAATGGATGTCCCGTATCCGTCACGGACGTCGTAAGTCCCTCCAATCCACC
GGAGGACCAAACGACACCCTCGACATCCTCGGAGTCGACGGAACCGAGGAGAAGCTCTGGTTCCCAATCGGAGCC
TACGTCTCCTGCATCTGCATCTACTGCTCCATCGGATCCGCCATGTTCATCACCTGGGAGCGTACCTGGTCCTTC
ATCCACGCCTTCCACTTCGGATTCAACCTCATCGTCACCGTCGGACTCGGAGACATCGTCGTCACCGACTACATC
TTCCTCTCCCTCATCGTCGCCTTCGTCATCGTCGGATTCTCCGTCGTCACCATGTGCGTCGACCTCGCCTCCACC
CACCTCAAGGCCTACTTCACCCGTATCCACTACTTCGGACGTGCCAAGCGTTTCCTCGGAATGTCCGAGGAGCTC
AAGGAGATCGTCGCCCTCCTCGGAGCCATGCGTCGTAAGAAGGGAGGAAAGGTCACCTGGAACGACGTCCGTGAC

TTCCTCGACAACGAGCTCCGTGACCGTCCATTCGAGCCACACGAGCTCCTCATGAAGCTCCGTTTCATCGACGAG
ACCTCCTCCGGAATGTCCACCATCCGTCACAACTCCTTCCAATCCGACTTCTTCCGTGAGTCCGAGTACATCCGT
CGTGTCGCCGCCCTCCGTCCAGAGCAACCAGCCTACCTCGGATCCGGATCCGGATCCGGATCCGGAATGTCCGAG
CTCATCAAGGAGAACATGCACATGAAGCTCTACATGGAGGGAACCGTCAACAACCACCACTTCAAGTGCACCTCC
GAGGGAGAGGGAAAGCCATACGAGGGAACCCAAACCATGCGTATCAAGgtaagtttaaacatatatatactaact
aaccctgattatttaaattttcagGCCGTCGAGGGAGGACCACTCCCATTCGCCTTCGACATCCTCGCCACCTCC
TTCATGTACGGATCCAAGACCTTCATCAACCACACCCAAGGAATCCCAGACTTCTTCAAGCAATCCTTCCCAGAG
GGATTCACCTGGGAGCGTGTCACCACCTACGAGGACGGAGGAGTCCTCACCGCCACCCAAGACACCTCCCTCCAA
GACGGATGCCTCATCTACAACGTCAAGATCCGTGGAGTCAACTTCCCATCCAACGGACCAGTCATGCAAAAGAAG
ACCCTCGGATGGGAGGCCTCCACCGAGACCCTCTACCCAGCCGACGGAGGACTCGAGGGACGTGCCGACATGGCC
CTCAAGCTCGTCGGAGGAGGACACCTCATCTGCAACCTCAAGgtaagtttaaacatgattttactaactaactaa
tctgatttaaattttcagACCACCTACCGTTCCAAGAAGCCAGCCAAGAACCTCAAGATGCCAGGAGTCTACTAC
GTCGACCGTCGTCTCGAGCGTATCAAGGAGGCCGACAAGGAGACCTACGTCGAGCAACACGAGGTCGCCGTCGCC
CGTTACTGCGACCTCCCATCCAAGCTCGGACACCGTTAAaaatcatatgtttttctc

PHX4416 *aptf-1(gk794)II; flp-11(syb1445 syb4416) X*

>flp-11promoter-unc-58(e665)-linker(GSGSGSGSG)-mKate2(two introns)
tctcttcggaatctacaacgccccctcctaatacatcgtttcaggtataaaaagactgcgcctagccgctcgtct
cactttttgcagttcatactgaataATGGCTCCACTGACTGTGAAAAGCTCACCTCCAAAAAAGGCAAAAGGAAT
ATCAAAATTTCGGAGAAAAAAGAAGCAGCCACCACCAGACTCAACCGTATTCGTCGCATGGGCACTCCGAAGTGT
CCGAGAGTCTCTGATCCAAGTTGATCCATTGGCCGCTGCACTTGCACATCAGGCTCGAAAGACAAATAGTGTGCC
AGCTGTCTCGAGAACTCCACTGCTTCTACAGTTCACTCCTTTCGGACCACCTCTCAGTGCGTATCATGTGACAGC
TCGGTGGGAAGGTGCAAATATCAATTCACAATCAGCATTGCTCGATGCAGATGATGGAGCTACAGTTATCACAGA
TACCATCAAAGATGACCAAGATGATAAAGAACCAAAAAGCTGCCCGCAACAGACTGTCAAATACATCAAAATACT
TACACCTCACGTGATCTTGGTGTCAGTGTTAATTGGATATTTATGCTTGGGAGCTTGGATACTCATGTTACTGGA
AACAAGGACGGAACTTCTTGCCAGATCCAAAAAACTTGTCAGGTTAACAAATTTGATGTCAAACTTCACTGCCGA
AAGTTGGAAGATGCTCAATAATGCTCAACACGGGGTTAGTAATATGGATGAAGGTGAATGGGCTGCAACATTTCG
AGAATGGATGGTACGAGTATCAGAAACAGTGGACGATAGGAGACCTATACGACGTGAATTAAACCGGCCTGATGA
CTTATCAAATATGCATAATAAATGGACATTTCCAACTGCAATATTATATGTTCTCACTGTGTTAACTACTTGCGG
TTATGGAGAAGTATCTGTCGACACAGACGTCGGAAAGGTTTTCTCAGTAGCATTCGCGCTTGTTGGTATACCACT
TATGTTCATAACAGCTGCCGATATTGGTAAATTTTTATCTGAAACATTACTCCAGTTTGTGAGCTTTTGGAATCG
AAGTGTCCGAAAAGTGAAGCAATGGATGAGTCGTATTCGTCACGGCAGGAGAAAGTCATTACAATCAACGGGTGG
TCCCAACGATACTCTCGATATTCTTGGTGTCGACGGAACTGAAGAGAAACTTTGGTTCCCAATAGGTGCATATGT
ATCATGTATTTGCATATATTGCTCAATTGGGTCTGCCATGTTTATCACATGGGAAAGAACTTGGTCTTTCATTCA
TGCGTTTCATTTTGGTTTCAATTTGATTGTAACAGTCGGACTCGGAGATATCGTTGTGACTGATTACATATTTTT
ATCACTTATCGTTGCATTTGTGATAGTTGGTTTTTCCGTAGTGACCATGTGCGTGGATCTTGCGTCCACACATCT
CAAGGCGTACTTCACCAGAATTCACTACTTTGGTCGAGCAAAACGATTCTTAGGAATGAGTGAGGAACTCAAAGA
AATCGTTGCTTTACTGGGGGCGATGCGACGGAAAAAAGGCGGTAAAGTTACATGGAATGATGTGCGAGACTTCCT
GGATAACGAACTCCGCGATCGACCTTTTGAACCTCATGAGCTTCTGATGAAGCTCAGATTTATTGACGAAACATC
TTCTGGAATGTCTACAATCCGTCACAATTCCTTCCAGTCAGATTTTTCCGAGAATCAGAGTACATCCGAAGAGT
GGCTGCGCTGAGGCCAGAACAGCCAGCATATTTGGGATCCGGATCCGGATCCGGATCCGGAATGTCCGAGCTCAT
CAAGGAGAACATGCACATGAAGCTCTACATGGAGGGAACCGTCAACAACCACCACTTCAAGTGCACCTCCGAGGG
AGAGGGAAAGCCATACGAGGGAACCCAAACCATGCGTATCAAGgtaagtttaaacatatatatactaactaaccc
tgattatttaaattttcagGCCGTCGAGGGAGGACCACTCCCATTCGCCTTCGACATCCTCGCCACCTCCTTCAT
GTACGGATCCAAGACCTTCATCAACCACACCCAAGGAATCCCAGACTTCTTCAAGCAATCCTTCCCAGAGGGATT
CACCTGGGAGCGTGTCACCACCTACGAGGACGGAGGAGTCCTCACCGCCACCCAAGACACCTCCCTCCAAGACGG
ATGCCTCATCTACAACGTCAAGATCCGTGGAGTCAACTTCCCATCCAACGGACCAGTCATGCAAAAGAAGACCCT
CGGATGGGAGGCCTCCACCGAGACCCTCTACCCAGCCGACGGAGGACTCGAGGGACGTGCCGACATGGCCCTCAA
GCTCGTCGGAGGAGGACACCTCATCTGCAACCTCAAGgtaagtttaaacatgattttactaactaactaatctga
tttaaattttcagACCACCTACCGTTCCAAGAAGCCAGCCAAGAACCTCAAGATGCCAGGAGTCTACTACGTCGA
CCGTCGTCTCGAGCGTATCAAGGAGGCCGACAAGGAGACCTACGTCGAGCAACACGAGGTCGCCGTCGCCCGTTA
CTGCGACCTCCCATCCAAGCTCGGACACCGTTAAaaatcatatgtttttctctctcacactctctttttttcatac
tctctcttgctgtctagaatttgattggtgtcgcttaaccccccctttccctccgaagga

### Supplementary Figures

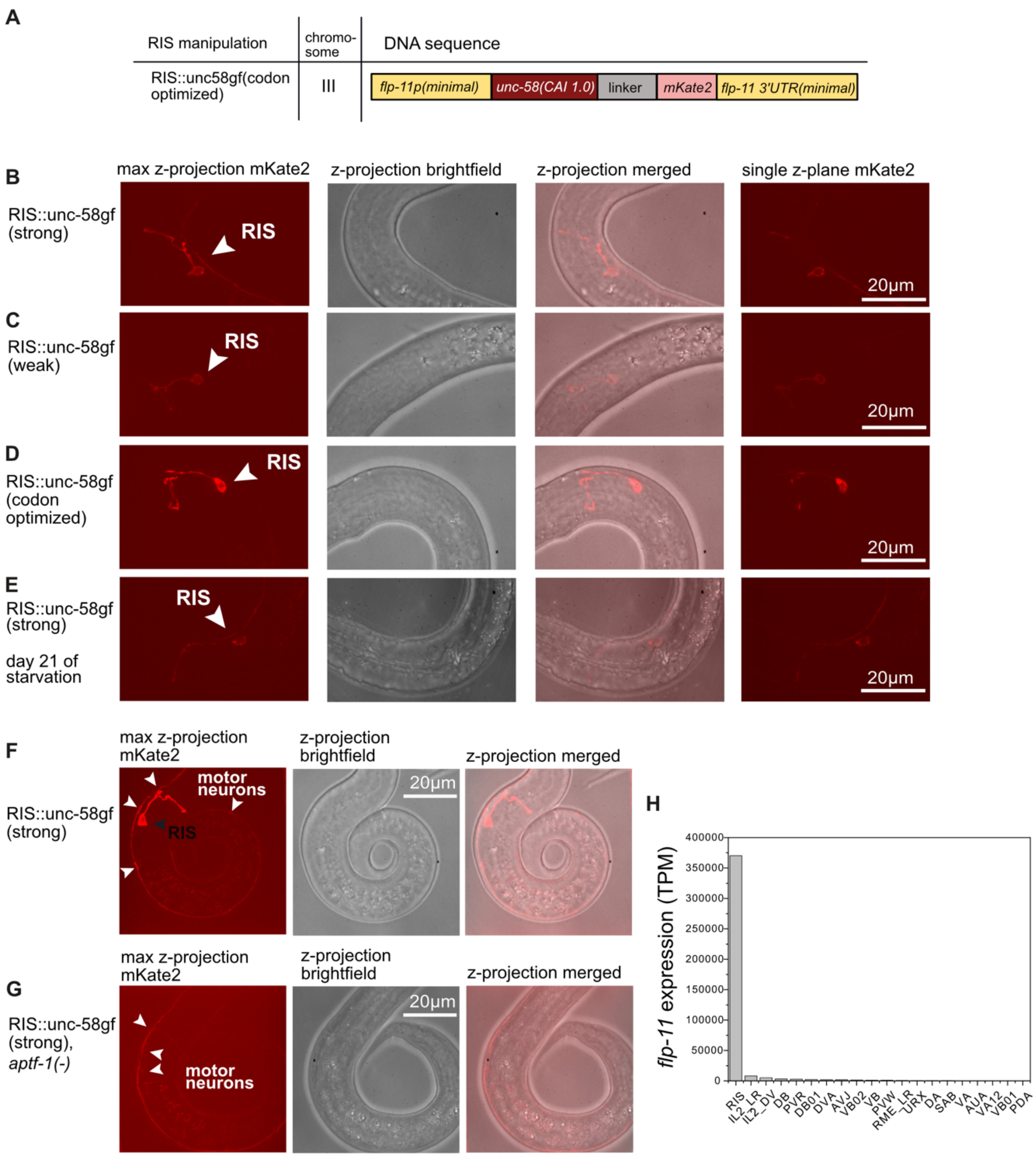


**Fig. S1 – Expression characteristics of *unc-58gf* transgenes and the *flp-11* promoter**

A) Scheme of the genetic design of the *RIS::unc-58gf(weak)* transgene.

B-D) Localization and expression of the different RIS depolarization strains.

E) On day 21 of starvation RIS appears to be intact in the strong depolarization strain (*RIS::unc-58gf(strong)*).

F) Imaging conditions that oversaturate the mKate2 signal in RIS in the *RIS::unc-58gf(strong)* strain reveal a weak expression of the tool in neurons in the ventral cord.

G) The weak expression in the ventral cord neurons but not in RIS is preserved in an *aptf-1* deletion.

H) Transcriptomic data from the Cengen project suggest that the weakly expressing neurons could be IL2 and motor neurons [1].

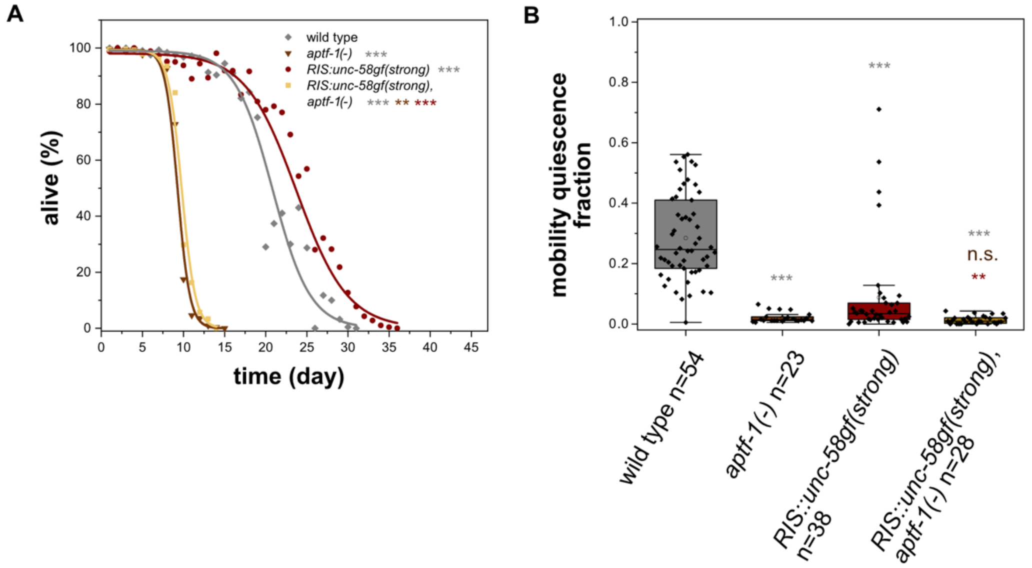


**Fig. S2 *aptf-1(-)* suppresses survival benefit of *RIS::unc-58gf(strong)***

A) The survival benefit of *RIS::unc-58gf(strong)* strongly decreased in the *RIS::unc-58gf(strong), aptf-1(-)*. Fisher's Exact Test was conducted on day 11) when *aptf-1(-)* was the shortest-lived condition and on day 23) when wild type was the shortest-lived condition. The plot includes data from two replicates ***p<0.001.

B) *RIS::unc-58gf(strong)* abolishes quiescence behavior in most individuals similar to but not as strongly as *aptf-1(-)*. A few *RIS::unc-58gf(strong)* animals showed a high fraction of quiescence. To test whether the rare occurrence of increased quiescence phenotype stemmed from *unc-58gf* expression in RIS, we measured quiescence in an *aptf-1(-)* background. This completely abolished quiescence in all animals

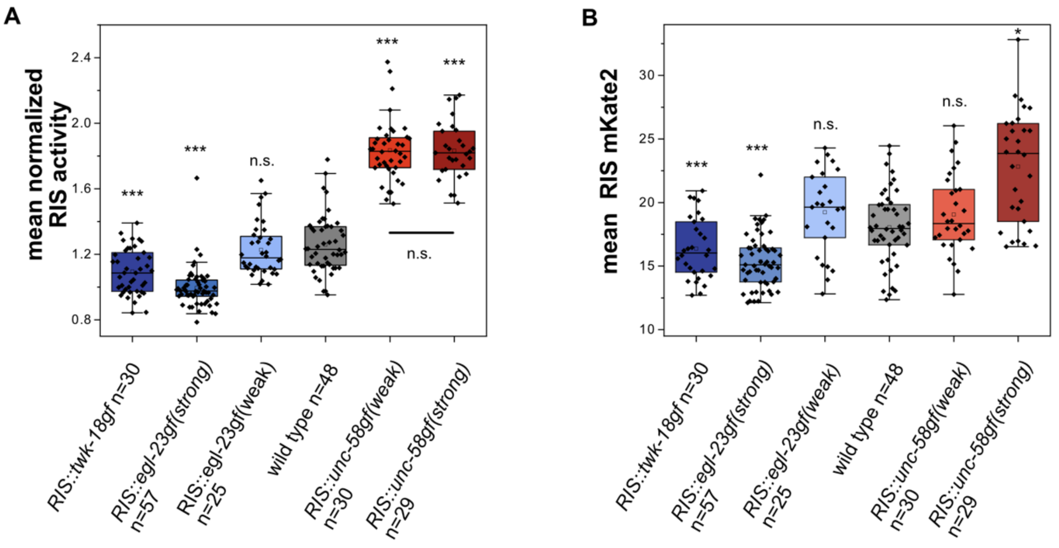


**Fig. S3 – Characterization of RIS activity and behavior across time for the RIS-manipulating transgenes**

A) Mean RIS activity of individual worms of the different RIS activity strains. n.s. p>0.05, **p<0.001, ***p<0.001, Welch test with FDR correction for multiple testing.

B) Mean RIS mKate2 intensities of individual worms as a transcriptional reporter. n.s. p>0.05, *p<0.05, ***p<0.001, Welch test with FDR correction for multiple testing.

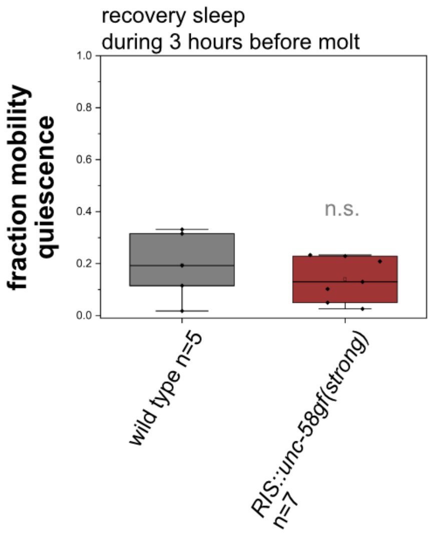


**Fig. S4 - Recovery L1 lethargus after 12 days of larval starvation**

There was no detectable rebound sleep during L1 lethargus after worms were fed after 12 days of starvation. n.s. p>0.05, Welch test.

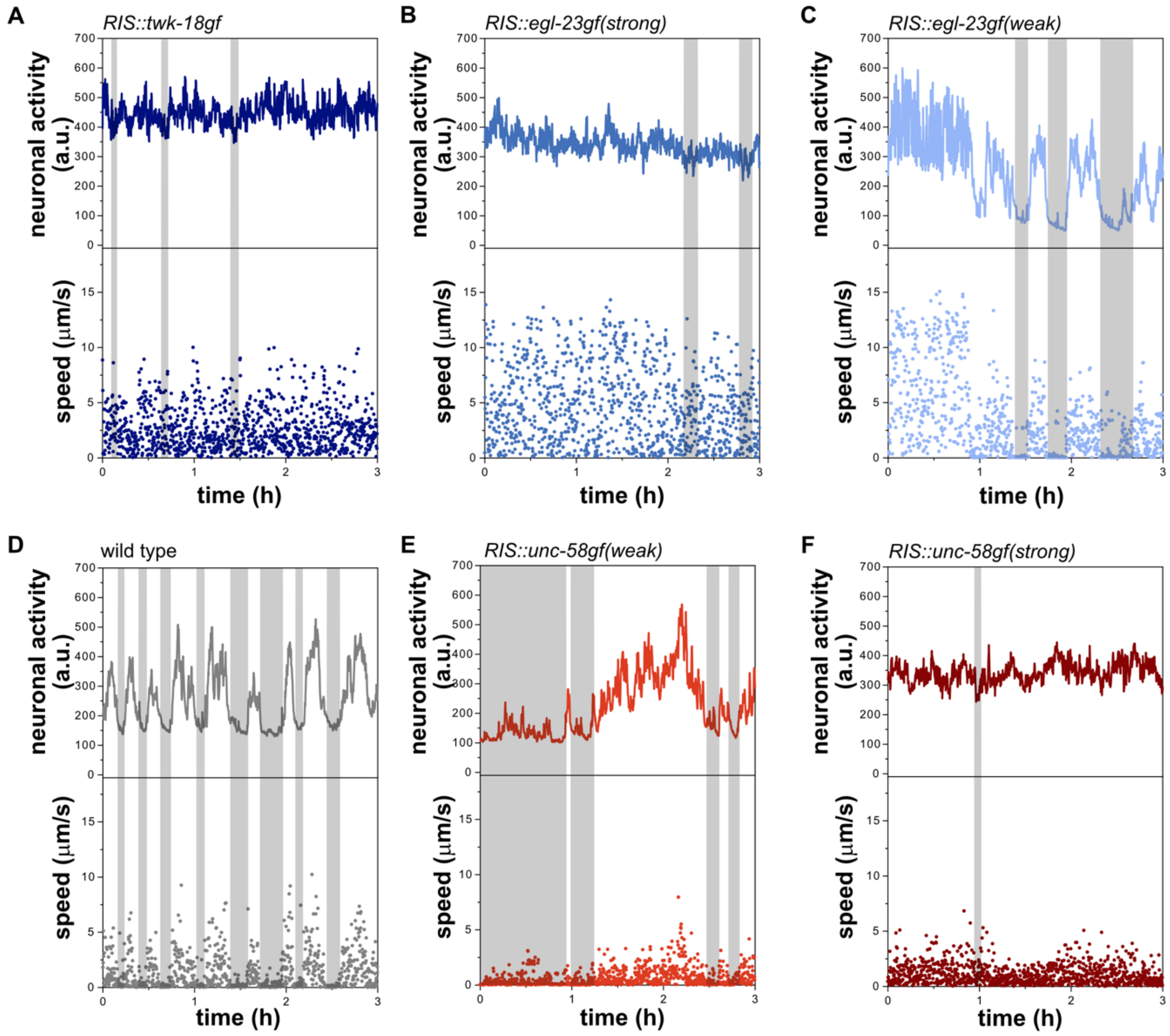


**Fig. S5 – Sample traces of neuronal activity of strains with different RIS activity and optogenetic activation of *RIS::unc-58gf(strong)***

A-F) Sample traces of different RIS activity strains showing neuronal activity and speed after 48h starvation in L1 arrest. Neuronal inactivity bouts, which are depicted in a grey shade, strongly correlate with mobility quiescence.

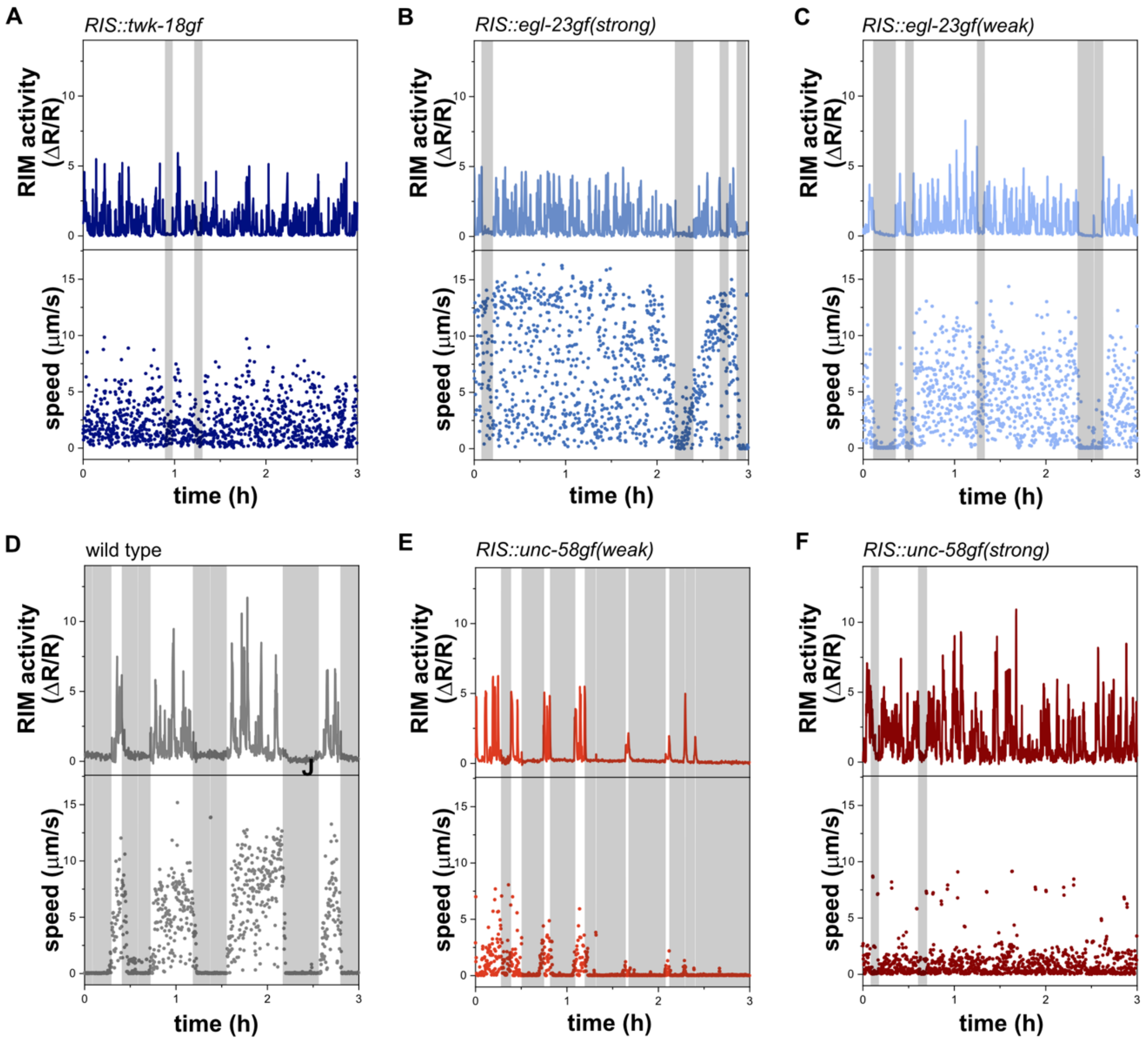


**Fig. S6 – Sample traces of RIM activity of different RIS activity strains**

A-F) Sample traces of strains with different RIS activity are showing RIM activity and speed after 48h starvation in first larval stage arrest. Times of RIM inactivity (grey shaded area) and immobility strongly correlate.

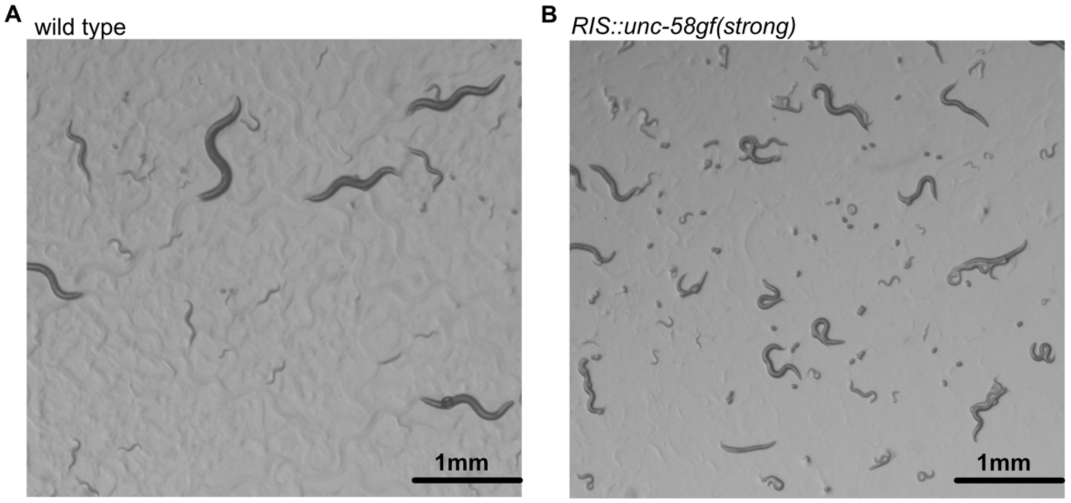


**Fig. S7 The overactivation strain appears to be uncoordinated**
A) Image of wild-type worms on an NGM plate.
B) Image of *RIS::unc-58gf(strong)* worms on an NGM plate.

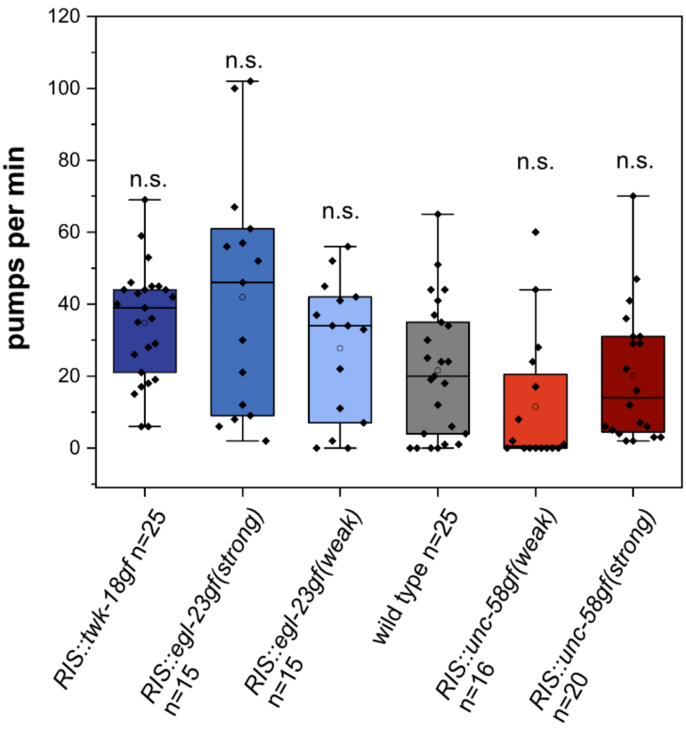


**Fig. S8 Pharyngeal pumping rates**
The pumping rate of *RIS::unc-58gf(strong)* in arrested L1 worms is comparable to the wild type.

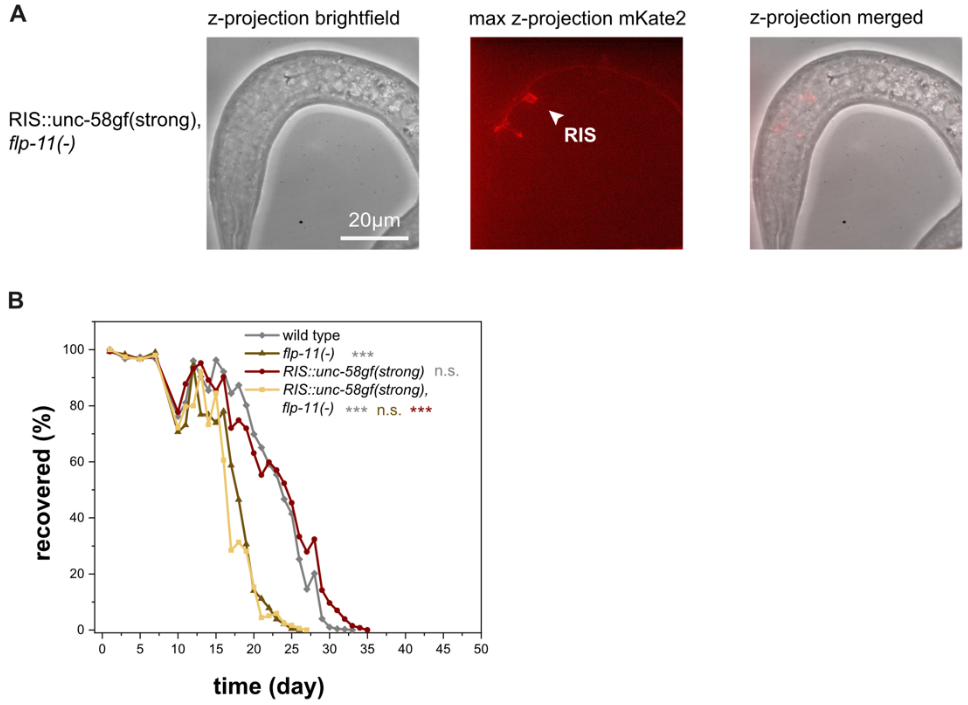


**Fig. S9 – Characterization of the *RIS::unc-58gf(strong), flp-11(-)* strain**

A) The UNC-58gf channel is still expressed at the plasma membrane of RIS in *RIS::unc-58gf(strong), flp-11(-)*.

B) *flp-11(-)* causes strongly reduced recovery rates (Fisher's Exact Test was conducted on day 18) for comparisons when *flp-11(-)* was the shortest lived condition, day 20) when *RIS::unc-58gf(strong), flp-11(-)* was the shortest-lived condition or day 24) when wild type was the shortest-lived condition. The p-values were FDR corrected with Benjamini-Hochberg procedure with a 5% false discovery rate. The plot includes data from three replicates. ***p<0.001.

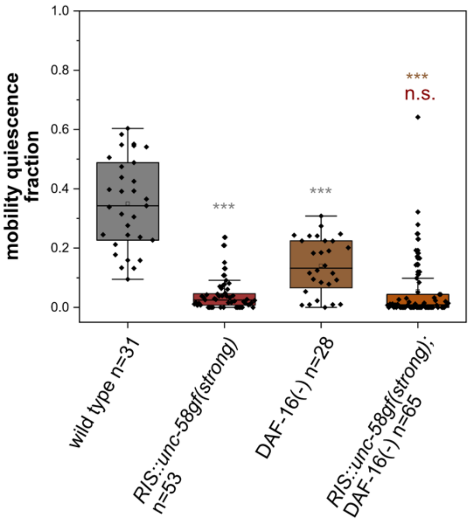


**Fig. S10 Sleep fractions without functional DAF-16.**

Loss of DAF-16 does not change mobility quiescence of *RIS::unc-58gf(strong)*

**Table S1.**

$$y = \frac{a}{1 + e^{-k*(x-x_c)}}$$

| Figure | strain average | a | k | $x_c$ | $R^2$ |
|---|---|---|---|---|---|
| 2B | RIS::twk-18gf | 99.35847 ± 0.66637 | -0.97206 ± 0.12292 | 10.82146 ± 0.33634 | 0.99956 |
| 2B | RIS::egl-23gf(strong) | 100.04807 ± 0.44986 | -0.48094 ± 0.02632 | 17.15292 ± 0.2396 | 0.99956 |
| 2B | RIS::egl-23gf(weak) | 97.32902 ± 0.63724 | -0.41377 ± 0.03302 | 21.60212 ± 0.40667 | 0.99842 |
| 2B | wild type | 98.76819 ± 0.81028 | -0.36143 ± 0.0284 | 21.87268 ± 0.44843 | 0.99803 |
| 2B | RIS::unc-58gf(weak) | 100.02342 ± 1.01326 | -0.24451 ± 0.01495 | 22.19817 ± 0.40979 | 0.99807 |
| 2B | RIS::unc-58gf(strong) | 98.12395 ± 0.74149 | -0.30653 ± 0.02368 | 25.51191 ± 0.46922 | 0.99738 |
| 12B | wild type | 98.96086± 0.83837 | -0.36524± 0.03139 | 22.58115± 0.48943 | 0.99735 |
| 12B | *flp-11(-)* | 98.56064± 0.71073 | -0.48376± 0.04287 | 17.90444± 0.39309 | 0.99859 |
| 12B | RIS:unc-58gf(strong) | 97.36395± 0.51755 | -0.34025± 0.02311 | 26.12587± 0.37504 | 0.99827 |
| 12B | RIS:unc-58gf(strong), *flp-11(-)* | 98.50514± 0.81164 | -0.38458± 0.03107 | 19.2531± 0.40608 | 0.99817 |
| S2A | RIS::unc-58gf(strong), *aptf-1(-)* | 99.28789 ± 0.30363 | -1.2192 ± 0.11022 | 9.78636± 0.20379 | 0.99985 |
| S2A | *aptf-1(-)* | 99.46017± 0.3177 | -0.32131± 0.0232 | 23.73166± 0.44811 | 0.99989 |
| S2A | RIS::unc-58gf(strong) | 98.15331± 0.66903 | -0.32131± 0.0232 | 23.73166± 0.44811 | 0.99774 |
| S2A | wild type | 99.01336± 0.50766 | -0.43604± 0.03443 | 20.83339± 0.37968 | 0.99867 |

3 parameter logistic fit for lifespans.

**Table S2.**

| | |
|---|---|
| CF1038 | *daf-16(mu86) I.* CGC |
| HBR4 | *goeIs3(pmyo-3p::SL1-GCamP3.35-SL2::unc54-3'utr, unc-119(+)).* [2] |
| HBR507 | *flp-11(tm2706)* X. [3] |
| HBR560 | *goeIs120 (tdc-1p::SL1-GCaMP3.35-SL2::mKate2-unc-54-3'utr, unc119(+)).* [4] |
| HBR923 | *goeIs207(mec-4p::SL1-GCaMP6s::mkate2-unc-54-3'utr, unc-119(+)).* |
| HBR1361 | *goeIs304(flp-11p::SL1-GCaMP3.35-SL2::mKate2-unc-54-3'UTR, unc-119(+)).* [5] |
| HBR2340 | *flp-11(syb1445[flp-11-SL2-unc-58(L428F)-linker-mKate2])* X. Generated for this study by outcrossing PHX1445 with N2. |
| HBR2370 | *flp-11(syb2193[flp-11-SL2(gpd-2)-mKate2-linker-twk-18(e1913)])* X*; goeIs304(flp-11p::SL1-GCaMP3.35-SL2::mKate2-unc-54-3'UTR, unc-119(+)).* Generated for this study by crossing HBR1361 with PHX2193. |
| HBR2371 | *flp-11(syb1445[flp-11-SL2-unc-58(L428F)-linker-mKate2])* X*; goeIs304(flp-11p::SL1-GCaMP3.35-SL2::mKate2-unc-54-3'UTR.* Generated for this study by crossing HBR1361 with HBR2340. |
| HBR2421 | *flp-11(syb1445[flp-11-SL2-unc-58(L428F)-linker-mKate2])* X*; otIs672(rab-3p::NLS::GCaMP6s, arrd-4p:NLS:::GCaMP6s).* Generated for this study by crossing OH15265 with HBR2340. |
| HBR2446 | *flp-11(syb1445[flp-11-SL2-unc-58(L428F)-linker-mKate2]) X, goeIs120(tdc-1p::SL1-GCaMP3.35-SL2::mKate2-unc-54-3'utr,unc119(+)).* Generated for this study by crossing PHX1445 and HBR560. |
| HBR2470 | *flp-11(syb2193[flp-11b-SL2(gpd-2)-mKate2-linker-twk-18(e1913)]) X, otIs672(rab-3p::NLS::GCaMP6s, arrd-4p:NLS:::GCaMP6s).* Generated for this study by crossing PHX2193 and OH15265. |
| HBR2508 | *flp-11(syb2193[flp-11b-SL2(gpd-2)-mKate2-linker-twk-18(e1913)]) X; goeIs120(tdc-1p::SL1-GCaMP3.35-SL2::mKate2-unc-54-3'utr,unc119(+)).* Generated for this study by crossing PHX2193 and HBR560. |
| HBR2522 | *lgc-38(syb2346[flp-11p::dpy-10 site::flp-11 3'UTR], syb3190[unc-58(e665)-linker(GSGSGSGSG)-mKate2 ]) III; goeIs304(flp-11p::SL1-GCaMP3.35-SL2::mKate2-unc-54-3'UTR, unc-119(+)).* Generated for this study by crossing PHX3190 and HBR1361. |

| | |
|---|---|
| HBR2523 | *lgc-38(syb2346[pflp-11::dpy-10 site::flp-11 3'UTR], syb2493[ReaChR-linker-mKate2]) III; goeIs304(pflp-11::SL1-GCaMP3.35-SL2::mKate2-unc-54-3'UTR, unc-119(+))*. Generated for this study by crossing HBR1361 and PHX2493. |
| HBR2540 | *flp-11(syb1433[flp-11-SL2-egl-23cDNA(A383V)-linker-mKate2-N2])X; otIs672(rab-3p::NLS::GCaMP6s, arrd-4p:NLS:::GCaMP6s).* Generated for this study by crossing PHX1433 and OH15265. |
| HBR2541 | *flp-11(syb1464)[flp-11-SL2-egl-23cDNA(L229N)-linker-mKate2-N2]) X;otIs672(rab-3p::NLS::GCaMP6s, arrd-4p:NLS:::GCaMP6s).* Generated for this study by crossing PHX1464 and OH15265. |
| HBR2542 | *flp-11(syb1433[flp-11-SL2-egl-23cDNA(A383V)-linker-mKate2-N2]) X; goeIs304(flp-11p::SL1-GCaMP3.35-SL2::mKate2-unc-54-3'UTR, unc-119(+)).* Generated for this study by crossing PHX1433 and HBR1361. |
| HBR2543 | *flp-11(syb1464)[flp-11-SL2-egl-23cDNA(L229N)-linker-mKate2-N2]) X; goeIs304(flp-11p::SL1-GCaMP3.35-SL2::mKate2-unc-54-3'UTR, unc-119(+)).* Generated for this study by crossing PHX1464 and HBR1361. |
| HBR2544 | *lgc-38(syb2346[flp-11p::dpy-10 site::flp-11 3'UTR] III, syb3190[unc-58(e665)-linker(GSGSGSGSG)-mKate2 ]); goeIs120(tdc-1p::SL1-GCaMP3.35-SL2::mKate2-unc-54-3'utr,unc119(+)).* Generated for this study by crossing PHX3190 and HBR560. |
| HBR2545 | *daf-16(mu86) I; flp-11(syb1445[flp-11-SL2-unc-58(L428F)-linker-mKate2])* X. [6] |
| HBR2584 | *lgc-38(syb2346[flp-11p::dpy-10 site::flp-11 3'UTR], syb2493[ReaChR-linker-mKate2])* III *; flp-11(syb1445[flp-11-SL2-unc-58(L428F)-linker-mKate2])* X*; goeIs304(flp-11p::SL1-GCaMP3.35-SL2::mKate2-unc-54-3'UTR.* Generated for this study by crossing HBR2371 and PHX2493. |
| HBR2587 | *flp-11(syb1433[flp-11-SL2-egl-23cDNA(A383V)-linker-mKate2-N2] ) X; goeIs120(tdc-1p::SL1-GCaMP3.35-SL2::mKate2-unc-54-3'utr,unc119(+)).* Generated for this study by crossing HBR560 and PHX1433. |
| HBR2628 | *flp-11(syb1464[flp-11-SL2-egl-23cDNA(L229N)-linker-mKate2-N2]) X; goeIs120(tdc-1p::SL1-GCaMP3.35-SL2::mKate2-unc-54-3'utr,unc119(+)).* Generated for this study by crossing HBR560 and PHX1464. |
| HBR2629 | *lgc-38(syb2346[flp-11p::dpy-10 site::flp-11 3'UTR], syb3190[unc-58(e665)-linker(GSGSGSGSG)-mKate2 ]) III, otIs672(rab-3p::NLS::GCaMP6s, arrd-4p:NLS:::GCaMP6s).* Generated for this study by crossing PHX3190 and OH15265. |
| HBR2652 | *goeIs207(mec-4p::SL1-GCaMP6s::mkate2-unc-54-3'utr, unc-119(+)); flp-11(syb2193[flp-11-SL2-mKate2-linker-twk-18(e1913)])* X. Generated for this study by crossing PHX2193 with HBR923. |

| | |
|---|---|
| HBR2656 | *goeIs207(mec-4p::SL1-GCaMP6s::mkate2-unc-54-3'utr, unc-119(+)); flp-11(syb1445[flp-11-SL2-unc-58(L428F)-linker-mKate2])* X. Generated for this study by crossing HBR2340 and HBR923. |
| HBR2658 | *flp-11(syb1445[flp-11-SL2-unc-58(L428F)-linker-mKate2] syb4416[flp-11-deletion]) X* . Generated for this study by outcrossing PHX4416 with N2. |
| HBR2744 | *goeIs3(pmyo-3p::SL1-GCamP3.35-SL2::unc54-3'utr, unc-119(+)); flp-11(syb2193[flp-11-SL2-mKate2-linker-twk-18(e1913)])* X. Generated for this study by crossing PHX2193 and HBR4. |
| HBR2748 | *goeIs3(pmyo-3p::SL1-GCamP3.35-SL2::unc54-3'utr, unc-119(+)); flp-11(syb1445[flp-11-SL2-unc-58(L428F)-linker-mKate2])* X. Generated for this study by crossing HBR2340 and HBR4. |
| N2 | wild type (Bristol) [7] |
| OH15265 | *otIs672(rab-3p::NLS::GCaMP6s, arrd-4p:NLS::GCaMP6s).* [8] |
| PHX1433 | *flp-11(syb1433[flp-11-SL2-egl-23cDNA(A383V)-linker-mKate2])* X. Generated by Sunybiotech according to our design for this study. |
| PHX1445 | *aptf-1(gk794)II; flp-11(syb1445)[flp-11-SL2-unc-58(L428F)-linker-mKate2]* X. Generated by Sunybiotech according to our design for this study. |
| PHX1464 | *flp-11(syb1464[flp-11-SL2-egl-23cDNA(L229N)-linker-mKate2])* X. Generated by Sunybiotech according to our design for this study. |
| PHX2193 | *flp-11(syb2193[flp-11-SL2-mKate2-linker-twk-18(e1913)])* X. Generated by Sunybiotech according to our design for this study. |
| PHX2493 | *lgc-38(syb2346[flp-11p::dpy-10 site::flp-11 3'UTR], syb2493[ReaChR-linker-mKate2])* III. Generated by Sunybiotech according to our design for this study. |
| PHX3190 | *lgc-38(syb2346[flp-11p::dpy-10 site::flp-11 3'UTR], syb3190[unc-58(e665)-linker(GSGSGSGSG)-mKate2 ])* III. Generated by Sunybiotech according to our design for this study. |
| PHX4110 | *lgc-38(syb2346[flp-11p::dpy-10 site::flp-11 3'UTR], syb4110[unc-58gf-CAI-1.0-linker(GSGSGSGSG)-mKate2]) III.* Generated by Sunybiotech according to our design for this study. |
| PHX4416 | *aptf-1(gk794) II; flp-11(syb1445 syb4416) X.* Generated by Sunybiotech according to our design for this study. |

Strain list for this study.

**Table S3**

*aptf-1(gk794)*
CGACAATCTTCCCAAAGACC
CGGATCGATTGCTAGAGAGG
GCTTGGACGGCTTTAGTTGA

*flp-11(syb1445)* these primers were utilized for all strains in which a tool was knocked into the endogenous locus of *flp-11* except for PHX4416
ACGAGGAAGACTTTGCTCCA
AAACTCGCAAAAACGAGGAA
GACACCAATCAAATTCTAGACAGC

*flp-11p::SL2::unc-58(L428F)*
GACCACATGCACGACCTTTT
ATGACTTTCTCCTGCCGTGA

*flp-11(syb4416)*
ACTAGAACAAGCGTCCTCAA
TCCAATTAACACTGACACCA

*lgc-38(syb2346)* these primers were utilized for all strains in which a tool was integrated into the ski-lodge site on chromosome 3.
ATGGCGATGTCATTTTCATGTT
AGACCACCTACCGTTCCAAG
ATCCCAGTTGTTTGACGGTT

*flp-11(tm2706)*
TCTTCCAAATCGAACCAAGG
TAGCCGCTCGTCTCACTTTT
ATGATGAATTCGCCTCAGGA

List of primers utilized in this study.

**Table S4**

| Figure | group 1 | group 2 | statistical test | p-value |
|---|---|---|---|---|
| 1F | *RIS::unc-58gf(weak)* | *RIS::unc-58gf(strong)* | Welch test | 2.93904E-5 |
| 2B | wild type | *RIS::twk-18* | Fisher's Exact test with FDR correction for multiple testing | 2.6472e-26 |
| | wild type | *RIS::egl23(strong)* | Fisher's Exact test with FDR correction for multiple testing | 2.0137e-33 |
| | wild type | *RIS::egl-23(weak)* | Fisher's Exact test with FDR correction for multiple testing | 0.060809 |
| | wild type | *RIS::unc-58gf(weak)* | Fisher's Exact test with FDR correction for multiple testing | 0.23821 |
| | wild type | *RIS::unc-58gf(strong)* | Fisher's Exact test with FDR correction for multiple testing | 3.856e-15 |
| 2C | wild type | *RIS::twk-18* | Fisher's Exact test with FDR correction for multiple testing | 2.9467e-27 |
| | wild type | *RIS::egl23(strong)* | Fisher's Exact test with FDR correction for multiple testing | 9.507e-39 |
| | wild type | *RIS::egl-23(weak)* | Fisher's Exact test with FDR correction for multiple testing | 1 |

| | | | | |
|---|---|---|---|---|
| | wild type | *RIS::unc-58gf(weak)* | Fisher's Exact test with FDR correction for multiple testing | 0.011154 |
| | wild type | *RIS::unc-58gf(strong)* | Fisher's Exact test with FDR correction for multiple testing | 0.0023327 |
| 3A | wild type | *RIS::twk-18* | Welch test with FDR correction for multiple testing | 0.005595 |
| | wild type | *RIS::egl23(strong)* | Welch test with FDR correction for multiple testing | 5.6391e-05 |
| | wild type | *RIS::egl-23(weak)* | Welch test with FDR correction for multiple testing | 0.55774 |
| | wild type | *RIS::unc-58gf(weak)* | Welch test with FDR correction for multiple testing | 1.2537e-13 |
| | wild type | *RIS::unc-58gf(strong)* | Welch test with FDR correction for multiple testing | 6.23e-14 |
| | *RIS::unc-58gf(weak)* | *RIS::unc-58gf(strong)* | Welch test with FDR correction for multiple testing | 0.011028 |
| 4G | wild type pre bout RIS activity | wild type during bout RIS activity | Wilcoxon signed rank test | 8.6542e-12 |
| | *RIS::egl-23(weak)* pre bout RIS activity | *RIS::egl-23(weak)* during bout RIS activity | Wilcoxon signed rank test | 0.002747 |
| | *RIS::unc58gf(weak)* pre bout RIS activity | *RIS::unc58gf(weak)* during bout RIS activity | Wilcoxon signed rank test | 1.1124e-15 |
| | *RIS::unc58gf(strong)* pre bout RIS activity | *RIS::unc58gf(strong)* during bout RIS activity | Wilcoxon signed rank test | 0.7334 |

| | | | | |
|---|---|---|---|---|
| | *RIS::twk-18gf* pre bout RIS activity | *RIS::twk-18gf* during bout RIS activity | Wilcoxon signed rank test | 0.46484 |
| | wild type pre bout speed | wild type during speed | Wilcoxon signed rank test | 1.4524e-17 |
| | *RIS::egl-23(weak)* pre bout speed | *RIS::egl-23(weak)* during bout speed | Wilcoxon signed rank test | 3.4946e-18 |
| | *RIS::unc58gf(weak)* pre bout speed | *RIS::unc58gf(weak)* during bout speed | Wilcoxon signed rank test | 1.1813e-17 |
| | *RIS::unc58gf(strong)* pre bout speed | *RIS::unc58gf(strong)* during bout speed | Wilcoxon signed rank test | 0.00683 |
| | *RIS::twk-18gf* pre bout speed | *RIS::twk-18gf* during bout speed | Wilcoxon signed rank test | 0.026855 |
| 4H | wild type | *RIS::twk-18* | Welch test with FDR correction for multiple testing | 4.9986e-09 |
| | wild type | *RIS::egl23(strong)* | Welch test with FDR correction for multiple testing | 4.9986e-09 |
| | wild type | *RIS::egl-23(weak)* | Welch test with FDR correction for multiple testing | 0.00539 |
| | wild type | *RIS::unc-58gf(weak)* | Welch test with FDR correction for multiple testing | 0.00090302 |
| | wild type | *RIS::unc-58gf(strong)* | Welch test with FDR correction for multiple testing | 5.8071e-09 |
| | *RIS::unc-58gf(weak)* | *RIS::unc-58gf(strong)* | Welch test with FDR correction for multiple testing | 1.8193e-11 |
| 5G | wild type | *RIS::twk-18* | Welch test with FDR correction for multiple testing | 0.00010291 |

| | | | | |
|---|---|---|---|---|
| | wild type | *RIS::egl23(strong)* | Welch test with FDR correction for multiple testing | 0.0001291 |
| | wild type | *RIS::egl-23(weak)* | Welch test with FDR correction for multiple testing | 0.00754 |
| | wild type | *RIS::unc-58gf(weak)* | Welch test with FDR correction for multiple testing | 0.00010291 |
| | wild type | *RIS::unc-58gf(strong)* | Welch test with FDR correction for multiple testing | 0.0002277 |
| 5H | wild type | *RIS::unc-58gf(strong)* | Welch test | 6.02442e-04 |
| 6B | wild type pre bout neuronal activity | wild type during bout neuronal activity | Wilcoxon signed rank test | 2.3261e-09 |
| | *RIS::egl-23(weak)* pre bout neuronal activity | *RIS::egl-23(weak)* during bout neuronal activity | Wilcoxon signed rank test | 1.7486e-10 |
| | *RIS::unc58gf(weak)* pre bout neuronal activity | *RIS::unc58gf(weak)* during bout neuronal activity | Wilcoxon signed rank test | 0.012606 |
| | *RIS::twk-18gf* pre bout neuronal activity | *RIS::twk-18gf* during bout neuronal activity | Wilcoxon signed rank test | 0.26272 |
| | *RIS::unc58gf(strong)* pre bout neuronal activity | *RIS::unc58gf(strong)* during bout neuronal activity | Wilcoxon signed rank test | 0.049438 |
| | wild type pre bout speed | wild type during speed | Wilcoxon signed rank test | 2.8687e-17 |
| | *RIS::egl-23(weak)* pre bout speed | *RIS::egl-23(weak)* during bout speed | Wilcoxon signed rank test | 8.84e-17 |
| | *RIS::unc58gf(weak)* pre bout speed | *RIS::unc58gf(weak)* during bout speed | Wilcoxon signed rank test | 3.4075e-15 |
| | *RIS::twk-18gf* pre bout speed | *RIS:: twk-18gf* during bout speed | Wilcoxon signed rank test | 0.007 |
| | *RIS::unc58gf(strong)* pre bout speed | *RIS::unc58gf(strong)* during speed | Wilcoxon signed rank test | 0.001709 |
| 6D | wild type | *RIS::twk-18* | Welch test with FDR correction | 6.6283e-05 |

| | | | | |
|---|---|---|---|---|
| | | | for multiple testing | |
| | wild type | *RIS::egl23(strong)* | Welch test with FDR correction for multiple testing | 0.025867 |
| | wild type | *RIS::egl-23(weak)* | Welch test with FDR correction for multiple testing | 0.34555 |
| | wild type | *RIS::unc-58gf(weak)* | Welch test with FDR correction for multiple testing | 0.38777 |
| | wild type | *RIS::unc-58gf(strong)* | Welch test with FDR correction for multiple testing | 0.0042 |
| 6F | wild type pre bout RIM activity | wild type during bout RIM activity | Wilcoxon signed rank test | 1.1401e-05 |
| | *RIS::egl-23(weak)* pre bout RIM activity | *RIS::egl-23(weak)* during bout RIM activity | Wilcoxon signed rank test | 0.03467 |
| | *RIS::unc58gf(weak)* pre bout RIM activity | *RIS::unc58gf(weak)* during bout RIM activity | Wilcoxon signed rank test | 0.001616 |
| | *RIS::twk-18gf* pre bout RIM activity | *RIS::twk-18gf* during bout RIM activity | Wilcoxon signed rank test | 0.30078 |
| | *RIS::unc58gf(strong)* pre bout RIM activity | *RIS::unc58gf(strong)* during bout RIM activity | Wilcoxon signed rank test | 0.1288 |
| | wild type pre bout speed | wild type during speed | Wilcoxon signed rank test | 4.3681e-18 |
| | *RIS::egl-23(weak)* pre bout speed | *RIS::egl-23(weak)* during bout speed | Wilcoxon signed rank test | 2.3544e-13 |
| | *RIS::unc58gf(weak)* pre bout speed | *RIS::unc58gf(weak)* during bout speed | Wilcoxon signed rank test | 1.0993e-12 |
| | *RIS::twk-18gf* pre bout speed | *RIS::twk-18gf* during bout speed | Wilcoxon signed rank test | 2.732e-05 |
| | *RIS::unc58gf(strong)* pre bout speed | *RIS::unc58gf(strong)* during bout speed | Wilcoxon signed rank test | 1.1004e-07 |
| 6H | wild type | *RIS::twk-18* | Welch test with FDR | 0.00055411 |

| | | | | |
|---|---|---|---|---|
| | | | correction for multiple testing | |
| | wild type | *RIS::egl23(strong)* | Welch test with FDR correction for multiple testing | 0.00735 |
| | wild type | *RIS::egl-23(weak)* | Welch test with FDR correction for multiple testing | 0.00055411 |
| | wild type | *RIS::unc-58gf(weak)* | Welch test with FDR correction for multiple testing | 0.00645 |
| | wild type | *RIS::unc-58gf(strong)* | Welch test with FDR correction for multiple testing | 8.7669e-06 |
| 7D | wild type | *RIS::twk-18* | Welch test | 7.04184e-04 |
| | wild type | *RIS::unc-58gf(strong)* | Welch test | 0.0183 |
| 7E | wild type | *RIS::twk-18* | Welch test | 0.00183 |
| | wild type | *RIS::unc-58gf(strong)* | Welch test | 0.00398 |
| 7F | wild type quiescent bouts | *RIS::twk-18* quiescent bouts | Welch test | 0.03172 |
| | wild type quiescent bouts | *RIS::unc-58gf(strong)* quiescent bouts | Welch test | 0.54995 |
| | wild type mobile bouts | *RIS::twk-18* mobile bouts | Welch test | 0.00283 |
| | wild type mobile bouts | *RIS::unc-58gf(strong)* mobile bouts | Welch test | 0.84769 |
| | wild type quiescent bouts | wild type mobile bouts | Wilcoxon signed rank test | 0.00103 |
| | *RIS::twk-18* quiescent bouts | *RIS::twk-18* mobile bouts | Wilcoxon signed rank test | 2.26067e-07 |
| | *RIS::unc-58gf(strong)* quiescent bouts | *RIS::unc-58gf(strong)* mobile bouts | Wilcoxon signed rank test | 0.00103 |
| 8A | wild type quiescent bouts | *RIS::twk-18* quiescent bouts | Welch test | 0.00409 |
| | wild type quiescent bouts | *RIS::unc-58gf(strong)* quiescent bouts | Welch test | 0.04721 |
| | wild type mobile bouts | *RIS::twk-18* mobile bouts | Welch test | 0.06568 |
| | wild type mobile bouts | *RIS::unc-58gf(strong)* mobile bouts | Welch test | 0.00162 |

| | | | | |
|---|---|---|---|---|
| | wild type quiescent bouts | wild type mobile bouts | Wilcoxon signed rank test | 7.45058e-09 |
| | *RIS::twk-18* quiescent bouts | *RIS::twk-18* mobile bouts | Wilcoxon signed rank test | 0.00781 |
| | *RIS::unc-58gf(strong)* quiescent bouts | *RIS::unc-58gf(strong)* mobile bouts | Wilcoxon signed rank test | 1.90735e-06 |
| 8B | wild type quiescent bouts | *RIS::twk-18* quiescent bouts | Welch test | 5.57801e-10 |
| | wild type quiescent bouts | *RIS::unc-58gf(strong)* quiescent bouts | Welch test | 0.22981 |
| | wild type mobile bouts | *RIS::twk-18* mobile bouts | Welch test | 0.74543 |
| | wild type mobile bouts | *RIS::unc-58gf(strong)* mobile bouts | Welch test | 0.22981 |
| | wild type quiescent bouts | wild type mobile bouts | Wilcoxon signed rank test | 9.56917e-05 |
| | *RIS::twk-18* quiescent bouts | *RIS::twk-18* mobile bouts | Wilcoxon signed rank test | 2.88885e-05 |
| | *RIS::unc-58gf(strong)* quiescent bouts | *RIS::unc-58gf(strong)* mobile bouts | Wilcoxon signed rank test | 0.00166 |
| 8F | wild type quiescent bouts | *RIS::twk-18* quiescent bouts | Welch test | 0.71373 |
| | wild type quiescent bouts | *RIS::unc-58gf(strong)* quiescent bouts | Welch test | 0.00957 |
| | wild type mobile bouts | *RIS::twk-18* mobile bouts | Welch test | 0.76635 |
| | wild type mobile bouts | *RIS::unc-58gf(strong)* mobile bouts | Welch test | 0.03191 |
| | wild type quiescent bouts | wild type mobile bouts | Wilcoxon signed rank test | 1.29926e-04 |
| | *RIS::twk-18* quiescent bouts | *RIS::twk-18* mobile bouts | Wilcoxon signed rank test | 0.01563 |
| | *RIS::unc-58gf(strong)* quiescent bouts | *RIS::unc-58gf(strong)* mobile bouts | Wilcoxon signed rank test | 1.49012e-08 |
| 8G | wild type pre bout muscle activity | wild type during bout muscle activity | | 2.2874e-10 |
| | *RIS::twk-18gf* pre bout muscle activity | *RIS::twk-18gf* during bout muscle activity | | 0.82031 |
| | *RIS::unc58gf(strong)* pre bout muscle activity | *RIS::unc58gf(strong)* during bout muscle activity | | 1.6621e-07 |
| | wild type pre bout speed | wild type during speed | | 1.3599e-05 |
| | *RIS::twk-18gf* pre bout speed | *RIS::twk-18gf* during bout speed | | 9.9904e-06 |

| | | | | |
|---|---|---|---|---|
| | *RIS::unc58gf(strong)* pre bout speed | *RIS::unc58gf(strong)* during bout speed | | 6.7468e-05 |
| 8H | wild type | *RIS::twk-18gf* | | 0.01585 |
| | wild type | *RIS::unc-58gf(strong)* | | 0.02974 |
| 9A | wild type pre stimulation RIS activity | wild type during stimulation RIS activity | Wilcoxon signed rank test | 0.00012207 |
| | wild type pre stimulation speed | wild type during stimulation speed | Wilcoxon signed rank test | 0.001709 |
| | *RIS::unc-58gf(strong)* pre stimulation RIS activity | *RIS::unc-58gf(strong)* during stimulation RIS activity | Wilcoxon signed rank test | 0.39443 |
| | *RIS::unc-58gf(strong)* pre stimulation speed | *RIS::unc-58gf(strong)* during stimulation speed | Wilcoxon signed rank test | 0.97574 |
| 9B | wild type pre stimulation RIS activity | wild type during stimulation RIS activity | Wilcoxon signed rank test | 0.54688 |
| | wild type pre stimulation speed | wild type during stimulation speed | Wilcoxon signed rank test | 0.38281 |
| | *RIS::unc-58gf(strong)* pre stimulation RIS activity | *RIS::unc-58gf(strong)* during stimulation RIS activity | Wilcoxon signed rank test | 0.17881 |
| | *RIS::unc-58gf(strong)* pre stimulation speed | *RIS::unc-58gf(strong)* during stimulation speed | Wilcoxon signed rank test | 0.60509 |
| 10C | *RIS::ReaChR(-ATR)* first hour GCaMP | *RIS::ReaChR(+ATR)* first hour GCaMP | Welch test | 0.03655 |
| | *RIS::ReaChR(-ATR)* last hour GCaMP | *RIS::ReaChR(+ATR)* last hour GCaMP | Welch test | 0.049849 |
| | *RIS::ReaChR(-ATR)* first hour speed | *RIS::ReaChR(+ATR)* first hour speed | Welch test | 0.0005971 |
| 10E | *RIS::ReaChR(-ATR)* | *RIS::ReaChR(+ATR)* | Welch test | 0.02344 |
| 10F | *RIS::ReaChR(-ATR)* | *RIS::ReaChR(+ATR)* | Welch test | 0.00584 |
| 10G | *RIS::ReaChR(+ATR)* | *Wild type (+ATR)* | Logrank test | 0.09 |
| | | | Fisher's Exact test | 0.02 |
| | *RIS::ReaChR(-ATR)* | *Wild type (-ATR)* | Logrank test | 0.82 |
| 11C | Wild type control | Wild type stimulation | Welch test | 1.94557e-11 |
| | Wild type stimulation | *RIS::twk-18gf* stimulation | Welch test | 5.80697e-05 |
| 11D | Wild type stimulation | *RIS::twk-18gf* stimulation | Welch test | 0.01972 |
| 12A | wild type | *flp-11(-)* | Welch test with FDR correction for multiple testing | 4.013e-14 |
| | wild type | *RIS::unc-58gf(strong)* | Welch test with FDR | 4.0915e-15 |

| | | | | |
|---|---|---|---|---|
| | | | correction for multiple testing | |
| | wild type | *RIS::unc-58gf(strong), flp-11(-)* | Welch test with FDR correction for multiple testing | 3.6044e-14 |
| 12B | wild type | *flp-11(-)* | Fisher's Exact test with FDR correction for multiple testing | 1.05E-22 |
| | wild type | *RIS::unc-58gf(strong)* | Fisher's Exact test with FDR correction for multiple testing | 0.00016 |
| | wild type | *RIS::unc-58gf(strong), flp-11(-)* | Fisher's Exact test with FDR correction for multiple testing | 9.52E-14 |
| | *flp-11(-)* | *RIS::unc-58gf(strong), flp-11(-)* | Fisher's Exact test with FDR correction for multiple testing | 6.36E-05 |
| | *RIS::unc-58gf(strong)* | *RIS::unc-58gf(strong), flp-11(-)* | Fisher's Exact test with FDR correction for multiple testing | 5.33E-22 |
| S2A | wild type | *RIS::unc-58gf(strong)* | Fisher's Exact test | 7.26E-08 |
| | wild type | *RIS::unc-58gf(strong), aptf-1(-)* | Fisher's Exact test | 1.4798e-19 |
| | *RIS::unc-58gf(strong)* | *RIS::unc-58gf(strong), aptf-1(-)* | Fisher's Exact test | 3.5813e-16 |
| | *aptf-1(-)* | *RIS::unc-58gf(strong), aptf-1(-)* | Fisher's Exact test | 0.0026726 |
| S2B | wild type | *RIS::unc-58gf(strong)* | Welch test | 8.05065E-4 |
| | wild type | *aptf-1(-)* | Welch test | 2.60456E-13 |
| | wild type | *RIS::unc-58gf(strong), aptf-1(-)* | Welch test | 8.89446E-16 |
| | *aptf-1(-)* | *RIS::unc-58gf(strong), aptf-1(-)* | Welch test | 0.0767 |
| | *RIS::unc-58gf(strong)* | *RIS::unc-58gf(strong), aptf-1(-)* | Welch test | 0.00447 |

| | | | | |
|---|---|---|---|---|
| S3A | wild type | *RIS::twk-18* | Welch test with FDR correction for multiple testing | 2.9535e-08 |
| | wild type | *RIS::egl23(strong)* | Welch test with FDR correction for multiple testing | 3.1271e-13 |
| | wild type | *RIS::egl-23(weak)* | Welch test with FDR correction for multiple testing | 0.44566 |
| | wild type | *RIS::unc-58gf(weak)* | Welch test with FDR correction for multiple testing | 8.7484e-24 |
| | wild type | *RIS::unc-58gf(strong)* | Welch test with FDR correction for multiple testing | 3.4427e-20 |
| | *RIS::unc-58gf(weak)* | *RIS::unc-58gf(strong)* | Welch test with FDR correction for multiple testing | 0.96149 |
| S3B | wild type | *RIS::twk-18* | Welch test with FDR correction for multiple testing | 1.4404e-05 |
| | wild type | *RIS::egl23(strong)* | Welch test with FDR correction for multiple testing | 1.3078e-05 |
| | wild type | *RIS::egl-23(weak)* | Welch test with FDR correction for multiple testing | 0.98035 |
| | wild type | *RIS::unc-58gf(weak)* | Welch test with FDR correction for multiple testing | 0.80731 |
| | wild type | *RIS::unc-58gf(strong)* | Welch test with FDR correction | 0.0018333 |

| | | | | |
|---|---|---|---|---|
| | | | for multiple testing | |
| S4 | wild type | *RIS::unc-58gf(strong)* | Welch test | 0.44996 |
| S8 | wild type | *RIS::twk-18* | Welch test with FDR correction for multiple testing | 0.05085 |
| | wild type | *RIS::egl23(strong)* | Welch test with FDR correction for multiple testing | 0.10085 |
| | wild type | *RIS::egl-23(weak)* | Welch test with FDR correction for multiple testing | 0.41311 |
| | wild type | *RIS::unc-58gf(weak)* | Welch test with FDR correction for multiple testing | 0.16855 |
| | wild type | *RIS::unc-58gf(strong)* | Welch test with FDR correction for multiple testing | 0.79677 |
| S9B | wild type | *flp-11(-)* | Fisher's Exact test with FDR correction for multiple testing | 3.36E-24 |
| | wild type | *RIS::unc-58gf(strong)* | Fisher's Exact test with FDR correction for multiple testing | 0.081516 |
| | wild type | *RIS::unc-58gf(strong), flp-11(-)* | Fisher's Exact test with FDR correction for multiple testing | 6.83E-18 |
| | *flp-11(-)* | *RIS::unc-58gf(strong), flp-11(-)* | Fisher's Exact test with FDR correction for multiple testing | 0.081516 |
| | *RIS::unc-58gf(strong)* | *RIS::unc-58gf(strong), flp-11(-)* | Fisher's Exact test | 3.00E-13 |

| | | | | |
|---|---|---|---|---|
| | | | with FDR correction for multiple testing | |
| S10 | Wild type | *RIS::unc-58gf(strong)* | Welch test | 9.106e-13 |
| | Wild type | DAF-16(-) | Welch test | 4.73842e-08 |
| | Wild type | *RIS::unc-58gf(strong);* DAF-16(-) | Welch test | 1.28118e-12 |
| | *RIS::unc-58gf(strong)* | *RIS::unc-58gf(strong);* DAF-16(-) | Welch test | 0.31948 |
| | DAF-16(-) | *RIS::unc-58gf(strong);* DAF-16(-) | Welch test | 2.86134e-04 |

p-values and statistical tests for all experiments

**Movies S1-S8.**

**Movie S1-S6.**

The movies S1-6 show each a 3-hour recording of a sample worm for each RIS polarization strain from the experiment from Figures 5A-E, S7. The measured neuronal intensities and speeds of the worm are depicted in the right panel of the movie. Phases of neuronal inactivity and locomotion quiescence correlate. The time-lapse movies show 1 min of imaging in 1 sec.

**Movie S7**

The movie shows a growing plate of wild-type worms. The images were acquired at a framerate of 2Hz. The speed of the movie is 4fps.

**Movie S8**

The movie shows a growing plate of *RIS::unc-58gf* worms. The images were acquired at a framerate of 2Hz. The speed of the movie is 4fps.

### References


[1] S. R. Taylor *et al.*, "Molecular topography of an entire nervous system," *Cell*, vol. 184, no. 16, 2021, doi: 10.1016/j.cell.2021.06.023.

[2] J. Schwarz, J.-P. Spies, and H. Bringmann, "Reduced muscle contraction and a relaxed posture during sleep-like Lethargus," *Worm*, vol. 1, no. 1, 2012, doi: 10.4161/worm.19499.

[3] M. Turek, J. Besseling, J. P. Spies, S. König, and H. Bringmann, "Sleep-active neuron specification and sleep induction require FLP-11 neuropeptides to systemically induce sleep," *Elife*, vol. 5, no. MARCH2016, 2016, doi: 10.7554/eLife.12499.

[4] E. Maluck *et al.*, "A wake-active locomotion circuit depolarizes a sleep-active neuron to switch on sleep," *PLoS Biol.*, vol. 18, no. 2, 2020, doi: 10.1371/journal.pbio.3000361.

[5] Y. Wu, F. Masurat, J. Preis, and H. Bringmann, "Sleep Counteracts Aging Phenotypes to Survive Starvation-Induced Developmental Arrest in C. elegans," *Curr. Biol.*, vol. 28, no. 22, 2018, doi: 10.1016/j.cub.2018.10.009.

[6] A. Koutsoumparis *et al.*, "Sleep neuron depolarization promotes protective gene expression changes and FOXO activation.," *Curr. Biol.*, vol. 32, no. 10, pp. 2248-2262.e9, May 2022, doi: 10.1016/j.cub.2022.04.012.

[7] Brenner S. The genetics of Caenorhabditis elegans. Genetics. 1974;77(1):71-94. Epub 1974/05/01. PubMed PMID: 4366476.

[8] E. Yemini *et al.*, "NeuroPAL: A Multicolor Atlas for Whole-Brain Neuronal

Identification in C. elegans," *Cell*, vol. 184, no. 1, 2021, doi: 10.1016/j.cell.2020.12.012.
